# Rapid two-step target capture ensures efficient CRISPR-Cas9-guided genome editing

**DOI:** 10.1101/2024.10.01.616117

**Authors:** Honglue Shi, Noor Al-Sayyad, Kevin M. Wasko, Marena I. Trinidad, Erin E. Doherty, Kamakshi Vohra, Ron S. Boger, David Colognori, Joshua C. Cofsky, Petr Skopintsev, Zev Bryant, Jennifer A. Doudna

## Abstract

RNA-guided CRISPR-Cas enzymes initiate programmable genome editing by recognizing a 20-base-pair DNA sequence adjacent to a short protospacer-adjacent motif (PAM). To uncover the molecular determinants of high-efficiency editing, we conducted biochemical, biophysical and cell-based assays on *S. pyogenes* Cas9 (*Spy*Cas9) variants with wide-ranging genome editing efficiencies that differ in PAM binding specificity. Our results show that reduced PAM specificity causes persistent non-selective DNA binding and recurrent failures to engage the target sequence through stable guide RNA hybridization, leading to reduced genome editing efficiency in cells. These findings reveal a fundamental trade-off between broad PAM recognition and genome editing effectiveness. We propose that high-efficiency RNA-guided genome editing relies on an optimized two-step target capture process, where selective but low-affinity PAM binding precedes rapid DNA unwinding. This model provides a foundation for engineering more effective CRISPR-Cas and related RNA-guided genome editors.

## INTRODUCTION

The widespread utility of CRISPR-Cas9 for programmable genome editing in human cells, plants and other eukaryotes has propelled interest in understanding the determinants of efficient genome modification^1,2^. Cas9 recognizes target sequences within genomes using an ATP-independent process in which Cas9-bound guide RNAs base pair with a 20-nucleotide sequence within double-stranded DNA. This mechanism begins with Cas9 binding to a 2-4 base pair (bp) protospacer adjacent motif (PAM), triggering DNA unwinding and RNA-DNA hybridization to form an *R*-loop^3–6^ (Fig. 1A). While stable target capture requires RNA-DNA hybridization at the seed region (5-10 bps), efficient DNA cleavage necessitates complete *R*-loop formation with full RNA-DNA complementarity^3,4,7–10^ (Fig. 1A). The process of Cas9-mediated *R*-loop formation has been well-characterized^4,7–9,11,12^, but the mechanism by which RNA-guided enzymes avoid entrapment by the vast excess of non-specific sequences in the genome remains unclear^13,14^.

**Fig. 1.**
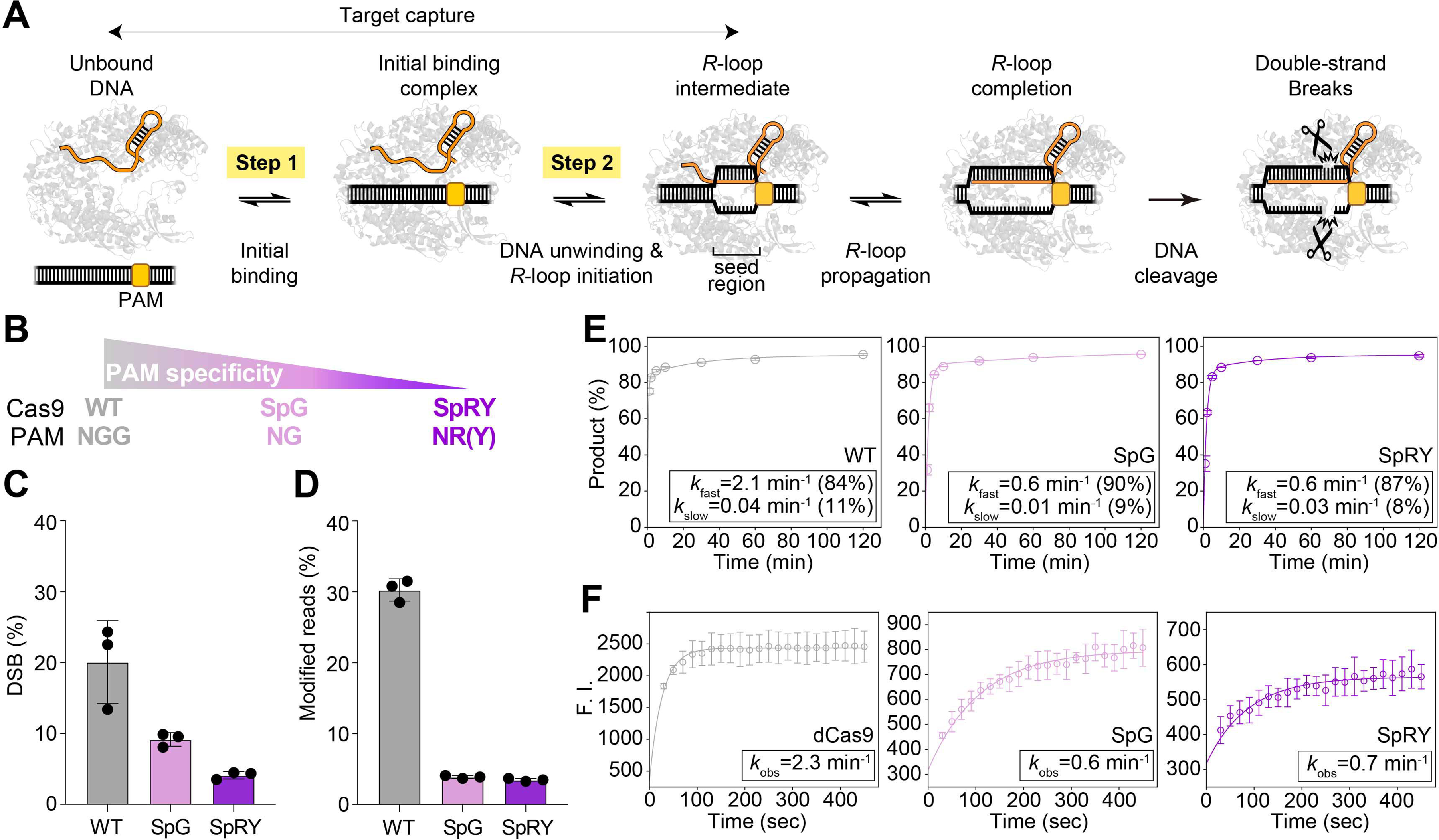
PAM-relaxed Cas9 variants display reduced genome editing efficiencies and bulk DNA cleavage kinetics on a canonical NGG PAM. **(A)** A model of RNA-guided DNA targeting by Cas9. The two-step target capture process that differentiates PAM-relaxed Cas9 variants from WT *Spy*Cas9 is highlighted. **(B)** The three *Spy*Cas9 proteins used in this study with their respective PAM recognition: WT *Spy*Cas9 (gray), SpG (pink) and SpRY (purple). **(C)** Quantifications of DNA double-stranded breaks (DSB) by droplet digital PCR (ddPCR) on the target site at 8 h post plasmid transfection (n=3 replicates). See also Fig. S1C for more data. **(D)** Quantifications of indels on the target site at 72 h post RNP nucleofection (n=3). See also Fig. S1E for more data. **(E)** Time-course analysis of average DNA cleavage products (n=3) for sgRNA2 in 5 mM Mg^2+^ at 24°C for TS (see Methods). The average rate constants, with the amplitudes from the observed double-exponential decay, are provided in figure legends. See also Fig. S2A for NTS cleavage and Fig. S2C-I for other sgRNAs and conditions. **(F)** Time-course analysis of the average fluorescence signal (n=3) in the 2AP assay for the same sequence in similar conditions as **(E)** (see Methods). For WT *Spy*Cas9, catalytically dead Cas9 (dCas9) is used here. The rate constants of the observed mono-exponential decay are provided in figure legends. See also Fig. S2B for the 2AP-labeling position (15th nt from PAM) and intercept determination.

Although *R*-loop formation and DNA cleavage occur within seconds to minutes upon target DNA recognition *in vitro*^12,15^ and in cells^16^, genome edits take hours to days, indicating that other steps are rate-limiting. For *S. pyogenes* Cas9 (*Spy*Cas9), the abundance of its NGG PAM in the genome (roughly every 16 dinucleotides) necessitates repeated PAM binding and release until the target is located^13,14^. Furthermore, the PAM requirement for target capture restricts *Spy*Cas9 to modifying sequences next to an NGG motif^3,4^. To expand target access, efforts have focused on reducing or eliminating PAM specificity^17–20^, yielding PAM-relaxed *Spy*Cas9 variants including SpG and SpRY^19^ that differ by only a few point mutations in the PAM-interacting domain yet possess “PAM-less” target recognition. These variants provide a basis for investigating the role of PAM specificity and affinity in *Spy*Cas9’s target capture, cleavage and genome editing efficiencies.

Using cell-based assays as well as biochemical and single-molecule experiments, we show that compared to wild-type (WT) *Spy*Cas9, the PAM-relaxed Cas9 variants SpG and SpRY are slower to identify target sequences due to non-specific DNA interactions. Even after arriving at the target, these variants unwind DNA more slowly because they become kinetically trapped in a stable initial binding complex and are less proficient at initiating *R*-loops. Efforts to accelerate unwinding and stabilize the *R*-loop at the target site only partially mitigate these effects, as off-target binding continues to limit overall efficiencies. Our findings support a model in which increased PAM specificity with limited DNA affinity can enhance genome editing efficiency. These results highlight the crucial trade-off between PAM flexibility and editing efficiency and show that the effectiveness of *Spy*Cas9 for genome editing arises from its intrinsic PAM recognition properties.

## RESULTS

### Reduced editing efficiencies arise from enzymatic limitations before R-loop completion

PAM-relaxed variants of *Spy*Cas9, SpG (PAM: NG) and SpRY (PAM: NR; R=A,G) (Fig. 1B), are less efficient genome editing enzymes relative to WT *Spy*Cas9 (PAM: NGG)^19,21^. We used these activity differences as a basis for exploring the mechanism by which PAM recognition influences enzyme-mediated editing. To quantify genome editing outcomes, we expressed WT, SpG and SpRY Cas9 separately in HEK293T cells along with a single guide RNA (sgRNA) targeting an NGG-proximal sequence in the gene *EMX1* (sgRNA1). Despite comparable expression levels of Cas9 and sgRNA at early time points after plasmid transfection in each case (Fig. S1A, B), SpG and SpRY were 2-4 fold slower than WT *Spy*Cas9 at both DNA cutting (Fig. 1C; Fig. S1C) and inducing genome edits (Fig. S1D). Furthermore, when defined amounts (10 pmol) of pre-assembled Cas9 ribonucleoproteins (RNPs) with similar active enzyme fractions (Fig. S1G) were nucleofected into HEK293T cells, WT *Spy*Cas9 induced ∼30% indels with sgRNA1 within 72 h, while SpG and SpRY reached only 4% and 3%, respectively, at 72 h (Fig. 1D). This inefficiency becomes even more pronounced with different sgRNAs and higher RNP doses (Fig. S1E), consistent with guide-dependent effects reported in previous studies^19,21,22^. These results suggest that reduced efficiencies of PAM-relaxed variants result from their intrinsic enzymatic properties.

To determine the molecular basis of the reduced efficiencies of SpG and SpRY, we focused on key steps in Cas9’s targeting mechanism: PAM binding, *R*-loop formation and DNA cleavage (Fig. 1A). We first examined whether differences in *R*-loop formation or DNA cleavage kinetics accounted for the observed editing efficiency differences. Bulk DNA cleavage assays using dual-fluorescently labeled double-stranded DNA (dsDNA) substrates revealed two phases of DNA cleavage: a dominant fast phase governed by *k*_fast_, and a slower phase governed by *k*_slow_ (Fig. 1E), consistent with prior studies^15,23^. WT *Spy*Cas9 completed the fast phase of target strand (TS) cutting within one minute, while SpG and SpRY showed ∼3.5-fold slower *k*_fast_ (Fig. 1E), with similar results for non-target strand (NTS) cleavage (Fig. S2A). The reduced cleavage rate was guide-dependent, as previously reported^19,21,22^, and occurred under reduced Mg^2+^ concentration (0.2 mM) conditions that reflect intracellular Mg^2+^ levels in mammalian cells^24,25^ (Fig. S2C-I). Using 2-aminopurine (2AP) labeled dsDNA to track Cas9-induced DNA unwinding^26,27^, we found that the apparent rate constants of *R*-loop formation (*k*_obs_) for SpG and SpRY were similarly ∼3.5-fold slower, matching the *k*_fast_ in the DNA cleavage assay (Fig. 1F; Fig. S2B). These results are consistent with the kinetic delays for SpG and SpRY occurring before *R*-loop completion, rather than from differences in DNA cleavage chemistry^28^.

### SpRY is inefficient at forming an initial stable R-loop intermediate

Bulk biochemical assays suggested that SpG and SpRY’s inefficiency arises before *R*-loop completion. To determine which substeps in *R*-loop formation are affected, we employed Gold Rotor Bead Tracking (AuRBT)^29^ to detect DNA structural transitions upon Cas9 binding at base-pair resolution. We immobilized a DNA tether, containing an NGG site next to a target sequence, and measured real-time changes in DNA twist (Δθ) during *R*-loop formation and collapse (Fig. 2A). In these experiments we focused on SpRY due to its most pronounced differences in kinetic behavior relative to WT *Spy*Cas9. To prevent DNA cleavage during data collection, we introduced D10A and H840A mutations that create catalytically dead versions of WT *Spy*Cas9 (dCas9) and SpRY (dSpRY) while retaining *R*-loop formation ability^4^. After introducing 0.8 nM dCas9 or 4 nM dSpRY RNP into the channel, we observed stepwise changes in Δθ corresponding to transitions between *R*-loop states with differing numbers of bp unwound (Fig. 2B, C). Unlike previous work that focused on the effects of supercoiling on *R*-loop dynamics^12^, here we monitored equilibrium fluctuations between states on a relaxed DNA tether.

**Fig. 2.**
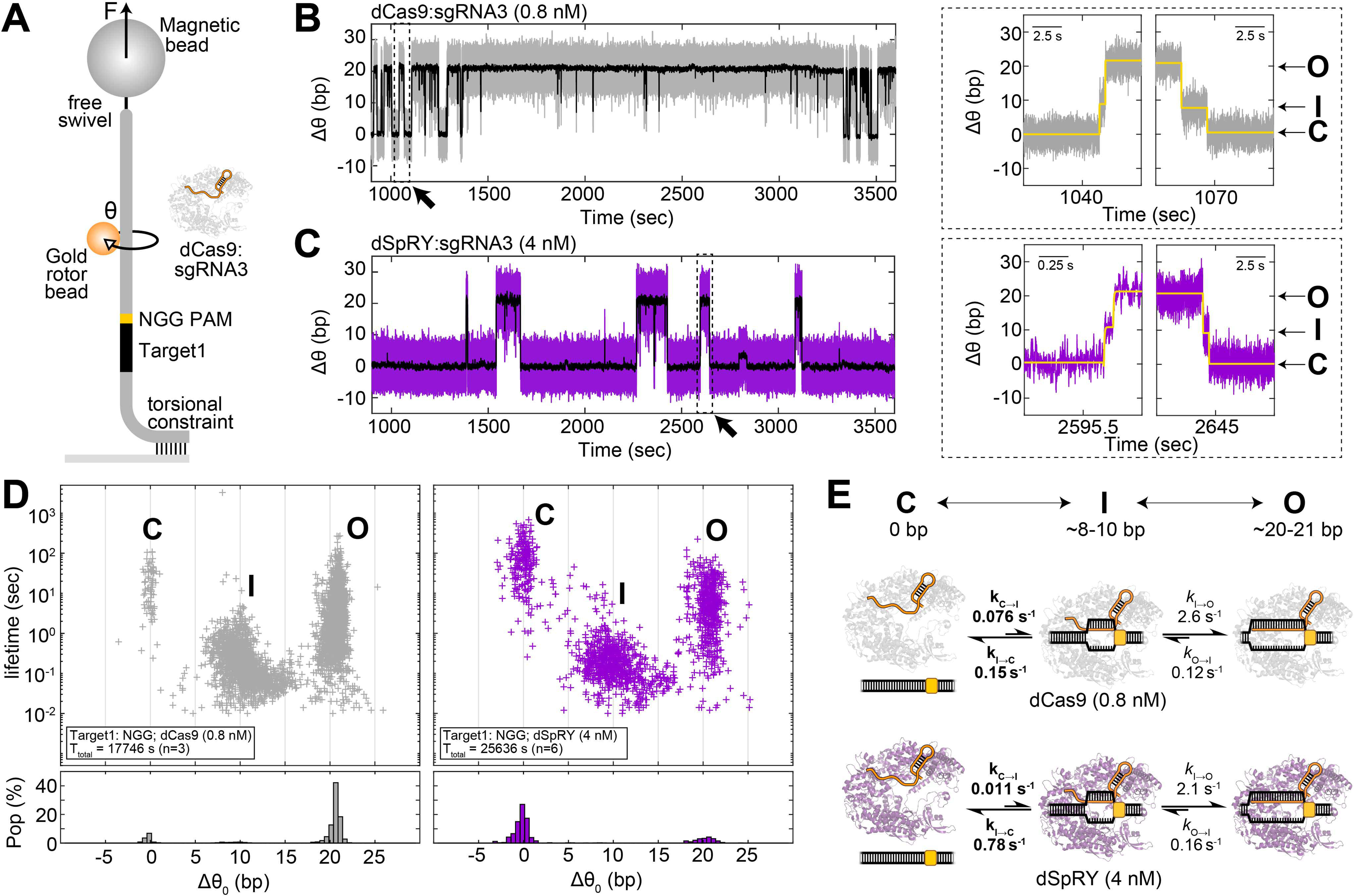
SpRY disfavors the formation of the *R*-loop intermediate state in AuRBT. **(A)** Schematic overview of the AuRBT experimental setup. dCas9 and dSpRY, both with similar active protein fractions (Fig. S1H), were used at concentrations of 0.8 nM and 4 nM, respectively. sgRNA3 contains a 20-bp match to the Target1 sequence flanking an NGG site. θ is the measured rotor bead angle. All AuRBT experiments were performed under F = 7 pN of tension. **(B-C)** Example traces from time-resolved measurements of DNA unwinding (Δθ) for **(B)** 0.8 nM dCas9 (gray) and **(C)** 4 nM dSpRY (purple). 250-ms averaged traces are shown in black. Diagonal arrows highlight zoomed-in regions of *R*-loop formation and collapse events through a discrete intermediate (right panels). Yellow lines represent idealized traces generated by the Steppi change-point analysis. **(D)** (Top) Scatter plots illustrating the unwinding lifetime and change in equilibrium twist (Δθ0) for merged Steppi-scored states (see Methods) across all binding events for 0.8 nM dCas9 (gray, left) and 4 nM dSpRY (purple, right), highlighting a closed DNA state “C”, an intermediate state “I” and an open DNA state “O”. The total collection time (Ttotal) and number of replicates (n) are provided in the legend. (Bottom) The lifetime weighted population at each corresponding Δθ0 with bin size of 0.5 bp (see Methods). **(E)** Rate constants of transitions between C and I and between I and O for dCas9 (gray, upper) and dSpRY (purple, lower) showing the primary distinction between dCas9 and dSpRY lies in the rates associated with the transition between C and I (bold). The average unwinding (bp) in each state is provided.

We identified *R*-loop states and characterized their kinetics using automated change-point detection (“Steppi”)^30^, scoring transitions between three different levels of DNA unwinding: a closed state (C) with no unwinding, an intermediate state (I) with ∼8-10 bp unwound, and an open state (O) with ∼20-21 bp unwound (Fig. 2D). This state structure was consistent between dSpRY and dCas9, and with previous measurements of dCas9 obtained using non-equilibrium twist ramping^12^. Complete *R*-loop formation and collapse proceeded through the transient I state, and we analyzed C ↔ I ↔ O transitions to obtain the corresponding rates *k*_C→I_, *k*_I→C_, *k*_I→O_, and *k*_O→I_. Compared to dCas9, dSpRY had a 7-fold slower *k*_C→I_ (despite being 5-fold higher in concentration), and a 5-fold faster *k*_I→C_ (Fig. 2E), indicating dSpRY is far less likely to form the *R*-loop intermediate, instead favoring the closed state. The subsequent *R*-loop propagation (I ↔ O) kinetics were similar between dSpRY and dCas9 (Fig. 2E). These results show that differences in *R*-loop formation between the two enzymes are confined to the earliest steps involved in unwinding the bps adjacent to the PAM site, leading to slow and unfavorable *R*-loop completion that could explain the observed reduction in SpRY’s DNA cleavage rate in bulk assays.

### SpRY is kinetically trapped in a low-energy initial binding complex

To determine why SpRY is less efficient at melting DNA bps to enable guide RNA strand invasion and form the *R*-loop intermediate, we used AuRBT to dissect the target capture process (Fig. 1A). Because the C ↔ I transition involves both binding of the RNP to the target and DNA unwinding, *k*_C→I_ is expected to depend on the RNP concentration ([RNP]). For dCas9, *k*_C→I_ increases linearly with rising [RNP], as expected for weak PAM binding^6,23,28,31^ that is far from saturation within our experimental range. At low concentrations *k*_C→I_ also increases with [RNP] for SpRY, but with characteristically lower rates, reflecting the slow formation of the seed intermediate. Furthermore, *k*_C→I_ reaches a plateau at moderate [RNP], suggesting a rate-limiting step after initial binding for dSpRY (Fig. 3A). The remaining kinetic rates, which involve transitions between bound states, are approximately independent of RNP concentration as expected, with closely equivalent kinetics between the two enzymes for the late I ↔ O step and faster collapse of the intermediate *k*_I→C_ for dSpRY (Fig. 3B, S3E).

**Fig. 3.**
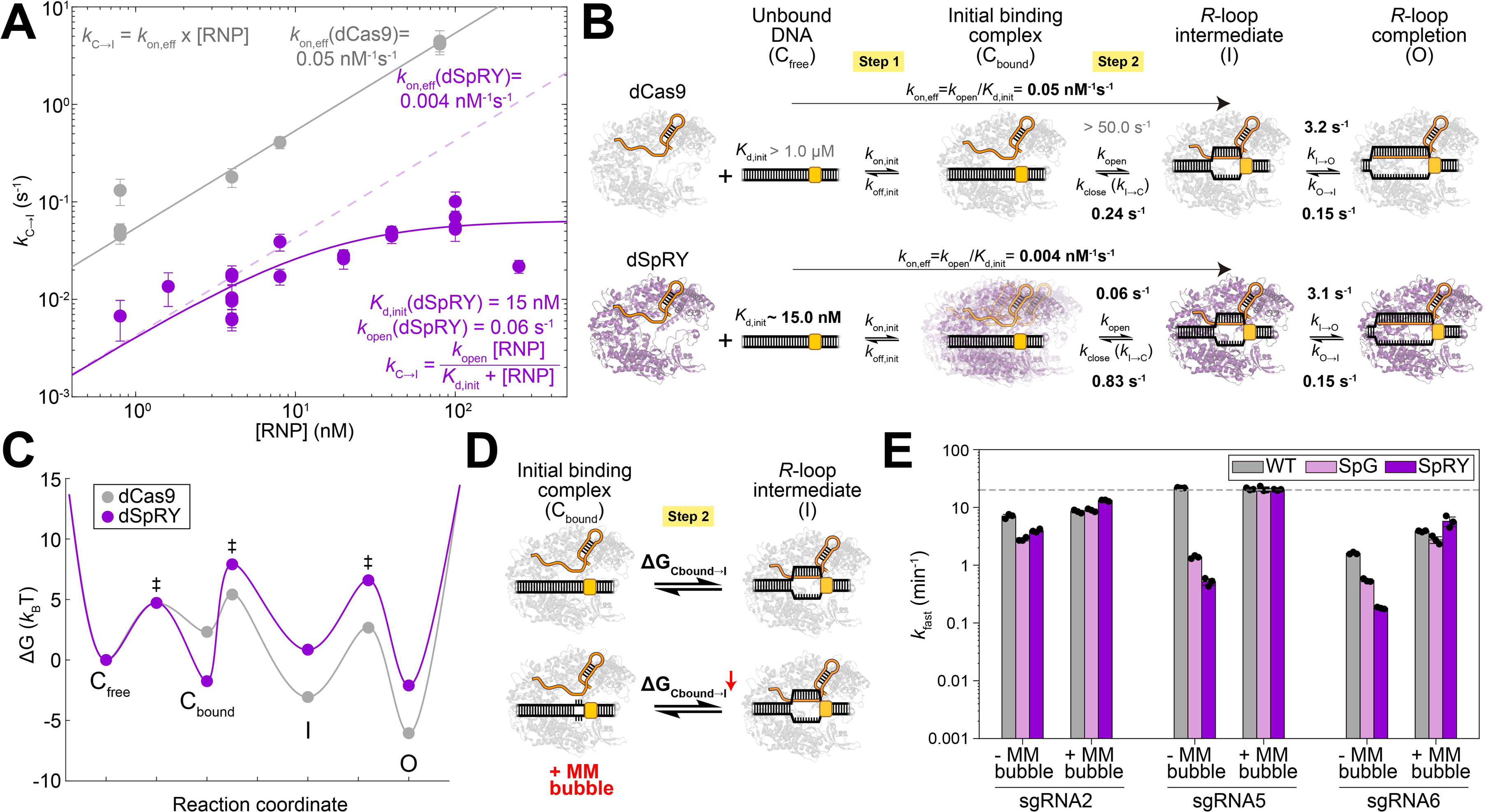
SpRY is trapped in a low-energy initial binding complex. **(A)** *k*C→I as a function of [RNP] for dCas9 (gray) and dSpRY (purple) using the same DNA and RNA sequences in Fig. 2A. Errorbars were calculated assuming Poisson statistics. Solid lines are fit of *k*C→I vs [RNP] (RNP concentration), assuming a linear model with *k*on,eff(dCas9) for dCas9 and a hyperbolic saturation model with initial affinity *K*d,init(dSpRY) and maximal unwinding rate *k*open(dSpRY) for dSpRY. The equations for each model are provided as insets. The faded dashed line has a slope *k*on,eff(dSpRY) = *k*open(dSpRY)/*K*d,init(dSpRY) and shows the linear dependence of *k*C→I on dSpRY [RNP] in the low [RNP] regime. See also Note S1 for model selection, Table S1 for rate constants, Table S2 for fit details, Fig. S3A-C for example raw traces, and Fig. S3E-G for other rate constants (*k*I→C, *k*I→O and *k*O→I). **(B)** A model illustrating distinct kinetics in the two steps of target capture between dCas9 and dSpRY. Rate constants in bold are derived from our AuRBT measurements, while those in gray are inferred from model constraints or prior measurements^6,12,23,28,31^. See also Note S1 for model details. **(C)** The free energy landscape of the kinetic model including Cfree, Cbound, I and O, assuming *K*d,init(dCas9) = 1 μM for dCas9, *k*on,init(dCas9) = *k*on,init(dSpRY) = 0.1 nM^-1^s^-1^, and [RNP] = 100 nM. The “‡” symbol indicates a transition state. See also Table S3 and Note S1 for the chosen values to build the free energy diagram. **(D)** Schematic showing how a mismatch (MM) bubble next to an NGG PAM reduces the energetic cost of DNA unwinding. **(E)** Bar graph depicting the average fast-phase rate constant (*k*fast) for DNA cleavage with and without an MM bubble adjacent to an NGG PAM. The gray dashed line represents the detection limit of the assay. See also Fig. S2D, G, I and Fig. S3H for time-course data.

Based on these measurements, we developed a kinetic model (Fig. 3B; Note S1) that describes Cas9’s target capture as two key steps: initial binding (C_free_ ↔ C_bound_) and subsequent DNA unwinding/*R*-loop intermediate formation (C_bound_ ↔ I). Here, the directly observed closed DNA conformation (state C) is kinetically separated into a free state C_free_ where the target site is unoccupied, and a bound state C_bound_ where an enzyme at the target site is poised to initiate *R*-loop formation. In this model, dCas9 shows weaker initial binding (*K*_d,init_ ∼ 1-10 μM)^6,23,28,31^ but rapidly transitions to form the *R*-loop intermediate (*k*_open_ > 50 s^-1^). In contrast, dSpRY has much stronger initial binding (*K*_d,init_ ∼ 15 nM) but transitions to the intermediate over 800 times slower than dCas9 (*k*_open_ ∼ 0.06 s^-1^), explaining the slower and hyperbolic kinetics of dSpRY. This kinetic model can be depicted as a free energy landscape (Fig. 3C; Note S1) that shows dSpRY initially occupying a low-energy binding state, and then encountering a high energy barrier towards subsequent DNA unwinding. Importantly, this low-energy state may represent an ensemble of binding modes by dSpRY rather than a single defined conformation. Based on this energy landscape, the primary cause for dSpRY’s inefficiency at forming an *R*-loop is its tendency to become kinetically trapped in this initial binding complex. In addition, the *R*-loop intermediate state in dSpRY is thermodynamically disfavored (Fig. 3C), likely due to weaker stabilizing interactions in the seed region^28^. The combined effects of the kinetic trap and unstable *R*-loop intermediate result in a ΔG_Cbound→I_ that is approximately 8 *k*_B_T higher than that of dCas9.

Based on this model, we predict that reducing ΔG_Cbound→I_ could enhance SpRY’s reduced cleavage rate in bulk assays. To test this, we introduced a small mismatch (MM) bubble next to the NGG PAM, aiming to lower the energy barrier for the C_bound_ ↔ I transition by destabilizing the target duplex next to the NGG^32,33^ (Fig. 3D). In DNA cleavage assays, this MM bubble had little to no impact on WT *Spy*Cas9’s cleavage rate but significantly improved SpG and SpRY’s rates, rendering their bulk cleavage activities to match WT *Spy*Cas9 (Fig. 3E; Fig. S2D, G, I; Fig. S3H). This result supports our hypothesis that difficulties in unwinding the initial seed region contribute to the reduced efficiencies of SpG and SpRY, and suggests that lowering the energy barrier towards the *R*-loop intermediate can overcome these limitations.

### Kinetic traps induce strong and promiscuous DNA binding

Given that Cas9 must navigate through many non-target sequences during genome surveillance, we asked if SpRY’s kinetic trapping also exacerbates non-specific binding. To address this, we investigated how SpRY interrogates non-specific sequences compared to WT *Spy*Cas9.

Using AuRBT, we monitored the dynamics of target capture by introducing 50 nM RNP with a sgRNA containing only a 3-bp match next to an NGG PAM. For WT *Spy*Cas9, we observed short-lived, small-magnitude spikes in Δθ, indicating transient DNA unwinding events (Fig. 4A, B). These states were consistently recorded across various sgRNA-DNA combinations (Fig. S4A) but disappeared when NGG was replaced with NCG (Fig. 4C, D; Fig. S4B) or when apo Cas9 was used (Fig. S4C). The amount of unwinding (∼2 bps) was also consistent with predictions from cryo-EM structures (Fig. S4D) (PDB: 7S38)^6^, and the unwinding lifetime aligns with dwell times seen in single-molecule FRET studies of DNA binding (Fig. S4E)^34^. In sharp contrast, SpRY exhibited a prolonged Δθ baseline shift under the same conditions (Fig. 4E). This prolonged shift was [RNP]-dependent (Fig. 4F; Fig. S4F, G), independent of the NGG PAM (Fig. S4H), and saturated at moderate [RNP]. These observations suggest SpRY strongly interacts with non-specific sites, leading to a slight unwinding signal.

**Fig. 4.**
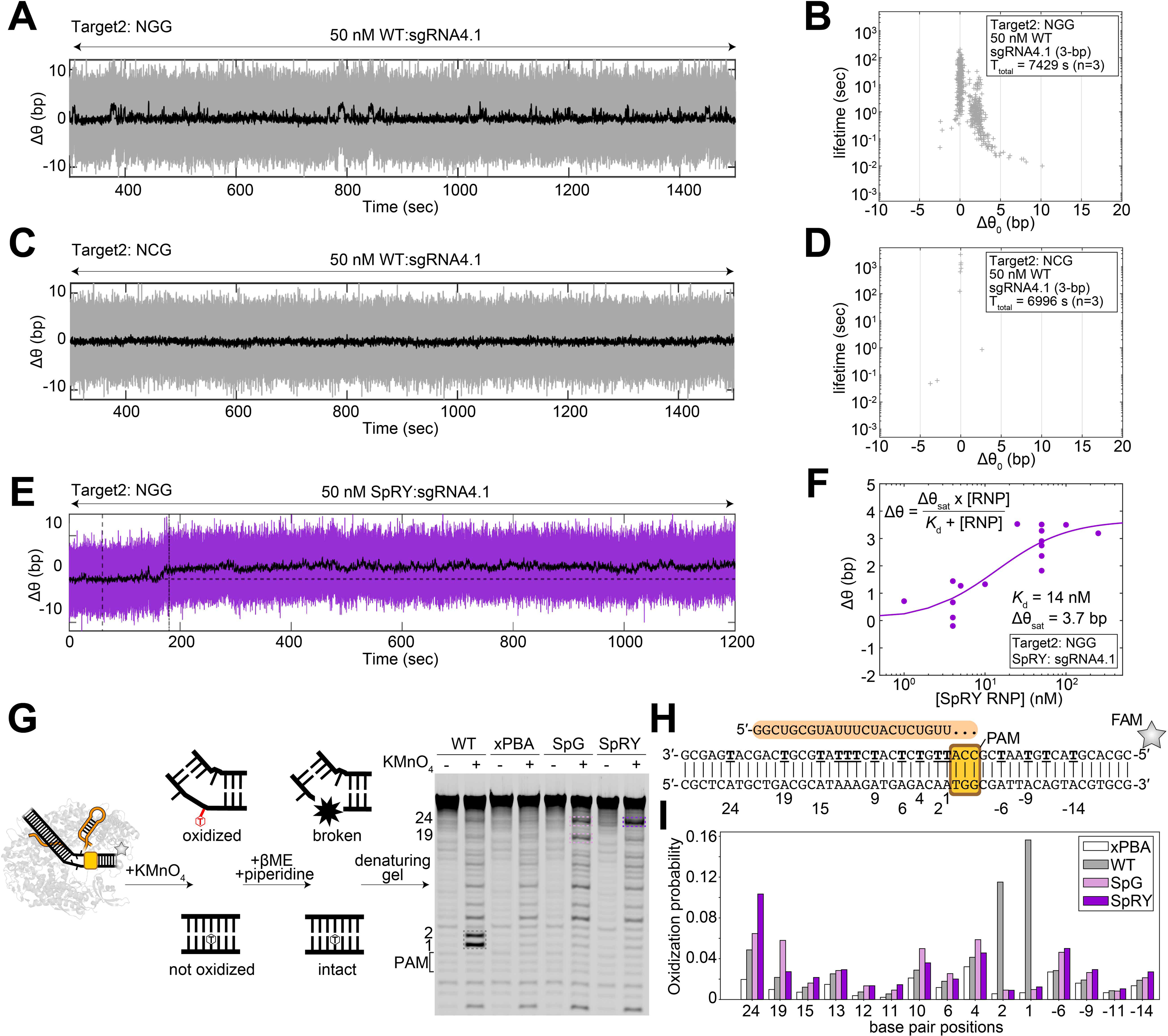
Promiscuous DNA binding by PAM-relaxed Cas9 variants. **(A, C, E)** Example traces of time-resolved measurements of equilibrium twist change Δθ over time for **(A and C)** WT *Spy*Cas9 (50 nM) and **(E)** SpRY (50 nM), each paired with sgRNA4.1, containing a 3-bp match to the Target2 sequence flanking **(A and E)** an NGG site or **(C)** an NCG site on the DNA tether. 250-ms averaged traces are shown in black. For **(E)**, the vertical dashed lines (--) and dash-dot lines (-.) indicate the start and end, respectively, of the flow of SpRY RNP into the chamber. **(B, D)** Scatter plots of unwinding lifetime and Δθ0 for merged Steppi-scored states corresponding to experiments described in **(A, C)**. The total collection time (Ttotal) and number of replicates (n) are provided in the legend. See also Fig. S4A-C for more data with different sequences and conditions. **(F)** Fit of average Δθ baseline shift and [RNP] for SpRY to a binding equation yielded an apparent *K*d ∼ 14 nM for sgRNA4.1 on the Target2 sequence flanking an NGG site. See also Fig. S4F-H for raw traces. **(G)** Schematic overview of the permanganate DNA footprinting assay. The UREA-PAGE gel image used to generate the graph in **(I)** highlights KMnO4-dependent enriched bands for WT *Spy*Cas9 (gray), SpG (pink) and SpRY (purple) in colored dashed boxes. **(H)** Sequence of the sgRNA (sgRNA3) and the dsDNA substrate with all the reactive thymines on the TS underlined. **(I)** Oxidation probabilities of thymines across the dsDNA substrate.

Fitting this [RNP]-dependent Δθ baseline shift for SpRY yielded an effective *K*_d,eff_ of ∼14 nM (Fig. 4F). We obtained a similar result (*K*_d,eff_ ∼ 16 nM) by fitting an underlying baseline shift observed in the previous dSpRY experiments using full-match sgRNA (see Methods, Fig. S3D), indicating consistent non-specific binding behavior across conditions. The agreement between these measurements and dSpRY’s affinity for the initial binding complex (*K*_d,init_ ∼ 15 nM, from Fig. 3A) hints that a common non-specific binding mode may be responsible for both observations: the kinetic trap hindering target DNA capture also leads to prolonged, unproductive off-target interactions.

To visualize the locations of DNA binding and unwinding between these Cas9 variants in the absence of sgRNA sequence complementarity but in the presence of a canonical NGG PAM, we used a permanganate footprinting assay. Consistent with previous findings^6^, WT *Spy*Cas9 induced bp melting adjacent to the NGG site, while the xPBA variant, lacking PAM-binding arginines, showed minimal signal (Fig. 4G-I). In contrast, SpG and SpRY did not induce detectable bp melting near the NGG site but caused promiscuous thymine exposure at various locations (Fig. 4G-I). Together, these results suggest that PAM-relaxed Cas9 variants tend to strongly and promiscuously bind DNA.

### Relaxed PAM specificity increases non-specific DNA interference

Based on these observations of SpRY’s promiscuous and strong non-specific binding, we hypothesized that the DNA cleavage efficiency of PAM-relaxed Cas9 variants would be more significantly inhibited when exposed to excess non-specific DNA. We tested this by examining the DNA cleavage kinetics of these Cas9 variants in the presence of excess non-specific DNA (Fig. 5A), mimicking genomic conditions in cells. We repeated the DNA cleavage assay from Fig. 1E at 37°C with 100 nM RNP (sgRNA2) and 10 nM fluorescent target DNA substrate in the presence of ∼8 nM (0.8×) 2.2-kb supercoiled DNA plasmids lacking complementarity to sgRNA2 as non-specific competitor. The plasmid competitor did not affect WT *Spy*Cas9’s cleavage rate, but the *k*_fast_ for SpG and SpRY was approximately ∼5-fold and ∼15-fold slower, respectively, compared to conditions without competitor (Fig. 5B, C; Fig. S2D; Fig. S5A). In addition, SpRY exhibited a larger slow phase (30%) (Fig. 5B, C), suggesting that non-specific competitor interactions could contribute to the slow phase of DNA cleavage. The slower DNA cleavage by SpG and SpRY was not due to plasmid supercoiling, as similar inhibition was observed with a linear PCR product with identical DNA sequences (Fig. S5B). Electrophoretic Mobility Shift Assay (EMSA) further confirmed SpRY’s higher affinity for the linearized plasmid competitor DNA, as 0.5 μM SpRY RNP caused a band shift, while WT *Spy*Cas9 showed no binding even at 16 μM (Fig. S6A-B).

**Fig. 5.**
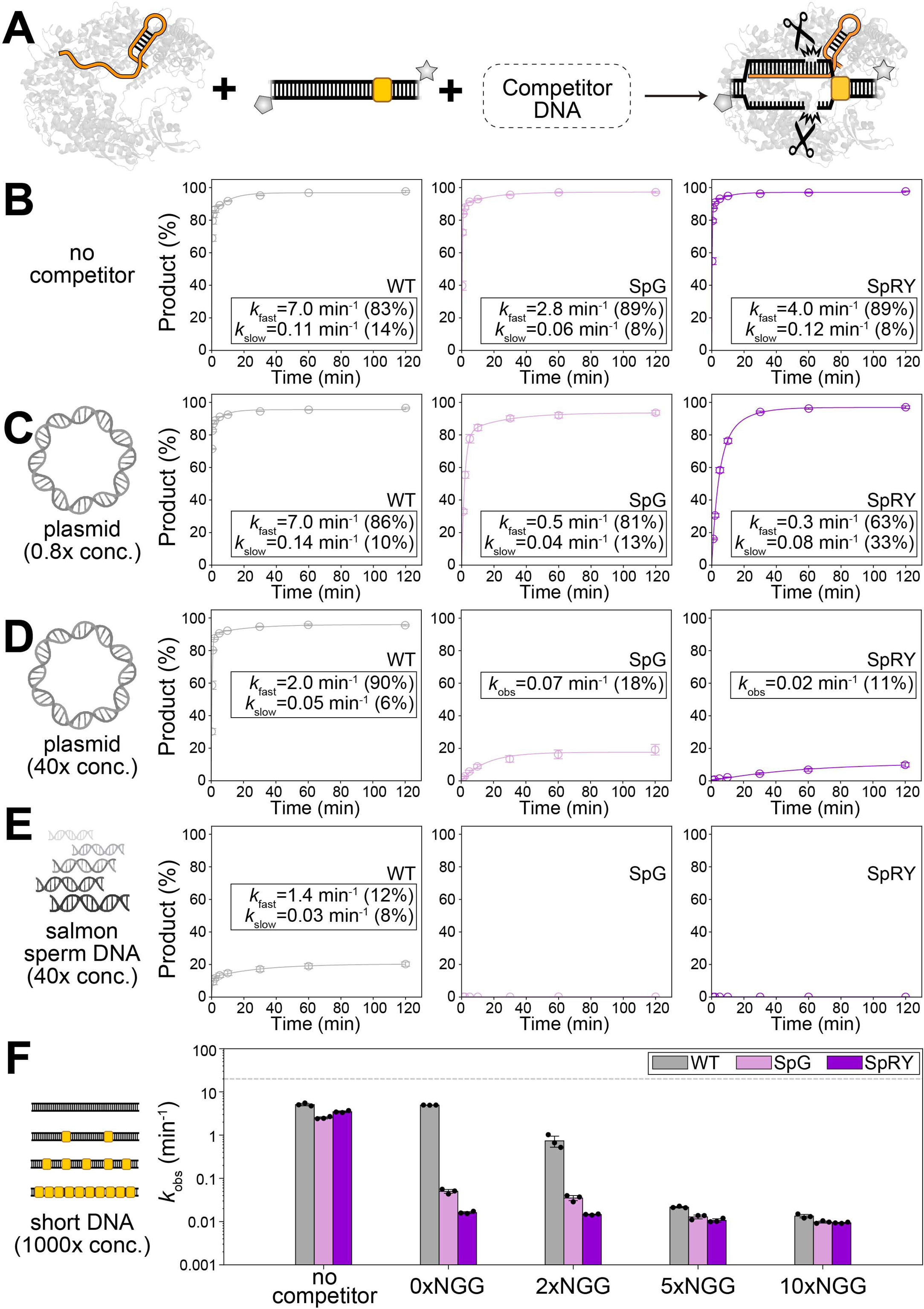
Inhibition of PAM-relaxed Cas9 variants cleavage activity by DNA competitors. **(A)** Schematic overview of bulk DNA cleavage assay conducted in the presence of various DNA competitors. **(B-E)** Time-course analysis of average DNA cleavage products (n=3) for sgRNA2 at 10 mM Mg^2+^ and 37°C under different conditions: **(B)** without competitors **(C)** with a 0.8× 2.2-kb plasmid competitor **(D)** with a 40× 2.2-kb plasmid competitor, and **(E)** with a 40× salmon sperm DNA competitor. The average rate constants (*k*obs for a mono-exponential decay model; *k*fast, *k*slow for a double-exponential decay model) along with their amplitudes are provided in figure legends (n=3). See also Fig. S2D and Fig. S5A, C, D for NTS cleavage. **(F)** Bar graph depicting the average rate constant (*k*obs) of the mono-exponential decay for DNA cleavage in the presence of a short dsDNA competitor containing various numbers of 3-bp seed sequences flanking an NGG site (n=3). The gray dashed line represents the detection limit of the assay. See also Fig. S5G for time-course data. All errorbars represent standard deviations.

To further challenge the DNA cleavage kinetics, we increased the concentration of the plasmid competitor to 400 nM (40×), resulting in significantly greater inhibition. Under this condition, WT *Spy*Cas9 showed a minimal 3.6-fold reduction in the *k*_fast_ (Fig. 5D; Fig. S5C). However, with the data adequately fit to a mono-exponential model, the *k*_obs_ for SpG was ∼40-fold slower, and for SpRY, it was ∼200-fold slower than the predominant *k*_fast_ without competitor (Fig. 5D; Fig. S5C). We then replaced the plasmid competitor with purified salmon sperm DNA (∼200-500 bps), resulting in both increased sequence diversity and possibly higher concentration of heterogeneous exposed ends, increasing non-specific binding sites. Repeating the cleavage assay at the same mass concentration (40×) showed that the *k*_fast_ for WT *Spy*Cas9 was ∼5-fold slower, and the product amplitude was reduced to ∼20% (Fig. 5E; Fig. S5D). For SpG and SpRY, no detectable DNA cleavage activity was observed (Fig. 5E; Fig. S5D). Substantial inhibition of SpG and SpRY was observed across different sgRNAs and different competitor concentrations (Fig. S5E), and at 0.2 mM Mg^2+^ (Fig. S5F).

We then tested whether DNA cleavage inhibition correlated with the number of NGG-dependent off-target sites. Using a 1000-fold molar excess of 60-mer dsDNA with zero, two, five or ten off-target sites (3-bp match of sgRNA2 next to NGG), we found that the cleavage rate reductions correlated with the number of NGG-dependent off-target sites for WT *Spy*Cas9 (Fig. 5F; Fig. S5G). In contrast, this correlation was less evident for SpG and minimal for SpRY (Fig. 5F; Fig. S5G), indicating that SpG and SpRY experience more generalized inhibition by short dsDNA competitors, irrespective of NGG presence. Taken together, these results suggest that increased PAM promiscuity in SpG and SpRY leads to widespread off-target interactions that prolong target identification and subsequent cleavage, correlating with their reduced editing efficiencies in cells.

### Facilitating on-target unwinding is insufficient to overcome off-target interactions

The competition experiments demonstrated that even with rapid unwinding kinetics like those of WT *Spy*Cas9, the presence of numerous PAM-containing off-target sites can still hinder cleavage efficiency. Given that SpG and SpRY possess broader PAM recognition and exhibit promiscuous DNA binding, we hypothesized that their inefficiencies could not be fully mitigated by simply accelerating on-target unwinding. Previously, by introducing a MM bubble next to an NGG PAM to lower the energy barrier for DNA unwinding (ΔG_Cbound→I_), we restored SpG and SpRY’s on-target activity to WT *Spy*Cas9 levels in bulk assays (Fig. 3D, E). Here, we repeated the competition assays using a target with an MM bubble and a 40× plasmid competitor. While WT SpyCas9 showed a modest cleavage rate increase (∼1.7-fold), SpG and SpRY exhibited significant improvements in *k*_obs_ (∼5-fold), and the total product amplitude increased from ∼15% to 80% with a MM bubble (Fig. 6A; Fig. S6C). However, despite these enhancements, SpRY’s rate remained 30-fold slower than WT under competitor conditions, indicating persistent off-target binding still hampers its efficiency as expected.

**Fig. 6.**
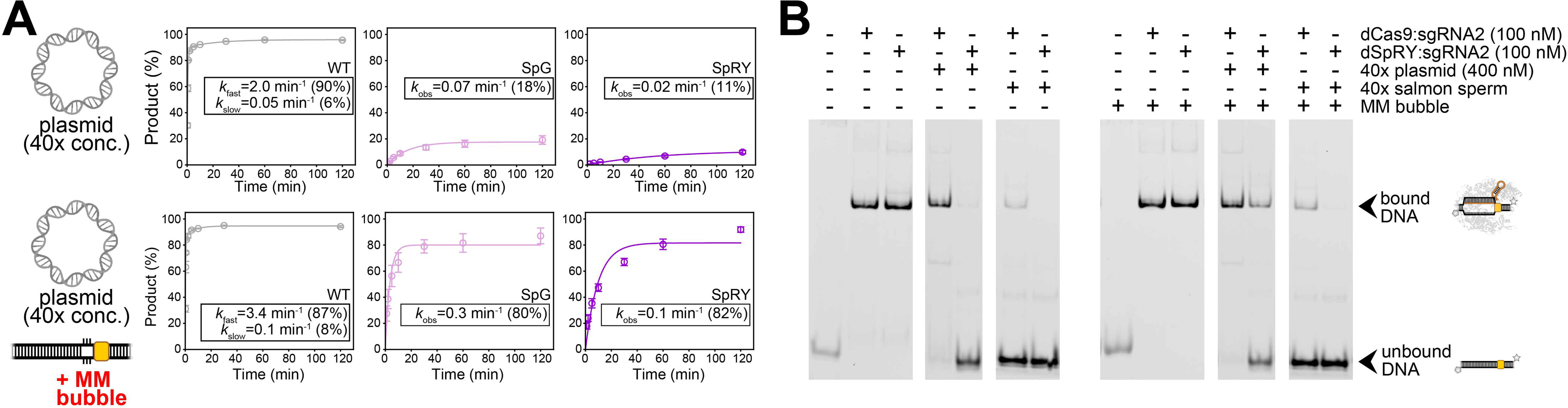
Facilitating on-target unwinding is insufficient to overcome off-target interactions for PAM-relaxed Cas9 variants. **(A)** Time-course analysis of average DNA cleavage products (n=3) for sgRNA2 at 10 mM Mg^2+^ and 37°C with a 40× plasmid competitor and an MM bubble. See also Fig. S6C for NTS cleavage. Note that the cleavage data without MM bubbles shown in Fig. 5D is also shown here (top) for direct comparison. **(B)** EMSA analysis of target DNA binding by dCas9 or dSpRY, comparing conditions with and without competitor DNA, with or without an MM bubble. See also Fig. S6D for data on sgRNA1. All the errorbars represents standard deviations.

To further investigate, we performed EMSA assays on dCas9 and dSpRY binding to a target DNA with a MM bubble (Fig. 6B). In the presence of a 40× plasmid competitor, dSpRY’s on-target binding improved from 2% to 29% with a MM bubble while dCas9’s binding remained unaffected (∼90%). However, with 40× salmon sperm DNA, dSpRY’s on-target binding was completely abolished, regardless of the MM bubble (Fig. 6B; Fig. S6D), highlighting that strong non-specific DNA binding remains a major barrier for SpRY. ChIP-seq data in human cells supported this, showing diminished on-target signals and higher non-specific binding for dSpRY compared to dCas9 (Fig. S6E-G; Note S2). Therefore, while reducing the energy barrier for on-target unwinding significantly improves SpG and SpRY’s cleavage activity, it is insufficient to overcome the strong non-specific DNA binding that continues to hinder efficiency under competitive conditions.

## DISCUSSION

Understanding the molecular basis for genome editing efficiency by CRISPR-Cas9 is important for both fundamental knowledge and practical improvement of editing tools used in clinical and agricultural applications. In this study we investigated the observed loss of editing efficacy in Cas9 variants derived from the highly effective *Spy*Cas9 enzyme but lacking specificity for the target-adjacent PAM sequence^5,6,28^. PAM-relaxed variants, such as SpRY, are slower to engage correct target DNA sequences^28^ and bind off-target sequences more frequently *in vitro*^28^ and in bacteria^35^.

Our findings show that SpRY’s inefficiency stems from fundamental issues in the target capture process, which follows a two-step mechanism: initial PAM binding followed by DNA unwinding. WT *Spy*Cas9 excels due to its selective, low-affinity PAM binding that transitions rapidly to unwinding, enabling high-efficiency genome editing, including base and prime editing^36–38^ and transcriptional regulation^39,40^. In contrast, two factors drive SpRY’s inefficiency (Fig. 7). First, it becomes kinetically trapped in a nonproductive binding state due to relaxed PAM specificity, leading to promiscuous interactions and a prolonged search time. Second, even after reaching the target site, the interactions responsible for kinetic trapping also slow target DNA unwinding. While artificially enhancing on-target unwinding aids enzyme performance, it doesn’t overcome SpRY’s persistent off-target interactions that continue to reduce efficiency. Based on these findings, we propose that increasing PAM specificity, while limiting initial binding affinity, accelerates genome editing.

**Fig. 7.**
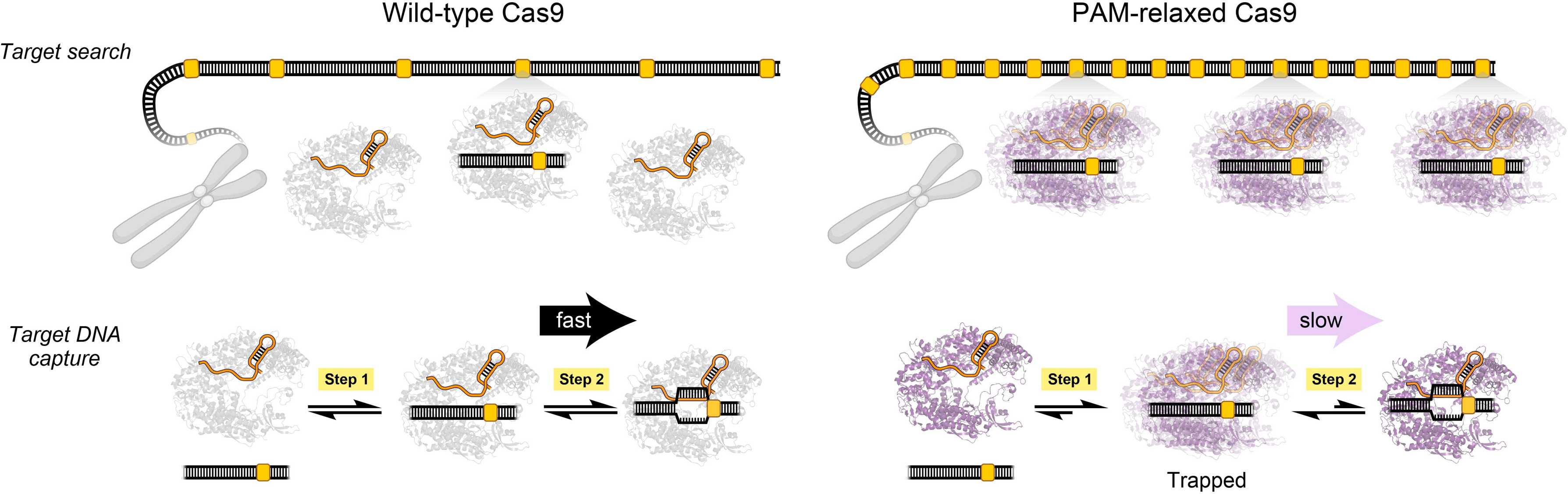
A model explaining the reduced efficiency of PAM-relaxed Cas9 variants. The reduced genome editing efficiencies of PAM-relaxed Cas9 variants are explained by two factors: (Top) PAM-relaxed Cas9 variants display prolonged target search times, largely due to strong non-specific binding, which reduces the pool of free RNPs available for on-target identification. (Bottom) Upon reaching the target site, these variants become kinetically trapped in a low-energy initial binding complex and unwind DNA slowly.

Despite the potential utility of a single Cas enzyme that can target all possible PAM sequences while maintaining robust on-target genome editing activity, our data show that specific PAM binding is fundamental to RNA-guided genome manipulation. While it might seem intuitive that extending the length of PAM motifs could improve editing efficiency, previous studies have shown that longer PAM motifs, such as those recognized by *Spy*Cas9 variants like EQR (PAM=NGAG) and VRER (PAM=NGCG), do not necessarily improve editing efficiency^41^. Indeed, both EQR and VRER demonstrated similar or even reduced DNA cleavage efficiencies relative to WT *Spy*Cas9 in the presence of competitor DNA (Fig. S7). These results align with a recent theoretical study suggesting that two-base-pair PAMs optimize target search speed for Cas9^42^. Together, these findings underscore the importance of carefully balancing PAM length and affinity when developing a comprehensive “PAM-catalog” of Cas9 or related RNA-guided enzymes^43^.

Furthermore, our findings suggest that optimizing ground-state PAM binding may be insufficient for developing more efficient genome editors. Future engineering efforts to improve genome editing efficiency should focus on reducing the energy barrier between initial DNA encounters and DNA unwinding, preventing enzyme stalling at intermediate states. This shift toward optimizing transition-state kinetics presents a promising new strategy for advancing the efficiency of other RNA-guided genome editors.

In summary, these results highlight a two-step target capture mechanism ensuring high-efficiency Cas9-mediated genome editing and reveal the inherent limitations of PAM-relaxed enzymes. This work not only provides crucial insights into the kinetic bottlenecks that reduce editing efficiency but also offers a framework for future engineering strategies to overcome these challenges. Extending these studies to other CRISPR-Cas enzymes and ancestral RNA-guided endonucleases, such as IscB and TnpB, could reveal the central principles governing RNA-guided target search and recognition. Ultimately, these insights will drive the next generation of genome editing technologies, enhancing both precision and versatility in therapeutic and functional applications.

## Acknowledgements

We thank Mr. Peter Yoon and other members of the Doudna lab and the Bryant lab for helpful discussions. We would also like to acknowledge Ms. Netravathi Krishnappa (NGS Core Operations Manager and Sequencing Specialist, Center for Translational Genomics, Innovative Genomics Institute, UC Berkeley) and Dr. Suhua Feng (Associate Project Scientist, High-Throughput Sequencing Core, Broad Stem Cell Research Center, UCLA) for NGS. H.S. is an HHMI Fellow of The Jane Coffin Childs Fund for Medical Research. N.A. is a recipient of the National Science Foundation Graduate Research Fellowship. P.S. is supported by the Swiss National Science Foundation Mobility fellowship (P500PB_214418). E.E.D. is supported by a fellowship award from the National Institutes of Health (F32GM153031). J.C.C. is a fellow of the Helen Hay Whitney Foundation. J.A.D. is an Investigator of the Howard Hughes Medical Institute (HHMI). This project was supported by the National Institutes of Health (U01AI142817 to J.A.D. and R01GM106159 to Z.B.). Figure cartoons generated with biorender.com.

## Author contributions

Conceptualization: H.S., Z.B., and J.A.D.; experimental studies: H.S., N.A., K.M.W., E.E.D., K.V., D.C.; data analysis: H.S., N.A., K.M.W., M.I.T., E.E.D., R.S.B., P.S.; supervision: Z.B. and J.A.D.; manuscript writing: H.S. and J.A.D. with critical input from Z.B., N.A., K.M.W., J.C.C.

## Declaration of generative AI and AI-assisted technologies

During the preparation of this work, the authors used GPT-4o to assist with text editing. After using this tool or service, the authors reviewed and edited the content as needed, and take full responsibility for the content of the publication.

## Supplemental information

Document S1: Figure S1–S8, Table S1-S17 and Note S1-S2.

## METHODS

### Plasmid constructions

Mammalian expression plasmids were derived from pSpCas9(BB)-2A-Puro (PX459) V2.0, which encodes Cas9 with N- and C-terminal nuclear localization signals (NLS) (Table S4-6) driven by a CAG promoter, and a sgRNA under control of a U6 promoter. The 2A sequence and puromycin resistance protein were removed. SpG, SpRY, and catalytically dead mutants were generated via site-directed mutagenesis and assembled using Gibson assembly. The U6 promoter, sgRNA sequence and protein sequences are detailed in Table S5-6. For ChIP-seq experiments, the sgRNA or Cas9 expression cassettes were removed by restriction digestion to generate plasmids expressing solely Cas9 protein or a sgRNA separately.

Bacterial expression vectors for Cas9 and its variants were generated by assembling gBlocks (Integrated DNA Technologies) containing specific Cas9 mutations into custom pET-based vectors using Gibson assembly (Table S4). These vectors feature a T7 promoter, followed by an N-terminal His10-tag, maltose-binding protein (MBP), a tobacco etch virus (TEV) protease cleavage site, and the respective Cas9 variants (Table S6). Cas9 and its variants used in this study include WT *Spy*Cas9, SpG, SpRY, EQR, VRER, xPBA, dCas9, dSpRY, as well as WT *Spy*Cas9-2NLS, SpG-2NLS, and SpRY-2NLS (Table S7).

### Nucleic acid preparations

sgRNAs used in biochemical and biophysical experiments were generated by *in-vitro* transcription, following the protocol described by Cofsky et al^6^. Briefly, transcription reactions contained 25 μg DNA template (Table S8) assembled by polymerase chain reaction (PCR), 20 mM of each nucleoside triphosphate, 30 mM Tris-HCl (pH 8.0), 25 mM MgCl_2_, 10 mM DTT, 0.01% (v/v) Triton X-100, 2 mM spermidine, and 100 µg/mL T7 RNA polymerase. Reactions were incubated at 37°C for 4 h, followed by purification by 8% denaturing urea polyacrylamide gel electrophoresis (UREA-PAGE). The gel containing RNA was crushed and soaked in DEPC-treated water overnight at 4°C, followed by six washes with DEPC water. sgRNA for cellular experiments were purchased from Integrated DNA Technologies, with chemical modifications at 3*’*- or 5*’*-ends for enhanced stability (details in Table S9) and resuspended in IDTE Buffer (10 mM Tris-HCl, pH 7.5; 0.1 mM EDTA). All sgRNA were annealed (80 °C for 2 min, then directly placed on ice) before use.

All the DNA oligonucleotides used in this study were synthesized by Integrated DNA Technologies. DNA primers were ordered with standard desalting, while DNA oligonucleotides for biochemical assays were HPLC-purified. For the double-stranded DNA (dsDNA) substrates used in DNA cleavage assays and electrophoretic mobility shift assays (EMSA), the target strand (TS) was 5*’*-labeled with FAM, and the non-target strand (NTS) was 5*’*-labeled with Cy5. In the 2AP fluorescence assay, the NTS of the dsDNA substrate contained a single 2-aminopurine (2AP) modification within the target sequence, while the TS remained unlabeled. For the short dsDNA competitors used in competition cleavage assay, the competitor DNA was unlabeled. In the permanganate footprinting assay, the TS of the dsDNA substrate was 5*’*-labeled with FAM, while the NTS remained unlabeled. All the dsDNA substrates were annealed by heating a 1:1 NTS/TS molar (unless otherwise specified) mixture of complementary single-stranded oligonucleotides to 95°C followed by gradually cooling to 35°C over 45 min in DNA Annealing Buffer (10 mM Tris-HCl, pH 8.0; 100 mM NaCl; 1 mM EDTA). The plasmid (pGGAselect) used in competition cleavage assay was ordered from New England Biolabs and prepared in-house using a HiSpeed Plasmid Mega EF kit (QIAGEN). Purified and sheared salmon sperm DNA (10 mg/mL, Invitrogen) with size of 200-500 bps was purchased from ThermoFisher Scientific, in which the sheared ends may comprise a range of structures and single strand overhangs.

The A260 absorbance of both DNA and RNA oligonucleotides was measured using a NanoDrop spectrophotometer (ThermoFisher Scientific). Concentrations were calculated based on previously reported extinction coefficients^44^. The sequences of all DNA and RNA oligonucleotides used in this study are listed in Table S9-17.

### Tissue culture and DNA transfection

HEK293T cells (UC Berkeley Cell Culture Facility) were cultured in DMEM (Corning) supplemented with 10% fetal bovine serum (Gibco) and 100 U/ml penicillin-streptomycin (Gibco). 5×10^6^ cells were seeded into a 10 cm tissue culture dish (Corning) and transfected with 10 µg plasmid DNA using Lipofectamine 3000 according to the manufacturer’s instructions. Cells were harvested at 8, 16, 24, and 72 h post-transfection, followed by genomic DNA extraction.

### Electroporation of ribonucleoprotein

100 pmol Cas9 (WT *Spy*Cas9-2NLS, SpG-2NLS, or SpRY-2NLS) was diluted in Protein Storage Buffer (20 mM HEPES, pH 7.5; 150 mM KCl; 10% (v/v) glycerol; 1 mM TCEP) and mixed with an equal volume of 125 pmol sgRNA in IDTE Buffer (1:1.25 molar ratio) for a total of 5 µL RNP. RNP was allowed to complex at room temperature for 10 min prior to diluting as necessary in the same mixture of Protein Storage Buffer and IDTE Buffer (1:1 by volume). HEK293T cells were trypsinized, washed once with PBS, and resuspended in supplemented Lonza SF Cell Line Nucleofector Solution. 2×10^5^ cells in 20 µL SF solution were mixed with 5 µL of pre-complexed RNP and 25 µL was transferred to the cuvette. Electroporation was performed with a Lonza 4D-Nucleofector 96-well Unit using the pulse code DS-150. 75 µL pre-warmed media was added to the cells immediately after electroporation and the contents of each cuvette were divided among three wells of a 96-well culture plate containing pre-warmed media. Cells were incubated for 72 h, at which point the three wells from each nucleofection were re-pooled prior to genomic extraction.

### Genomic DNA extraction

HEK293T cells were trypsinized and pellet for genomic DNA extraction. Genomic DNA was extracted from HEK293T cells by resuspending live or snap-frozen cell pellets in QuickExtract DNA Extraction Solution (Biosearch Technologies) followed by incubation at 65°C for 15 min, 68°C for 15 min, then 98°C for 10 min.

### Next-Generation Sequencing (NGS) library prep and sequencing

The extracted genomic DNA was then subjected to NGS library preparation through a two-step PCR process using Q5 High-Fidelity 2× Master Mix (New England Biolabs). The first PCR step involved amplifying the genomic loci and attaching adapter sequences (primers listed in Table S10), and the second step involved adding illumina index and P5, P7 sequences. Each sample was pooled at equimolar concentrations into a library and sequenced on the Illumina NextSeq 1000/P1 platform (2×150 bp) to obtain at least 100,000 reads per sample.

The paired-end sequencing reads were first trimmed using the BBDuk tool in Geneious Prime (https://www.geneious.com/prime), setting a minimum quality threshold of 20 and a minimum length of 20 bps. Following trimming, the reads were merged with the BBmerge tool in Geneious Prime. These merged reads were then analyzed with CRISPResso2 (https://github.com/pinellolab/CRISPResso2) to quantify the indel rate, according to the methods described by Ma et al^45^.

### Droplet digital PCR (ddPCR) for double-strand breaks

The ddPCR assay was designed following methods in previous studies^46,47^. Two ∼150 bp amplicons were designed: Amplicon 1 spans the EMX1 target site, while Amplicon 2 is located about 200 bp downstream. A double-strand break (DSB) at the target site would prevent amplification of Amplicon 1, whereas Amplicon 2 would remain unaffected. Probes for the target and reference amplicons were ordered labeled with the fluorophores FAM and HEX, respectively.

The DSB-ddPCR reactions were assembled with ddPCR Supermix for Probes (No dUTP), 900 nM of each primer, 250 nM of each probe, and 15 ng of genomic DNA, quantified via Qubit dsDNA Quantification Assay (ThermoFisher Scientific). The sequences for all the primers and probes are provided in Table S11. Droplets were formed using a Bio-Rad QX200 Droplet Generator according to the manufacturer’s instructions. Thermal cycling was performed according to the following protocol: (95°C for 10:00; 40 cycles of 94°C for 0:30 followed by 63.3°C for 3:00; 97°C for 10:00; held at 4°C). Droplets were analyzed on a Bio-Rad X200 Droplet Reader the next day and data were analyzed using the QX Manager Software. The percentage of alleles with double-strand breaks was calculated from the number of droplets that amplified the target amplicon (labeled with FAM) compared to those that amplified the reference amplicon (labeled with HEX). The equation utilized is as follows:

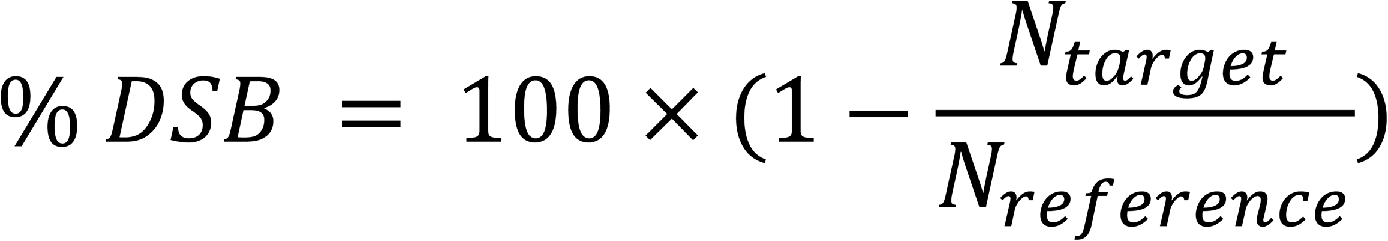

where *N*_target_ and *N*_reference_ represents the number of droplets that amplified the target amplicon and the reference amplicon, respectively.

### Western blot

Fresh or snap-frozen cell pellets were resuspended in RIPA Lysis and Extraction Buffer (ThermoFisher Scientific) containing 1× Protease Inhibitor Cocktail (ThermoFisher Scientific). Lysis was allowed to continue for 30 min, occasionally passing lysate through a syringe needle to thoroughly shear nuclei. Debris was removed by centrifugation at 20,000×g for 20 min at 4°C and the total protein concentration in the supernatant was measured using a Pierce BCA assay kit. 4 µg lysate was denatured at 95°C for 5 min and resolved by SDS-PAGE. Protein was transferred to a PVDF membrane. The membrane was blocked with Blocking Buffer (1× PBS; 0.1% (v/v) Tween-20, 5% (m/v) milk) for 1 h at room temperature, incubated with primary antibody in Blocking Buffer overnight at 4°C, washed three times with PBS/0.05% Tween-20 for 5 min each, incubated with dye-conjugated secondary antibody in Blocking Buffer for 1 h at room temperature and washed three times again with PBS/0.05% Tween-20 for 5 min each. Protein bands were visualized on an LI-COR Odyssey CLx with Image Studio v5.2 software using 700 nm and 800 nm channels.

### Quantitative reverse transcription PCR (RT-qPCR) for sgRNA

At the indicated time points post-transfection, 1×10^6^ cells were washed, resuspended in QIAzol, and stored at −80°C. RNA extraction was performed using an miRNeasy Mini Kit according to the manufacturer’s instructions. 1 µg total RNA per sample was used for reverse transcription using the High-Capacity cDNA Reverse Transcription Kit according to the manufacturer’s instructions. qPCR was performed using Power SYBR Green PCR Master Mix and primers specific for either the Cas9 sgRNA or beta-actin (Table S12) on a CFX96 Touch Real-Time PCR Detection System.

### ChIP-seq

5×10^6^ HEK293T cells were seeded on a tissue-culture treated 10 cm plate the day before transfection. Cells were transfected using Lipofectamine 3000 with 7.5 µg of plasmid encoding dCas9 or dSpRY protein and 12.5 µg of plasmid encoding an EMX1-targeting sgRNA. Cells were passaged into a 15 cm plate one day post-transfection. At 72 h post-transfection, 50 million cells were harvested and fixed in 1% formaldehyde in PBS for 10 min. Fixation reaction was quenched with the addition of 125 mM glycine, cells were washed twice with ice-cold PBS, and pellets were flash-frozen in liquid nitrogen and stored at −80°C. Cells were thawed on ice, resuspended in Lysis Buffer 1 (50 mM HEPES, pH 7.5; 140 mM NaCl; 0.5% IGEPAL CA-630; 0.25% (v/v) Triton X-100; 10% (v/v) glycerol; 1 mM EDTA; supplemented with protease inhibitor cocktail, cOmplete EDTA-free, Roche) and incubated at 4°C with rotation for 10 min. Nuclei were pelleted, resuspended in Lysis Buffer 2 (10 mM HEPES, pH 7.5; 200 mM NaCl; 1 mM EDTA; 0.5 mM EGTA; 200 µg/mL RNase A; supplemented with protease inhibitor cocktail, cOmplete EDTA-free, Roche) and incubated at 4°C with rotation for 10 min. Nuclei were pelleted again and resuspended in 1.5 mL Sonication Buffer (10 mM HEPES, pH 7.5; 100 mM NaCl; 1 mM EDTA; 0.5 mM EGTA; 0.1% sodium deoxycholate; 0.5% N-Lauroylsarcosine; supplemented with protease inhibitor cocktail, cOmplete EDTA-free, Roche). Genomic DNA was sheared using a BioRuptor Pico (10 cycles, 30 s on, 30 s off). Anti-Cas9 antibody (Diagenode) was co-incubated with Protein A beads for ≥ 6 h prior to immunoprecipitation. Lysate was mixed with antibody-bound magnetic beads and rotated at 4°C overnight. Beads were washed 5 times with RIPA Buffer (50 mM HEPES, pH 7.5; 500 mM LiCl; 1 mM EDTA; 1% IGEPAL CA-630; 0.7% sodium deoxycholate), once with TEN Buffer (10 mM Tris-HCl, pH 8.0; 50 mM NaCl; 1 mM EDTA), and once with TE Buffer, with 3 min of rotation at 4°C between each wash, prior to elution in TES Buffer (50 mM Tris-HCl, pH 8.0; 10 mM EDTA; 1% SDS). 8 units of Proteinase K were added to samples and incubated at 55°C for 1 h. Reverse crosslinking was performed at 65°C overnight. DNA was purified using phenol-chloroform precipitation.

Sequencing libraries were generated with the NEBNext Ultra II DNA Library Prep Kit for Illumina according to the manufacturer’s instructions. DNA was purified using SPRIselect beads and library quality was confirmed on an Agilent 2100 Bioanalyzer prior to sequencing. Sequencing was performed on an Illumina NovaSeq 6000 using a 100×100 paired-end configuration and an S1 Reagent Kit (v1.5), achieving an average depth of ∼160 million reads per sample. FASTQ files were aligned to GRCh38 using Bowtie 2 (version 2.5.2)^48^ with parameters ’--no-discordant --local --no-mixed --maxins 1000’ to limit alignment artifacts for peak calling. Alignments were deduplicated, normalized to Counts Per Million (CPM) and converted into BigWig format using deepTools2 (version 3.5.1)^49^. The processed data was visualized in Integrative Genomics Viewer (IGV)^50^.

To assess the quality and reproducibility of the ChIP-seq experiment, pseudoreplicates were generated. Mapped reads from the original alignments were extracted and downsampled with seqtk (version 1.3) (https://github.com/lh3/seqtk) to create three pseudoreplicates, each with 50 million paired-end reads. Peak calling was conducted with MACS2 (version 2.2.9.1)^51^, using pseudoreplicates of the corresponding apo sample as background controls and a p-value threshold of 0.01. The resulting peaks were used to calculate the signal-to-noise ratio, defined as the ratio of reads within peaks to those outside. To further investigate noise, pseudoreplicate peak reproducibility was determined using the Irreproducible Discovery Rate (IDR) framework^52^ with the IDR package (version 2.0.4.2) (https://github.com/kundajelab/idr) at a false discovery rate of 0.05. Weak SNR (WT = 2.89, SpRY = 2.78) and low reproducibility (WT = 0.228%, SpRY = 0.151%) indicated suboptimal IP efficiency and background noise, rendering peak calling unfeasible. Instead, coverage was evaluated in genomic regions containing or lacking the seed region and PAM sequences—5*’*-AAGAANGG-3*’* for WT and 5*’*-AAGAANRN-3*’* for SpRY— specifically within a 25-bp range around the seed region and PAM. Coverage metrics, quantified as average reads per kilobase million (RPKM), were computed for designated regions in deduplicated bams (Picardtools version 2.21.9) (https://broadinstitute.github.io/picard/) and blacklisted regions of the genome were excluded, according to best practices^53^. Background subtraction was performed using averages from the corresponding apo replicates, ensuring only like-regions were adjusted. This process was facilitated by BEDTools (version 2.29.2)^54^ and a custom Python script.

### Protein expression and purification

To express and purify Cas9 variants, we employed a modified protocol based on Cofsky et al^6^. Briefly, *E. coli* Rosetta (DE3) cells (Sigma-Aldrich), transformed with the appropriate bacterial expression plasmids, were cultured in Terrific Broth (TB) medium (ThermoFisher Scientific) containing ampicillin (0.1 mg/mL) and chloramphenicol (0.034 mg/mL). The cultures were initiated from a 1:80 dilution of an overnight starter culture and grown at 37°C. Once the optical density at 600 nm (OD_600_) reached 0.6–0.8, protein expression was induced with 0.5 mM isopropyl β-d-1-thiogalactopyranoside (IPTG) following a cold shock. Induction proceeded overnight at 16°C.

The next day, cells were pelleted by centrifugation and resuspended in Bacterial Lysis Buffer (20 mM HEPES, pH 7.5; 500 mM KCl; 10 mM imidazole; 10% (v/v) glycerol; 1 mM TCEP; supplemented with protease inhibitor cocktail, cOmplete EDTA-free, Roche). The cells were lysed by sonication and were then ultra-centrifuged. The supernatant was applied to Ni-NTA resin (QIAGEN), washed with Wash Buffer (20 mM HEPES, pH 7.5; 500 mM KCl; 30 mM imidazole; 5% (v/v) glycerol; 1 mM TCEP) and then eluted with 300 mM imidazole in the same buffer. Proteins were treated with TEV protease overnight at 4°C during dialysis in the Dialysis Buffer (20 mM HEPES, pH 7.5; 300 mM KCl; 30 mM imidazole; 5% (v/v) glycerol; 1 mM TCEP). Digested and dialyzed proteins were separated using a HisTrap column (Cytiva), and the Cas9-containing flow-through was collected. This was followed by further purification using a HiTrap Heparin HP affinity column (Cytiva), with protein elution performed via a KCl gradient from 300 mM to 1 M. The final purification step involved size-exclusion chromatography on a Superdex 200 Increase 10/300 GL column (Cytiva) in the Protein Storage Buffer (20 mM HEPES, pH 7.5; 150 mM KCl; 10% (v/v) glycerol; 1 mM TCEP). Purified Cas9 proteins were aliquoted, snap-frozen, and stored at −80°C.

The protein purification was validated using SDS-PAGE analysis. Protein samples were prepared by mixing one volume of the sample with 0.25 volume of 5× SDS Loading Dye (250 mM Tris-HCl, pH 6.8; 75 mM EDTA; 30% (v/v) glycerol; 10% SDS supplemented with bromophenol blue), heated at 90°C for 2 min, and then 10 pmol was loaded onto 10% Mini-PROTEAN TGX Precast Protein Gels (Bio-Rad), with a PageRuler Prestained Protein Ladder (ThermoFisher Scientific). The gels were stained with Coomassie Staining Buffer (30% (v/v) ethanol; 10% (v/v) acetic acid; 1g R250-coomassie) followed by destaining (40% (v/v) ethanol; 10% (v/v) acetic acid) or with instant InstantBlue Coomassie Protein Stain (Abcam). The gels were imaged using a Bio-Rad ChemiDoc system.

### DNA cleavage assay

For most DNA cleavage assay, Cas9 (or its variants) and sgRNA were co-incubated at a 1:1.25 molar ratio to form RNP complexes at 24°C or 37°C for 15 min in the 1.05× cleavage reaction buffers, each tailored to different Mg^2+^ ion concentration. The 10 mM Mg^2+^ Cleavage Buffer (1×) contained 20 mM HEPES (pH 7.5), 100 mM KCl, 10 mM MgCl_2_, 1% (v/v) glycerol, 0.5 mM DTT. The 5 mM Mg^2+^ Cleavage Buffer (1×) contained 20 mM Tris-HCl (pH 7.5), 100 mM KCl, 5 mM MgCl_2_, 5% (v/v) glycerol. The 0.2 mM Mg^2+^ Cleavage Buffer (1×) contained 20 mM HEPES (pH 7.5), 100 mM KCl, 0.2 mM MgCl_2_, 1% (v/v) glycerol, 0.5 mM DTT. The Mg^2+^ free Cleavage Buffer contained 20 mM HEPES (pH 7.5), 100 mM KCl, 0.2 mM EDTA, 1% (v/v) glycerol, 0.5 mM DTT. The reaction was initiated by adding the RNP complex (19 volumes) to the target dsDNA substrate (1 volume). For competition cleavage assays, target dsDNA substrate and competitor DNA were pre-mixed before adding the RNP complex. In the pre-formed *R*-loop cleavage experiments, Cas9, sgRNA and target dsDNA substrate were incubated in 1.05×Mg2+ free buffer at 24°C or 37°C for 2 hours, and the reaction was triggered by mixing 19 volumes of the RNP-DNA complex and 1 volume of MgCl_2_. The final concentrations were 10 nM dsDNA substrate, 100 nM Cas9, 125 nM sgRNA, 1× cleavage buffer, unless otherwise specified. For protein active fraction experiments, the reactions involved 100 nM dsDNA substrate, 100 nM Cas9 and 125 nM sgRNA, with the fraction of DNA cleaved after 2 h representing the active enzyme fraction (the active fractions ranging from 37% to 50% as shown in Fig. S1G). Cleavage reactions were conducted at either 24°C or 37°C for up to 2 h and quenched at different time points by adding an equal volume of 2× Cleavage Quenching Buffer (94% (v/v) formamide; 30 mM EDTA; 400 μg/mL heparin, supplemented with bromophenol blue). Samples were then heated at 90°C for 5 min and resolved on a 15% UREA-PAGE gel.

The gel was scanned using a Typhoon (Amersham, GE Healthcare) with excitation at 488 nm and a Cy2 emission filter (525BP20) for FAM-labeled oligonucleotide, or with excitation at 635 nm and a Cy5 emission filter (670BP30) for Cy5-labeled oligonucleotide. Band intensities were quantified using Bio-Rad ImageLab 6.1 software, and the resulting data were fitted to a mono-exponential decay or double-exponential decay using the curve_fit function in *Scipy* Python package:

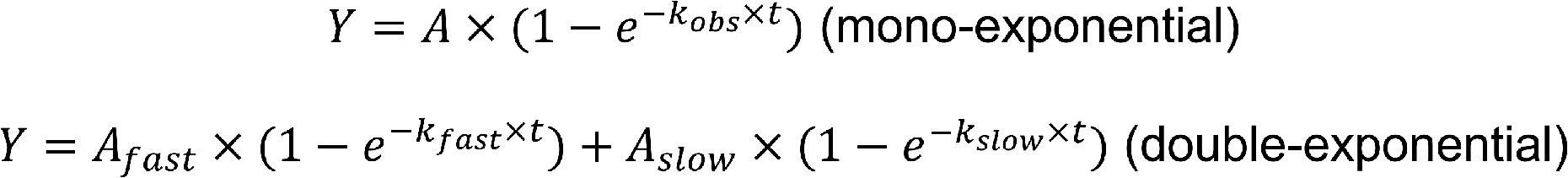

where Y is the cleaved DNA product (%) at a given time, A (or A_fast_ and A_slow_) represents the amplitude of the exponential decay, *k*_obs_ (or *k*_fast_ and *k*_slow_) represents the rate constant of the exponential decay, t is time (min). All data were initially fitted using a double-exponential model. If the second phase was poorly defined or its amplitude was close to zero, a mono-exponential model was then applied. All the experiments were performed in biological triplicates. The fitting parameters and their errors were calculated as the average and their standard deviation across the replicates.

### 2-aminopurine fluorescence assay

Cas9 and sgRNA were assembled at a 1:1.25 molar ratio of RNA to active Cas9 in 5 mM Mg^2+^ cleavage buffer (1×), and incubated at 24°C for 30 min. The reaction was initiated by adding the RNP complex to 2AP-containing dsDNA substrate at a final molar ratio of 8:1 (RNP complex to DNA) at room temperature (24°C) in a black 384 well plate. The final concentrations in the reactions were 1 μM dsDNA substrate, 8 μM Cas9 RNP, 5 mM Mg^2+^ Cleavage Buffer (1×). Fluorescence emission (λ_em_ 370 nm, λ_ex_ 320 nm) for each reaction was recorded every 20 seconds on a Cytation 5 plate reader (Biotek, software Gen v3.04). The fluorescence data over time were fitted to the monoexponential decay using the curve_fit function in *Scipy* Python package:

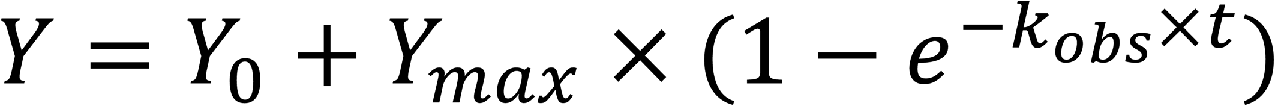

where Y is the fluorescence signal at a given time, Y_max_ is the fitted fluorescence endpoint, Y_0_ is the average value from a control reaction where the RNP complex (with a non-targeting sgRNA, sgRNA8) was added to 2AP containing dsDNA (Fig. S2B), and *k*_obs_ is the observed rate constant of the exponential decay. Each reaction was carried out in triplicate, and the average fluorescence values were used to fit the parameters, with their standard fitting errors reported.

### Permanganate DNA footprinting assay

We adapted a similar protocol based on Cofsky et al^6^. Briefly, a dsDNA substrate with an NTS:TS ratio of 1.5:1, where the TS was 5*’*-labeled with FAM, was mixed with Cas9 RNP in a 1.1× Permanganate Reaction Buffer and incubated at 30°C for 15 min. The 1× Permanganate Reaction Buffer included 20 mM Tris-HCl (pH 7.9), 24 mM KCl, 5 mM MgCl_2_, 0.1 mg/mL BSA, 0.01% (v/v) Tween-20, 1 mM TCEP. The reaction was initiated by combining the RNP-DNA complex (9 volume) with 50 mM KMnO_4_ (1 volume) and the mixture was incubated for 2 min at 30°C. The final concentrations in the reaction were 200 nM dsDNA substrate, 16 μM Cas9 RNP, 5 mM of KMnO_4_ and 1× Permanganate Reaction Buffer. The reaction was quenched by the addition of equal volume of 2× Permanganate Quench Solution (2 M β-mercaptoethanol, 30 mM EDTA).

The DNA from the quenched reaction was extracted using phenol:chloroform:isoamyl (PCI) and precipitated with 2.5 volumes of 100% ethanol at −20°C overnight. The ethanol precipitate was then washed twice using 70% ethanol and then resuspended in 70 µL of 10% (v/v) piperidine, followed by incubated at 90°C for 30 min. The samples were lyophilized using vacuum concentration, resuspended in 10 µL of 1× UREA-PAGE Loading Dye (50% (v/v) formamide; supplemented with bromophenol blue) and then analyzed on a 15% UREA-PAGE gel. The gels were imaged by Typhoon as described above.

The probability of cleavage occurring at a specific thymine i is defined as:

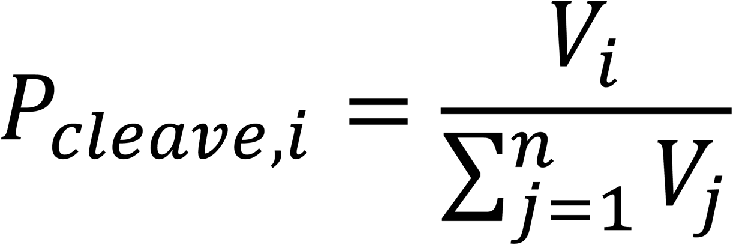

where V_i_ represents the volume of band corresponding to the cleavage at position i within a lane containing n bands (with band 1 being the smallest cleavage fragment and band n corresponding to the full length DNA). The oxidation probability of thymine i is then defined as:

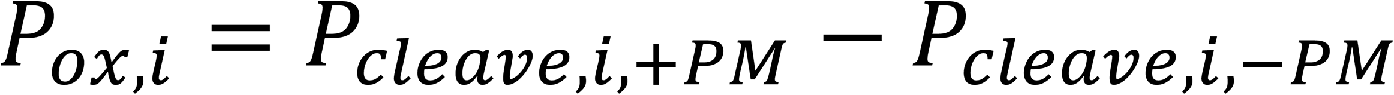

where +PM refers to the experiment with 5 mM KMnO_4_ and -PM refers to the control experiment without permanganate. An extensive description of the analysis is provided in Cofsky et al^55^.

### EMSA

Binding reactions were performed by incubating 100 nM catalytically inactive Cas9 RNP (dCas9 or dSpRY) with 10 nM target dsDNA substrate in the 10 mM Mg^2+^ Cleavage Buffer for 30 min. For binding experiments involving competitor DNA, the competitor was premixed with the dsDNA substrate, as done in the cleavage assay. The reactions were quenched by mixing with equal volume of 2× Native Gel Loading Solution (10 mM Tris-HCl, pH 7.5; 50% (v/v) glycerol) and analyzed on a 6% non-denaturing PAGE gel. The gels were imaged by Typhoon as described above.

In binding experiments of competitor DNA (without target dsDNA substrate), the competitor DNA was added to a final concentration of 15 nM and mixed with the RNP complex in the 10 mM Mg^2+^ Cleavage Buffer. The binding reactions were incubated at 37°C for 1 h, then mixed with 0.2 volumes of 6× Agarose Gel Loading Dye (Purple, no SDS, New England Biolabs) and analyzed on a 1% agarose gel containing SYBR Safe. The gels were imaged by a Bio-Rad ChemiDoc system.

### Gold rotor bead tracking (AuRBT)

Bottom-constrained DNA tethers were assembled by ligating restriction enzyme digested PCR products. These products included a segment with multiple incorporated dUTP-digoxigenin (Roche) for creating a torsionally constrained attachment, and another segment using a 5*’*-modified PCR primer (Integrated DNA Technologies) for an unconstrained attachment. The sequence of interest, which contained the 20-bp target sequence adjacent to either an NGG site (PAM), or an NCG site (no PAM controls), was assembled by annealing two 5*’* phosphorylated oligonucleotides (Integrated DNA Technologies). Detailed information on the tether construction, including building blocks, can be found in Fig. S8 and Table S15-17.

Experiments were performed on a custom-built AuRBT microscope, as previously described^29,56^. The rotor beads used were streptavidin-coated gold nanospheres (Cytodiagnostics, ACC-60-04-15) with a nominal diameter of 60 nm. For all AuRBT measurements, the DNA was held under 7 pN of tension in C9T Buffer (20 mM Tris-HCl, pH 7.5; 100 mM KCl; 5 mM MgCl_2_; 0.1 mM EDTA; 1 mM TCEP; supplemented with 0.2 mg/mL BSA). Movies were recorded at a frame rate of 5 kHz at room temperature (approximately 22 ± 1°C).

To ensure that the DNA tether has made stable attachments, we first record the thermal fluctuations of the rotor bead in C9T Buffer for at least 5 min. Following this, we introduced the RNP complexes. These Cas9-sgRNA complexes were formed by incubating the Cas9 with a 1.25× molar excess of sgRNA for 10 min at 37°C in C9T Buffer. The RNP was diluted to the specified concentrations in C9T Buffer supplemented with 0.2 mg/mL BSA before being introduced into the channel. Data sets for each combination of Cas9 protein, DNA sequence, and sgRNA sequence were compiled across at least two imaging sessions.

AuRBT imaging data were processed following the methods previously described^12,29^. We assume that any deviations from zero twist are due to RNP interactions leading to the unwinding of B-DNA, with helicity of 10.5 bp/turn. To analyze these unwinding events, we first filtered the data to 500 Hz and then employed the Steppi change-point analysis tool^30^, modeling the data as originating from an Ornstein-Uhlenbeck process. Global stiffness and nearest neighbor coupling parameters were fixed by analyzing a portion of the trace before the introduction of any RNP. We also set the level slope parameter to zero. The only free model parameters were the mean position of the rotor bead and the change point time.

For the data collected with WT (or dCas9) and sgRNAs containing a 3-bp match to the target DNA, we classified the DNA as “unwound” if the change in mean position of the rotor bead exceeded 1 bp; otherwise, the DNA is classified as “closed”. Consecutively scored events in the same state were merged. To retain the asymmetry in the raw scored data, we also merged states where the DNA was overwound by more than 1 bp. The extent of unwinding in these merged states was calculated as the weighted average of contributing states, with weights based on their lifetime, and the dwell times of these states were summed to obtain the total time spent in each state. The same analysis protocol was applied to no PAM control experiments. The average lifetime in the unwound state was calculated by fitting the dwell time distribution of states in the unwound cluster to a 100-ms left-censored double exponential using maximum likelihood estimation in MEMLET^57^.

Similar experiments were conducted with SpRY. For Target2, the sgRNA (sgRNA4.1) used consisted of up to 3-bp matching from the seed to the DNA below the rotor bead. After introducing SpRY:sgRNA4.1 to the tether, we observed a concentration-dependent and prolonged shift in the rotor bead’s mean angle, reflecting unwinding of the DNA helix (Fig. 4E, 4F). This unwinding was interpreted as SpRY interacting specifically or non-specifically with the DNA. The mean change in DNA twist was calculated by subtracting the initial mean rotor angle from the final mean rotor angle after SpRY introduction. These mean changes in DNA twist as a function of SpRY concentration were fit to a Langmuir model to estimate the dissociation constant *K*_D_ and the saturating angle ΔΘ_sat_

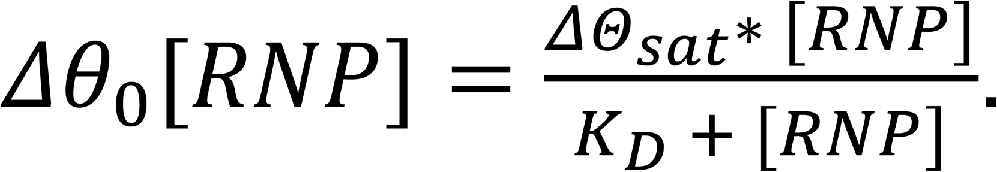

A similar analysis was applied to the non-specific binding events observed in the dSpRY:sgRNA3 experiments on Target1, where sgRNA3 has 20-bp match to the target DNA (to be introduced shortly) (Fig. S3D). For the data analysis in these experiments, the mean change in DNA twist was instead calculated by subtracting the initial mean rotor angle before SpRY introduction from the final mean rotor angle, which was determined by the averaged positions of the closed states.

For the data collected with sgRNA3 (20-bp match to Target1, respectively), states were categorized into “closed” (C), “intermediate” (I), or “open” (O) clusters and then merged as previously described^12^. State boundaries were set to align with state definitions from previous work^12^. For experimental traces obtained using SpRY or dSpRY, data were re-zeroed so that the predominant closed state corresponded to Δθ_0_ = 0 bp, accounting for minor unwinding from off-target binding that persisted throughout the data collection period. Transition rates between the three states were calculated by dividing the number of transitions from state i to state j by the total time spent in state i. Although direct transitions between the closed and open states were rare, transitions among all three states were considered, and transition rate constants (*k*_C→I_, *k*_I→C_, *k*_I→O_, *k*_O→I_, *k*_C→O_, and *k*_O→C_) were calculated (Table S1). Nonlinear least-squares fitting of a linear model for dCas9 and a hyperbolic saturation model for dSpRY was performed using MATLAB to fit *k*_C→I_ versus RNP concentration. Error bars were calculated assuming Poisson statistics.

Cartoons of free energy landscapes (Fig. 3C) were drawn with reference to the kinetic models shown in Fig. 3B^12^. Apparent equilibrium constants were computed using

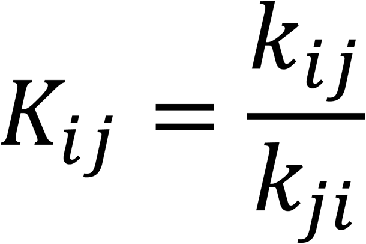

where *k_ij_* represents the transition rate from state i to state j. Estimated free energy differences were then computed using

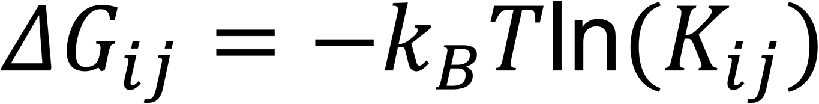

where *k*_B_ represents the Boltzmann constant and T is temperature. The landscapes were constructed using a concentration of 100 nM RNP. Well positions for C_bound_, I, and O states were set to the average bps unwound in each merged state cluster (0 bp, 10 bp, and 20 bp, respectively), and transition state locations were informed by previous work^12^. We arbitrarily set the locations of C_free_ for illustrative purposes. Relative barrier heights were derived from estimated and calculated transition rates and represented as

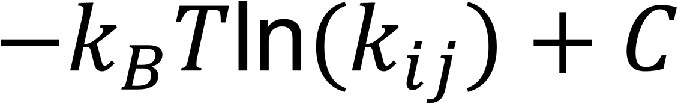

where C = 7 *k*_B_T is an arbitrary constant.

