## Supplementary material for "Rapid two-step target capture ensures efficient CRISPR-Cas9-guided genome editing": Document S1

### Supplemental Figures

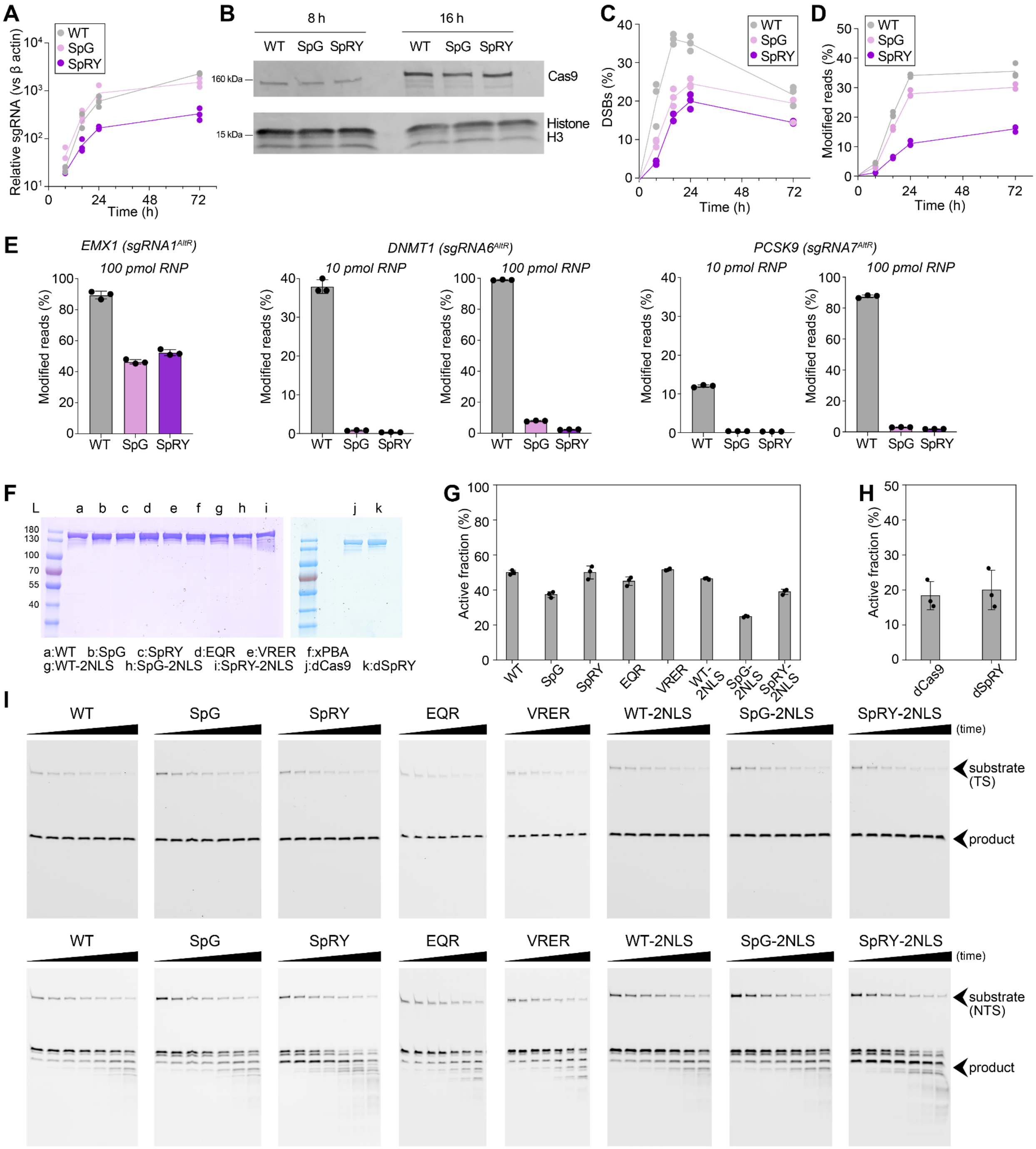


**Fig. S1. Additional results from human cell experiments and quality control of Cas9 proteins (related to Fig. 1). (A)** Relative quantifications of the sgRNA expression levels compared to β-actin mRNA using RT-qPCR (n=3). The >5-fold reduction of sgRNA expression at 72 h in the SpRY condition could be due to plasmid self-targeting by SpRY. **(B)** Western blot analysis of Cas9 expressions in human cells at 8 h and 16 h post plasmid transfection. **(C)** Quantifications of DNA DSB at various time points (t = 8, 16, 24, 72 h) post plasmid transfection (n=3). **(D)** Quantifications of indels at various time points (t = 8, 16, 24, 72 h) post plasmid transfection (n=3). **(E)** Quantifications of indels at 72 h post RNP nucleofection (10 or 100 pmol) targeting the *EMX1* gene, *DNMT1* gene, or *PCSK9* gene (n=3). **(F)** SDS-PAGE analysis of the Cas9 proteins used in the experiments. **(G)** Assessment of active Cas9 fractions through the percentage of cleaved dsDNA substrate (100 nM) after a 2-hour incubation with 100 nM Cas9 and 125 nM sgRNA2 at 10 mM $Mg^{2+}$ and 37°C (n=3). **(H)** Assessment of active dCas9/dSpRY fractions based on the percentage of dsDNA substrate (100 nM) bound, measured using EMSA after 2-hour incubation with 100 nM Cas9 and 125 nM sgRNA2 at 10 mM $Mg^{2+}$ and 37°C (n=3). **(I)** Confirmation of Cas9 proteins activity via DNA cleavage assays conducted on 15% UREA-PAGE gels, monitoring DNA cleavage over time for the (top) target strand (TS) and (bottom) non-target strand (NTS). The time points analyzed were 1 min, 2 min, 5 min, 10 min, 30 min, 60 min and 120 min.

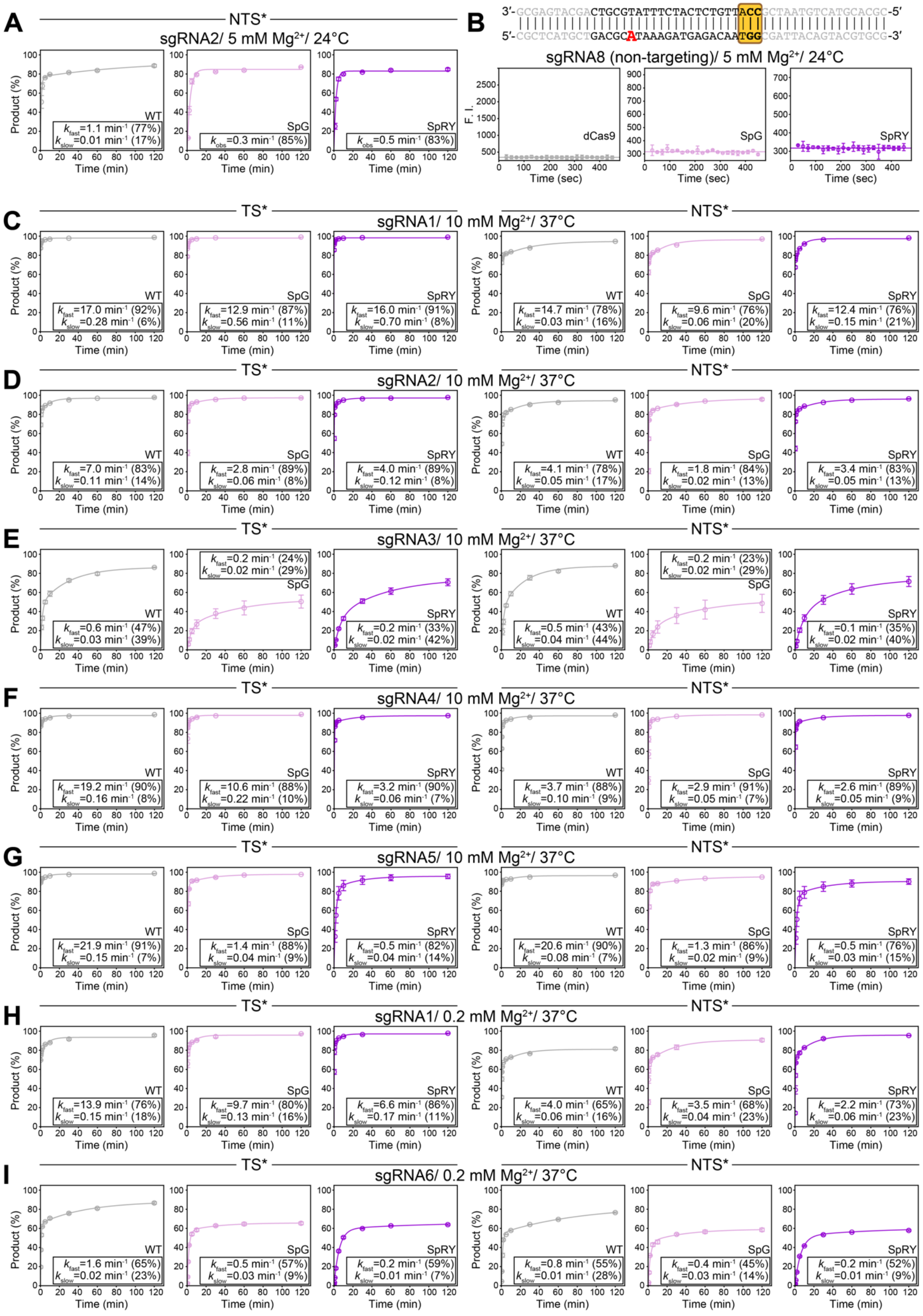

A
NTS*
sgRNA2/ 5 mM Mg2+/ 24°C
WT kfast=1.1 min-1 (77%) kslow=0.01 min-1 (17%)
SpG kobs=0.3 min-1 (85%)
SpRY kobs=0.5 min-1 (83%)
B
3′-GCGAGTACGACTGCGTATTTCTACTCTGTTACCGCTAATGTCATGCACGC-5′
5′-CGCTCATGCTGACGCATAAAGATGAGACAATGGCGATTACAGTACGTGCG-3′
sgRNA8 (non-targeting)/ 5 mM Mg2+/ 24°C
dCas9
SpG
SpRY
F. I.
Time (sec)
C
TS*
sgRNA1/ 10 mM Mg2+/ 37°C
NTS*
WT kfast=17.0 min-1 (92%) kslow=0.28 min-1 (6%)
SpG kfast=12.9 min-1 (87%) kslow=0.56 min-1 (11%)
SpRY kfast=16.0 min-1 (91%) kslow=0.70 min-1 (8%)
WT kfast=14.7 min-1 (78%) kslow=0.03 min-1 (16%)
SpG kfast=9.6 min-1 (76%) kslow=0.06 min-1 (20%)
SpRY kfast=12.4 min-1 (76%) kslow=0.15 min-1 (21%)
D
TS*
sgRNA2/ 10 mM Mg2+/ 37°C
NTS*
WT kfast=7.0 min-1 (83%) kslow=0.11 min-1 (14%)
SpG kfast=2.8 min-1 (89%) kslow=0.06 min-1 (8%)
SpRY kfast=4.0 min-1 (89%) kslow=0.12 min-1 (8%)
WT kfast=4.1 min-1 (78%) kslow=0.05 min-1 (17%)
SpG kfast=1.8 min-1 (84%) kslow=0.02 min-1 (13%)
SpRY kfast=3.4 min-1 (83%) kslow=0.05 min-1 (13%)
E
TS*
sgRNA3/ 10 mM Mg2+/ 37°C
NTS*
WT kfast=0.6 min-1 (47%) kslow=0.03 min-1 (39%)
SpG kfast=0.2 min-1 (24%) kslow=0.02 min-1 (29%)
SpRY kfast=0.2 min-1 (33%) kslow=0.02 min-1 (42%)
WT kfast=0.5 min-1 (43%) kslow=0.04 min-1 (44%)
SpG kfast=0.2 min-1 (23%) kslow=0.02 min-1 (29%)
SpRY kfast=0.1 min-1 (35%) kslow=0.02 min-1 (40%)
F
TS*
sgRNA4/ 10 mM Mg2+/ 37°C
NTS*
WT kfast=19.2 min-1 (90%) kslow=0.16 min-1 (8%)
SpG kfast=10.6 min-1 (88%) kslow=0.22 min-1 (10%)
SpRY kfast=3.2 min-1 (90%) kslow=0.06 min-1 (7%)
WT kfast=3.7 min-1 (88%) kslow=0.10 min-1 (9%)
SpG kfast=2.9 min-1 (91%) kslow=0.05 min-1 (7%)
SpRY kfast=2.6 min-1 (89%) kslow=0.05 min-1 (9%)
G
TS*
sgRNA5/ 10 mM Mg2+/ 37°C
NTS*
WT kfast=21.9 min-1 (91%) kslow=0.15 min-1 (7%)
SpG kfast=1.4 min-1 (88%) kslow=0.04 min-1 (9%)
SpRY kfast=0.5 min-1 (82%) kslow=0.04 min-1 (14%)
WT kfast=20.6 min-1 (90%) kslow=0.08 min-1 (7%)
SpG kfast=1.3 min-1 (86%) kslow=0.02 min-1 (9%)
SpRY kfast=0.5 min-1 (76%) kslow=0.03 min-1 (15%)
H
TS*
sgRNA1/ 0.2 mM Mg2+/ 37°C
NTS*
WT kfast=13.9 min-1 (76%) kslow=0.15 min-1 (18%)
SpG kfast=9.7 min-1 (80%) kslow=0.13 min-1 (16%)
SpRY kfast=6.6 min-1 (86%) kslow=0.17 min-1 (11%)
WT kfast=4.0 min-1 (65%) kslow=0.06 min-1 (16%)
SpG kfast=3.5 min-1 (68%) kslow=0.04 min-1 (23%)
SpRY kfast=2.2 min-1 (73%) kslow=0.06 min-1 (23%)
I
TS*
sgRNA6/ 0.2 mM Mg2+/ 37°C
NTS*
WT kfast=1.6 min-1 (65%) kslow=0.02 min-1 (23%)
SpG kfast=0.5 min-1 (57%) kslow=0.03 min-1 (9%)
SpRY kfast=0.2 min-1 (59%) kslow=0.01 min-1 (7%)
WT kfast=0.8 min-1 (55%) kslow=0.01 min-1 (28%)
SpG kfast=0.4 min-1 (45%) kslow=0.03 min-1 (14%)
SpRY kfast=0.2 min-1 (52%) kslow=0.01 min-1 (9%)
Product (%)
Time (min)

**Fig. S2. Additional DNA cleavage assays and 2AP assays (related to Fig. 1). (A)** Time-course analysis of average NTS DNA cleavage products (n=3) with sgRNA2 in 5 mM $Mg^{2+}$ at 24°C. The average rate constants ($k_{obs}$ for a mono-exponential decay model; $k_{fast}$, $k_{slow}$ for a double-exponential decay model), with the amplitudes from the observed exponential decay are provided. For SpG and SpRY, we assume a mono-exponential decay model, as the slow phase in the double-exponential decay model is diminished and poorly defined. **(B)** (Top) The 2AP-labeled DNA construct with the 2AP residue labeled at the 15th nt from PAM in the NTS. (Bottom) Control experiments of time-course analysis of the average fluorescence signals (n=3) in the 2AP assay for sgRNA8, which is not complementary to the target dsDNA substrate, under the same conditions as Fig. 1F. These experiments were performed to determine the intercept of the exponential fitting in Fig. 1F. For WT *Spy*Cas9, catalytically dead Cas9 (dCas9) is used here. **(C-I)** Time-course analysis of average DNA cleavage products (n=3) for various guide RNA at different conditions, presented separately for TS (left) and NTS (right). The average rate constants, with the amplitudes from the observed exponential decay are provided in figure legends. Note that the TS cleavage data in panel D are also shown in Fig. 5B for direct comparison. The $k_{fast}$ of TS cleavage in panel D, G and I are also shown in Fig. 3E as a bar graph.

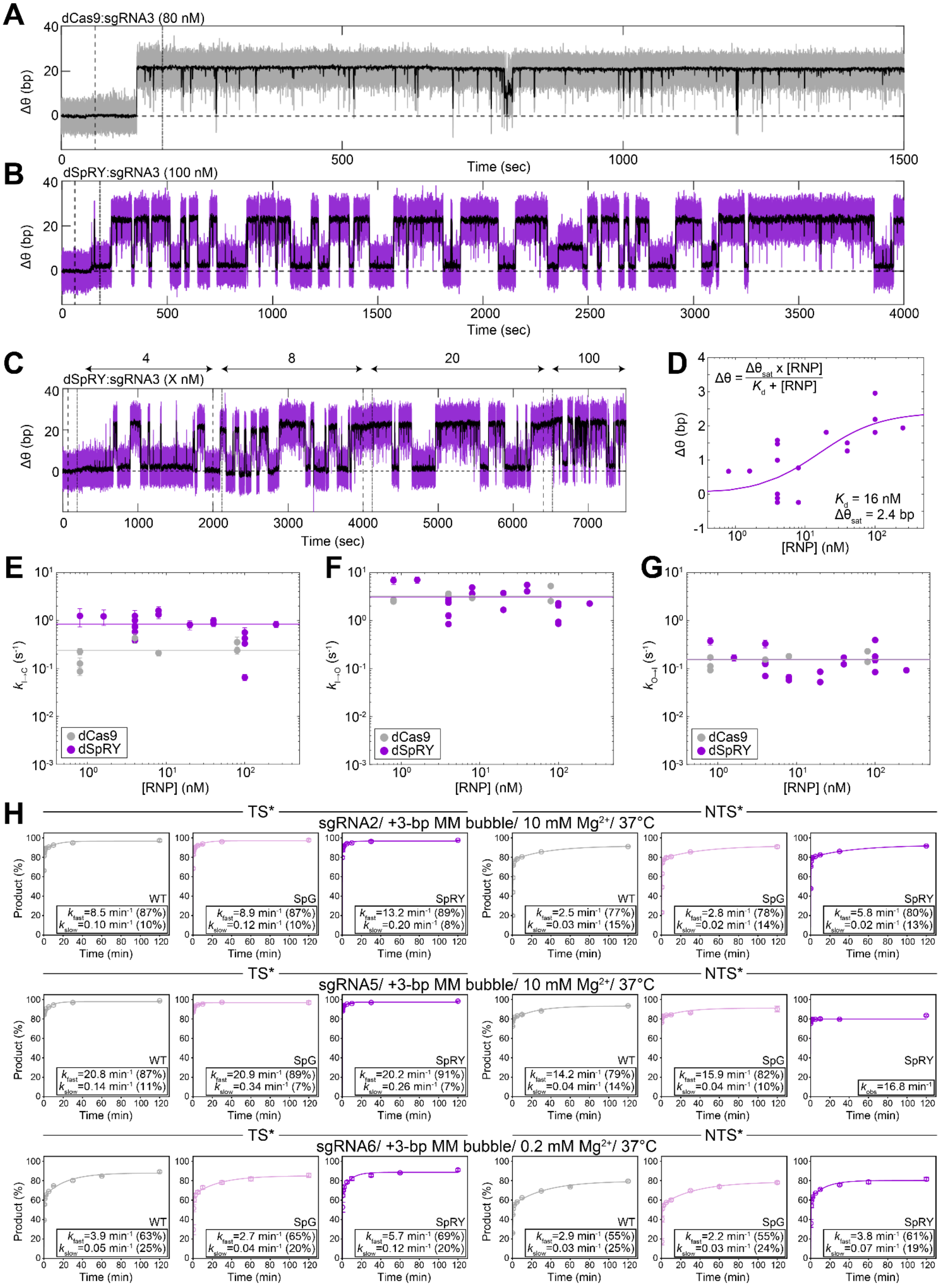
A
dCas9:sgRNA3 (80 nM)
Δθ (bp)
Time (sec)
B
dSpRY:sgRNA3 (100 nM)
Δθ (bp)
Time (sec)
C
4
8
20
100
dSpRY:sgRNA3 (X nM)
Δθ (bp)
Time (sec)
D
Δθ = Δθsat x [RNP] / Kd + [RNP]
Kd = 16 nM
Δθsat = 2.4 bp
Δθ (bp)
[RNP] (nM)
E
kI→C (s-1)
[RNP] (nM)
dCas9
dSpRY
F
kI→O (s-1)
[RNP] (nM)
dCas9
dSpRY
G
kO→I (s-1)
[RNP] (nM)
dCas9
dSpRY
H
TS*
sgRNA2/ +3-bp MM bubble/ 10 mM Mg2+/ 37°C
NTS*
WT
kfast=8.5 min-1 (87%)
kslow=0.10 min-1 (10%)
SpG
kfast=8.9 min-1 (87%)
kslow=0.12 min-1 (10%)
SpRY
kfast=13.2 min-1 (89%)
kslow=0.20 min-1 (8%)
WT
kfast=2.5 min-1 (77%)
kslow=0.03 min-1 (15%)
SpG
kfast=2.8 min-1 (78%)
kslow=0.02 min-1 (14%)
SpRY
kfast=5.8 min-1 (80%)
kslow=0.02 min-1 (13%)
TS*
sgRNA5/ +3-bp MM bubble/ 10 mM Mg2+/ 37°C
NTS*
WT
kfast=20.8 min-1 (87%)
kslow=0.14 min-1 (11%)
SpG
kfast=20.9 min-1 (89%)
kslow=0.34 min-1 (7%)
SpRY
kfast=20.2 min-1 (91%)
kslow=0.26 min-1 (7%)
WT
kfast=14.2 min-1 (79%)
kslow=0.04 min-1 (14%)
SpG
kfast=15.9 min-1 (82%)
kslow=0.04 min-1 (10%)
SpRY
kobs=16.8 min-1
TS*
sgRNA6/ +3-bp MM bubble/ 0.2 mM Mg2+/ 37°C
NTS*
WT
kfast=3.9 min-1 (63%)
kslow=0.05 min-1 (25%)
SpG
kfast=2.7 min-1 (65%)
kslow=0.04 min-1 (20%)
SpRY
kfast=5.7 min-1 (69%)
kslow=0.12 min-1 (20%)
WT
kfast=2.9 min-1 (55%)
kslow=0.03 min-1 (25%)
SpG
kfast=2.2 min-1 (55%)
kslow=0.03 min-1 (24%)
SpRY
kfast=3.8 min-1 (61%)
kslow=0.07 min-1 (19%)
Product (%)
Time (min)

**Fig. S3. Additional AuRBT analysis and DNA cleavage assays (related to Fig. 3). (A-C)** Example traces from time-resolved measurements of equilibrium twist change Δθ for **(A)** 80 nM dCas9 (gray) and **(B)** 100 nM dSpRY (purple). For **(C)**, varied RNP concentrations (4, 8, 20, 100 nM dSpRY) are marked by a double-sided arrow. 250-ms averaged traces are shown in black. The vertical dashed lines (--) and dash-dot lines (-.) indicate the start and end of the flow of different concentrations of RNP into the chamber. For **(B)** and **(C)**, dSpRY:sgRNA3 induces a Δθ baseline shift. **(D)** Fit of average Δθ baseline shift and [RNP] for dSpRY:sgRNA3 binding to Target1 to a binding equation yielded an effective $K_{d,eff}$ = 16 nM. **(E-G)** Transition rate constants as a function of [RNP] with sgRNA3 on Target1 for dCas9 (gray) and dSpRY (purple) and for **(E)** $k_{I\rightarrow C}$ **(F)** $k_{I\rightarrow O}$ **(G)** $k_{O\rightarrow I}$. Solid lines depict the average value across conditions. Error bars were calculated assuming Poisson statistics. **(H)** Time-course analysis of average DNA cleavage products (n=3) for various guide RNAs on dsDNA substrates containing a 3-bp mismatch bubble adjacent to the PAM, presented separately for TS (left) and NTS (right). The average rate constants, with the amplitudes from the observed double-exponential decay are provided in figure legends. The $k_{fast}$ of TS cleavage are also shown in Fig. 3E as a bar graph.

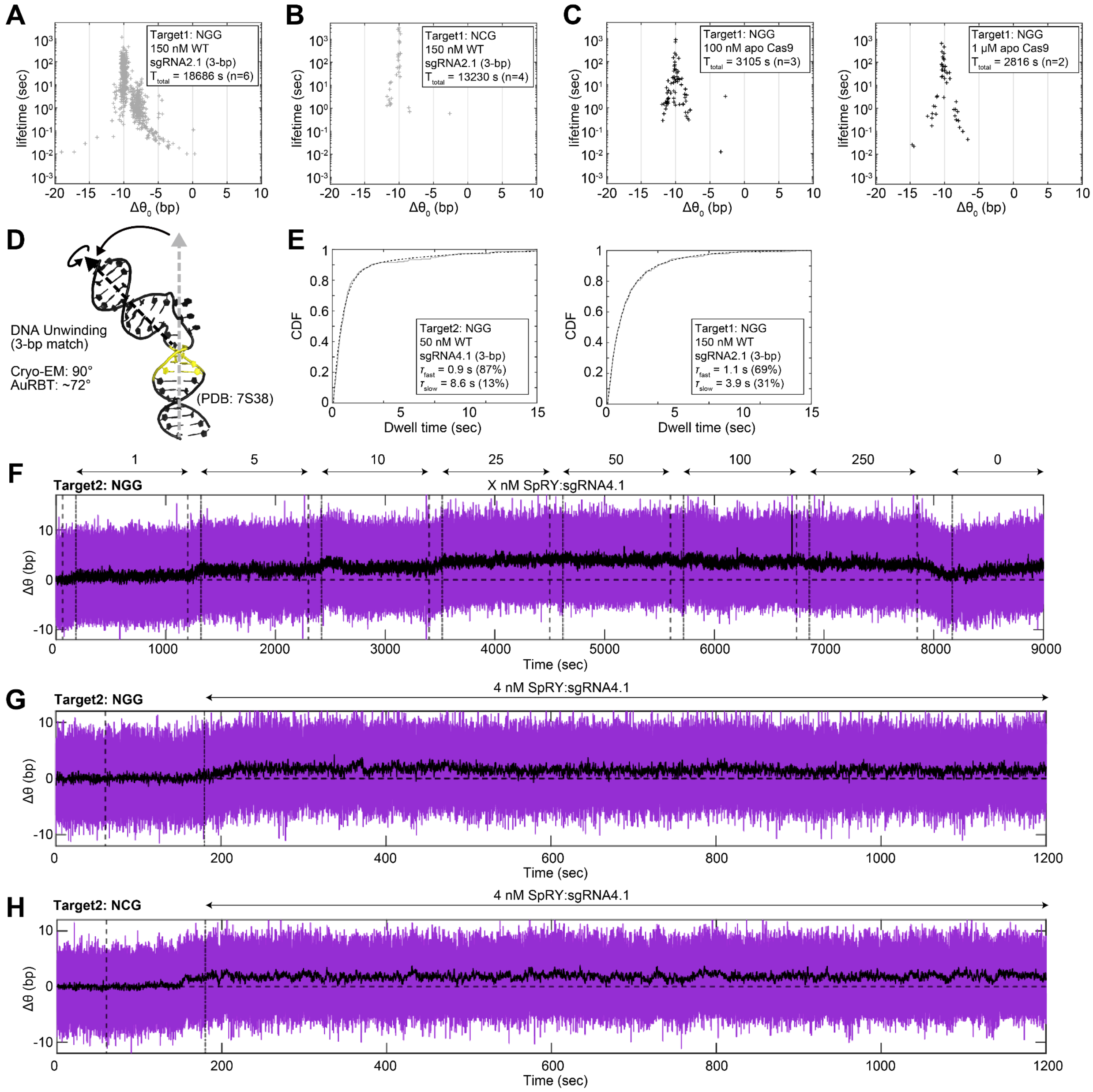


**Fig. S4. Additional AuRBT analysis of early target engagement by WT *Spy*Cas9 and SpRY (related to Fig. 4). (A-C)** Scatter plots illustrating the unwinding lifetime and Δθ for merged Steppi-scored states across all binding events for WT *Spy*Cas9, **(A-B)** with sgRNA2.1 containing only 3-bp match to Target1 flanking **(A)** an NGG site or **(B)** an NCG site, or **(C)** without any guide RNA. The total collection time ($T_{total}$) and number of replicates (n) are provided in the legend. The RNP concentrations are also specified in the legend. **(D)** The degree of DNA unwinding in the Cas9 surveillance complex with 3-bp seed match (PDB: 7S38)[1], measured by an Euler angle analysis[2], is consistent with AuRBT results. **(E)** Cumulative

Distribution Function (CDF) of binding dwell times for (left) sgRNA4.1 on Target2 and (right) sgRNA2.1 on Target1. These distributions are overlaid with a 100-ms left-censored double-exponential model (dashed lines) fit using maximum likelihood estimation in MEMLET[3]. The average lifetimes ($\tau$) for both phases and their amplitudes are provided in the legend. These unwinding lifetimes are comparable to smFRET measurements of bound state dwell times for Cas9 with sgRNA that has 0 or 4-bp matching to the DNA next to an NGG in Singh et al., in which the $\tau_{fast}$ is 0.2-0.4 s ($A_{fast}$~80%) and $\tau_{slow}$ is 2-6 s ($A_{slow}$~20%). **(F-H)** Example traces from time-resolved measurements of equilibrium twist change Δθ for SpRY with sgRNA4.1 on the Target1 sequence flanking an **(F,G)** NGG site or an **(H)** NCG site. For **(F)** varied RNP concentrations (1, 5, 10, 25, 50, 100, 250, 0 nM dSpRY) are marked by a double-sided arrow. 250-ms averaged traces are shown in black. The vertical dashed lines (--) and dash-dot lines (-.) indicate the start and end of the flow of different concentrations of SpRY RNP into the chamber. For **(G-H)**, 4 nM RNP is used.

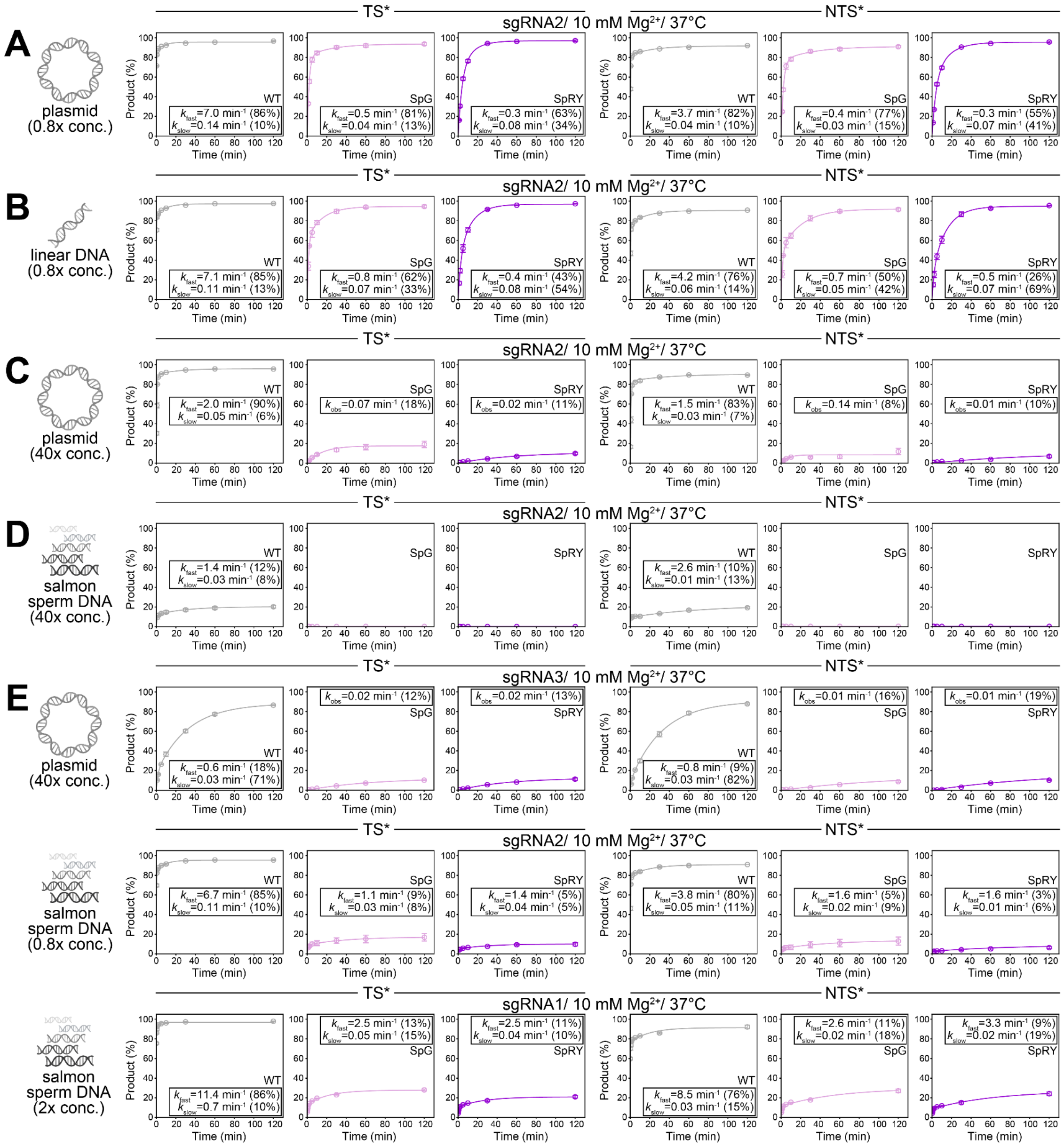


**Fig. S5. DNA cleavage assays with competitors (related to Fig. 5). (A-E)** Time-course analysis of average DNA cleavage products (n=3) for various guide RNAs at different conditions, presented separately for TS (left) and NTS (right). The identities and concentrations of competitors are provided on the left side of each panel. The average rate constants ($k_{obs}$ for a mono-exponential decay model; $k_{fast}$, $k_{slow}$ for a double-exponential decay model), with the amplitudes from the observed exponential decay are provided in figure legends. Note that the TS cleavage data in A, C, D are also shown in Fig. 5C-E and Fig. 6A for direct comparison.

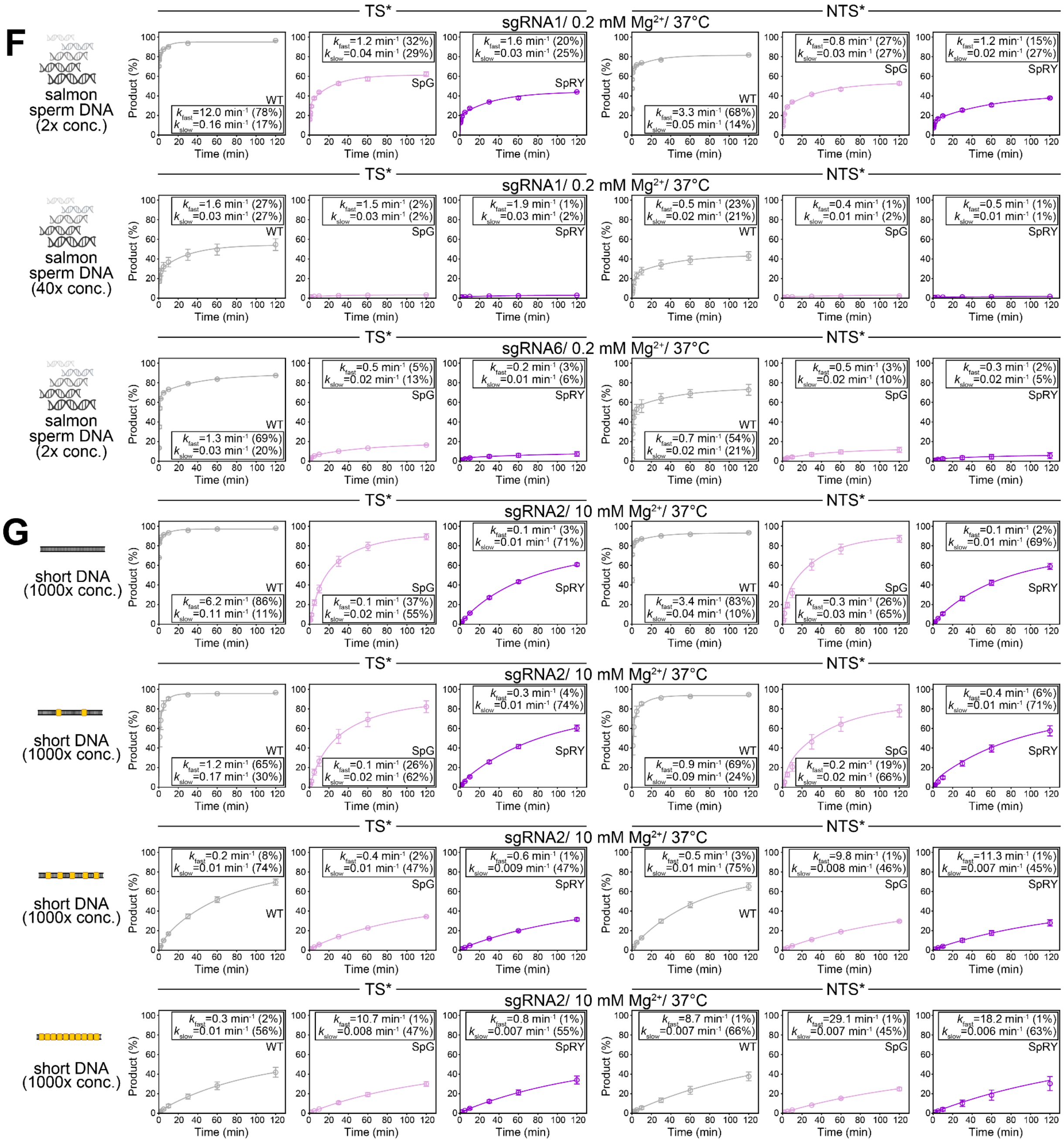


**Fig. S5 (continued). DNA cleavage assays with competitors (related to Fig. 5). (F-G)** Note that the $k_{obs}$ for TS cleavage in panel G, assuming a mono-exponential decay, are presented as a bar graph for direct comparison in Fig. 5F.

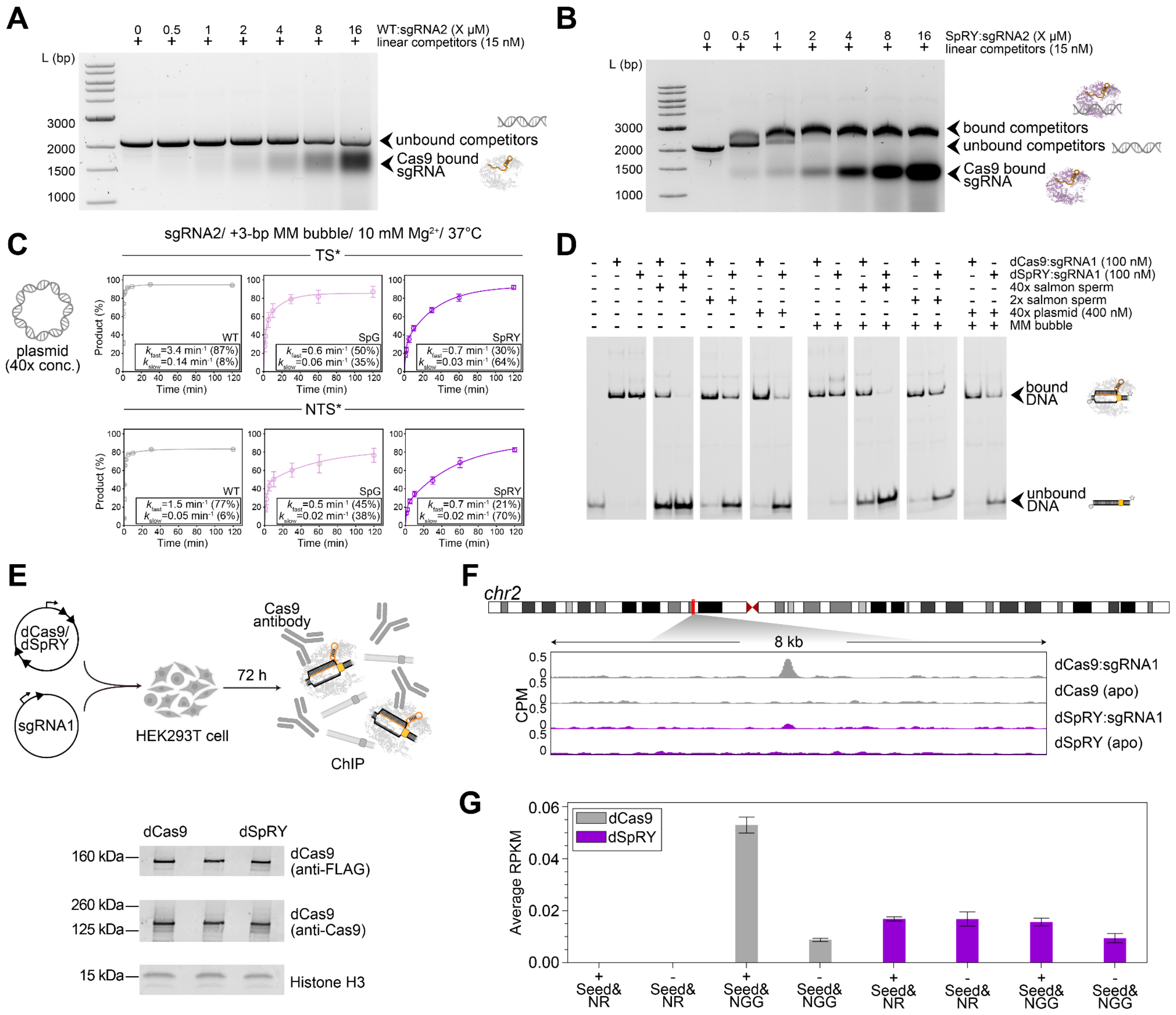


**Fig. S6. Additional data for non-specific DNA interactions (related to Fig. 5-6). (A-B)** EMSA analysis of WT *Spy*Cas9 and SpRY on competitor DNA without target DNA substrate. **(A)** WT *Spy*Cas9 or **(B)** SpRY with sgRNA2 are incubated with a linearized plasmid competitor DNA (2.2-kb, 15 nM) for 1 h at 10 mM $Mg^{2+}$ and 37°C. **(C)** Time-course analysis of average DNA cleavage products (n=3) for sgRNA2 on dsDNA substrates containing a 3-bp mismatch bubble flanking the PAM, presented separately for TS (top) and NTS (bottom), with 40× plasmid competitors. The average rate constants, with the amplitudes from the observed double-exponential decay are provided in figure legends. Note that the TS cleavage data is also shown in Fig. 6A with mono-exponential fitting for SpG and SpRY for direct comparison. **(D)** Additional EMSA analysis of dCas9 and dSpRY with sgRNA1 on the target dsDNA substrate with different competitor identities and concentrations in 0.2 mM $Mg^{2+}$ and 37°C. **(E)** (Top) Schematic overview of ChIP-seq experiments conducted in HEK293T cells. (Bottom) Western blot analysis showing that the anti-Cas9 antibody exhibits similar binding across different Cas9 variants. **(F)** Comparison of dCas9 and dSpRY binding at the on-target site demonstrated by counts-per-million (CPM) normalized ChIP-seq data. **(G)** dCas9 exhibits more specific targeting, exhibiting higher binding to regions containing both the seed and PAM compared to those without them, as measured by ChIP-seq Reads Per Kilobase per Million mapped reads (RPKM). Average RPKM values for regions with and without the specified seed and PAM are plotted for both dCas9 and dSpRY.

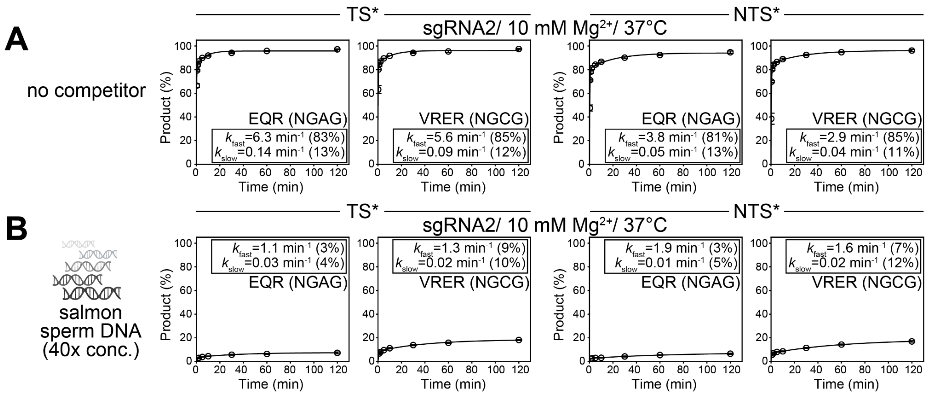


**Fig. S7. DNA cleavage assays using EQR and VRER (related to Fig. 7). (A-B)** Time-course analysis of average DNA cleavage products (n=3) for EQR (with NGAG PAM in the substrate) and VRER (with NGCG PAM in the substrate) with sgRNA2 at 10 mM $Mg^{2+}$ and 37°C, presented separately for TS (left) and NTS (right), under the following conditions: **(A)** without competitor or **(B)** with salmon sperm DNA as competitor. The average rate constants with the amplitudes from the observed exponential decay are provided in figure legends.

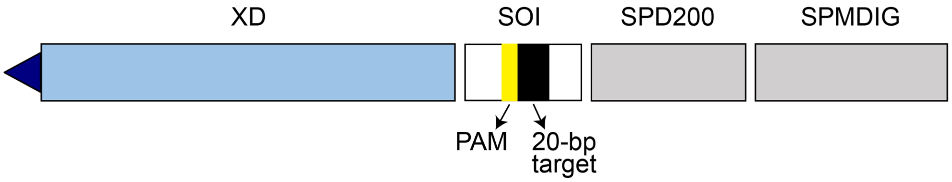


**Fig. S8. Schematic of the DNA tether used in AuRBT (related to Methods).** DNA tethers were constructed by ligating 3 PCR-generated segments (XD, SPD200, SPMDIG) and sequence of interest (SOI) from annealing ssDNA oligonucleotides. PCR primers, templates, restriction enzymes, and ssDNA oligos for making each piece are listed in Table S15 and Table S16. An example of the full tether sequence is provided in Table S17. The top segment (XD) contains one 5'-Fluorescein dT (blue triangle) to make a single attachment with the magnetic bead, and two biotin-modified internal nucleotides for attachment of the rotor bead. The SPMDIG segment contains nucleotides modified with digoxigenin-dUTPs (Roche) to make multiple attachments at the coverslip. The SOI contains an NGG PAM and adjacent target sequence for Cas9 binding. SPD200 and SPMDIG contain no other NGG PAM sites. The NGG is mutated to NCG for no PAM controls.

**Supplemental Tables**

**Table S1. Calculated transition rates from AuRBT experiments with 20-bp match conditions (related to Fig. 2E and Fig. 3A, B).**

| **Target1: dCas9 + sgRNA3 transition rate parameters** | | | | | | |
|---|---|---|---|---|---|---|
| **Concentration [nM]** | $k_{C\rightarrow I}$ [s$^{-1}$] | $k_{I\rightarrow C}$ [s$^{-1}$] | $k_{I\rightarrow O}$ [s$^{-1}$] | $k_{O\rightarrow I}$ [s$^{-1}$] | $k_{C\rightarrow O}$ [s$^{-1}$] | $k_{O\rightarrow C}$ [s$^{-1}$] |
| 0.8 | 0.131 ± 0.040 | 0.126 ± 0.038 | 2.695 ± 0.176 | 0.111 ± 0.007 | – | – |
| 0.8 | 0.046 ± 0.009 | 0.087 ± 0.017 | 2.662 ± 0.092 | 0.170 ± 0.006 | 0.002 ± 0.002 | 0.0002 ± 0.0002 |
| 0.8 | 0.052 ± 0.008 | 0.225 ± 0.034 | 2.478 ± 0.114 | 0.092 ± 0.004 | – | – |
| 4 | 0.180 ± 0.039 | 0.428 ± 0.093 | 3.604 ± 0.271 | 0.152 ± 0.011 | – | – |
| 8 | 0.408 ± 0.057 | 0.210 ± 0.029 | 2.934 ± 0.110 | 0.180 ± 0.007 | – | – |
| 80 | 4.458 ± 1.236 | 0.352 ± 0.098 | 5.228 ± 0.376 | 0.137 ± 0.010 | – | – |
| 80 | 4.156 ± 0.702 | 0.236 ± 0.039 | 2.539 ± 0.129 | 0.228 ± 0.012 | – | – |
| **Average** | – | 0.238 ± 0.119 | 3.163 ± 0.987 | 0.153 ± 0.045 | – | – |

| **Target1: dSpRY + sgRNA3 transition rate parameters** | | | | | | |
|---|---|---|---|---|---|---|
| **Concentration [nM]** | $k_{C\rightarrow I}$ [s$^{-1}$] | $k_{I\rightarrow C}$ [s$^{-1}$] | $k_{I\rightarrow O}$ [s$^{-1}$] | $k_{O\rightarrow I}$ [s$^{-1}$] | $k_{C\rightarrow O}$ [s$^{-1}$] | $k_{O\rightarrow C}$ [s$^{-1}$] |
| 0.8 | 0.007 ± 0.003 | 1.242 ± 0.507 | 6.832 ± 1.189 | 0.371 ± 0.065 | – | – |
| 1.6 | 0.014 ± 0.005 | 1.217 ± 0.430 | 6.997 ± 1.032 | 0.167 ± 0.025 | – | – |
| 4 | 0.010 ± 0.002 | 1.002 ± 0.183 | 2.472 ± 0.287 | 0.129 ± 0.015 | – | – |
| 4 | 0.017 ± 0.002 | 0.382 ± 0.040 | 0.839 ± 0.060 | 0.069 ± 0.005 | – | – |
| 4 | 0.006 ± 0.001 | 0.728 ± 0.121 | 2.346 ± 0.218 | 0.124 ± 0.011 | 0.0002 ± 0.0002 | 0.001 ± 0.001 |
| 4 | 0.006 ± 0.002 | 0.752 ± 0.182 | 2.700 ± 0.346 | 0.129 ± 0.017 | – | – |
| 4 | 0.010 ± 0.003 | 1.238 ± 0.373 | 3.152 ± 0.596 | 0.327 ± 0.061 | – | – |
| 4 | 0.018 ± 0.004 | 0.579 ± 0.123 | 1.262 ± 0.182 | 0.151 ± 0.022 | – | – |

| | | | | | | |
|---|---|---|---|---|---|---|
| 8 | 0.039 ± 0.008 | 1.596 ± 0.313 | 3.623 ± 0.472 | 0.066 ± 0.009 | 0.0015 ± 0.0015 | 0.001 ± 0.001 |
| 8 | 0.017 ± 0.003 | 1.323 ± 0.250 | 4.867 ± 0.480 | 0.057 ± 0.006 | – | 0.0006 ± 0.0006 |
| 20 | 0.026 ± 0.006 | 0.813 ± 0.177 | 3.717 ± 0.379 | 0.085 ± 0.009 | – | – |
| 20 | 0.028 ± 0.004 | 0.815 ± 0.118 | 1.665 ± 0.168 | 0.052 ± 0.005 | 0.0006 ± 0.0006 | 0.0005 ± 0.0005 |
| 40 | 0.049 ± 0.007 | 0.862 ± 0.121 | 5.511 ± 0.305 | 0.122 ± 0.007 | 0.0010 ± 0.0010 | 0.0004 ± 0.0004 |
| 40 | 0.045 ± 0.007 | 0.996 ± 0.164 | 4.037 ± 0.330 | 0.168 ± 0.014 | – | – |
| 100 | 0.101 ± 0.025 | 0.423 ± 0.103 | 2.289 ± 0.239 | 0.178 ± 0.019 | – | – |
| 100 | 0.052 ± 0.013 | 0.560 ± 0.140 | 2.064 ± 0.269 | 0.149 ± 0.019 | – | – |
| 100 | 0.056 ± 0.007 | 0.330 ± 0.039 | 0.934 ± 0.066 | 0.084 ± 0.006 | – | – |
| 100 | 0.070 ± 0.011 | 0.065 ± 0.010 | 0.853 ± 0.036 | 0.390 ± 0.016 | 0.0017 ± 0.0017 | – |
| 250 | 0.022 ± 0.003 | 0.825 ± 0.124 | 2.250 ± 0.205 | 0.092 ± 0.008 | – | 0.0008 ± 0.0008 |
| **Average** | – | 0.829 ± 0.390 | 3.074 ± 1.877 | 0.153 ± 0.101 | – | – |

Errors are calculated assuming Poisson statistics. Averages are unweighted, and errors are standard deviations.

**Table S2. Fits and fit parameters for $k_{C \to I}$ versus RNP concentration (related to Fig. 3A).**

| | **Fit** | **Fit Parameters** |
|---|---|---|
| **dCas9** | $f([dCas9]) = k_{eff} * [dCas9]$ | $k_{eff} = 0.054 \ (0.052, 0.056)$ nM$^{-1}$s$^{-1}$ |
| **dSpRY** | $f([dSpRY]) = \frac{k_{max} * [dSpRY]}{K_{d,init} + [dSpRY]}$ | $K_{d,init} = 14.9 \ (-2.02, 31.81)$ nM<br>$k_{max} = 0.065 \ (0.043, 0.086)$ s$^{-1}$ |

(95% confidence bounds)

**Table S3. The energy values used in the free energy landscapes (related to Fig. 3C).**

| | State | Energy [$k_B$T] |
|---|---|---|
| **dCas9:sgRNA3 on Target1** | $C_{free}$ | 0 |
| | $(C_{free} \leftrightarrow C_{bound})^‡$ | 4.70 |
| | $C_{bound}$ | 2.30 |
| | $(C_{bound} \leftrightarrow I)^‡$ | 5.39 |
| | I | -3.04 |
| | $(I \leftrightarrow O)^‡$ | 2.80 |
| | O | -6.07 |
| **dSpRY:sgRNA3 on Target1** | $C_{free}$ | 0 |
| | $(C_{free} \leftrightarrow C_{bound})^‡$ | 4.70 |
| | $C_{bound}$ | -1.90 |
| | $(C_{bound} \leftrightarrow I)^‡$ | 7.84 |
| | I | 0.65 |
| | $(I \leftrightarrow O)^‡$ | 6.53 |
| | O | -2.35 |

Assuming [RNP] = 100 nM, $k_{on,init}$(dCas9) = $k_{on,init}$(dSpRY) = 0.1 nM$^{-1}$s$^{-1}$, $k_{off,init}$(dCas9) = 100 s$^{-1}$, and the parameters shown in Fig. 3B and Table S1-2, with the arbitrary shift C=7 kT applied to barrier heights for display.

**Table S4. Plasmid vectors used in this study (related to Methods).**

| Internal ID | System | Purpose |
|---|---|---|
| pKMW129 | Human cells | Mammalian expression of Cas9 with EMX1-targeting sgRNA |

| | | |
|---|---|---|
| pKMW130 | Human cells | Mammalian expression of SpG with EMX1-targeting sgRNA |
| pKMW131 | Human cells | Mammalian expression of SpRY with EMX1-targeting sgRNA |
| pKMW124 | Human cells | Mammalian expression of dCas9 |
| pKMW281 | Human cells | Mammalian expression of dSpRY |
| pKMW306 | Human cells | Mammalian expression of EMX1-targeting sgRNA |
| pHS1 | Bacteria | Bacterial expression of WT *Spy*Cas9 |
| pHS23 | Bacteria | Bacterial expression of SpG |
| pHS24 | Bacteria | Bacterial expression of SpRY |
| pHS25 | Bacteria | Bacterial expression of EQR |
| pHS26 | Bacteria | Bacterial expression of VRER |
| pHS16 | Bacteria | Bacterial expression of xPBA |
| pHS9 | Bacteria | Bacterial expression of dCas9 |
| pHS484 | Bacteria | Bacterial expression of dSpRY |
| pHS71 | Bacteria | Bacterial expression of WT *Spy*Cas9-2NLS |
| pHS72 | Bacteria | Bacterial expression of SpG-2NLS |
| pHS73 | Bacteria | Bacterial expression of SpRY-2NLS |
| pGGAselect | Bacteria | Plasmid competitor DNA used in this study. |

**Table S5. U6 promoter and sgRNA sequences used in human cell expression vectors (related to Methods).**

| | **Sequence (5′-3′)** |
|---|---|
| U6 promoter | GAGGGCCTATTTCCCATGATTCCTTCATATTTGCATATACGAT ACAAGGCTGTTAGAGAGATAATTGGAATTAATTTGACTGTAAA CACAAAGATATTAGTACAAAATACGTGACGTAGAAAGTAATAA TTTCTTGGGTAGTTTGCAGTTTTAAAATTATGTTTTAAAATGGA CTATCATATGCTTACCGTAACTTGAAAGTATTTCGATTTCTTG GCTTTATATATCTTGTGGAAAGGACGAAACACC |
| sgRNA (EMX1 spacer) | **GAGTCCGAGCAGAAGAAGAA**GTTTTAGAGCTAGAAATAGCA AGTTAAAATAAGGCTAGTCCGTTATCAACTTGAAAAAGTGGCA CCGAGTCGGTGCTTTTTT |

**Bolded region** represents the 20-bp sgRNA spacer; yellow highlighted region represents the sgRNA scaffold sequence

**Table S6. WT *Spy*Cas9 protein sequences in different expression vectors (related to Methods).**

| | **Protein sequences** |
|---|---|
| WT *Spy*Cas9 in human cell vector | MDYKDHDGDYKDHDIDYKDDDDKMAPKKKRKVGIHGVPAADKK<br>YSIGLDIGTNSVGWAVITDEYKVPSKKFKVLGNTDRHSIKKNLIGA<br>LLFDSGETAEATRLKRTARRRYTRRKNRICYLQEIFSNEMAKVD<br>DSFFHRLEESFLVEEDKKHERHPIFGNIVDEVAYHEKYPTIYHLR<br>KKLVDSTDKADLRLIYLALAHMIKFRGHFLIEGDLNPDNSDVDKL<br>FIQLVQTYNQLFEENPINASGVDAKAILSARLSKSRRLENLIAQLP<br>GEKKNGLFGNLIALSLGLTPNFKSNFDLAEDAKLQLSKDTYDDDL<br>DNLLAQIGDQYADLFLAAKNLSDAILLSDILRVNTEITKAPLSASMI<br>KRYDEHHQDLTLLKALVRQQLPEKYKEIFFDQSKNGYAGYIDGG<br>ASQEEFYKFIKPILEKMDGTEELLVKLNREDLLRKQRTFDNGSIP<br>HQIHLGELHAILRRQEDFYPFLKDNREKIEKILTFRIPYYVGPLAR<br>GNSRFAWMTRKSEETITPWNFEEVVDKGASAQSFIERMTNFDK<br>NLPNEKVLPKHSLLYEYFTVYNELTKVKYVTEGMRKPAFLSGEQ<br>KKAIVDLLFKTNRKVTVKQLKEDYFKKIECFDSVEISGVEDRFNA<br>SLGTYHDLLKIIKDKDFLDNEENEDILEDIVLTLTLFEDREMIEERL<br>KTYAHLFDDKVMKQLKRRRYTGWGRLSRKLINGIRDKQSGKTIL<br>DFLKSDGFANRNFMQLIHDDSLTFKEDIQKAQVSGQGDSLHEHI<br>ANLAGSPAIKKGILQTVKVVDELVKVMGRHKPENIVIEMARENQT<br>TQKGQKNSRERMKRIEEGIKELGSQILKEHPVENTQLQNEKLYL<br>YYLQNGRDMYVDQELDINRLSDYDVDHIVPQSFLKDDSIDNKVL<br>TRSDKNRGKSDNVPSEEVVKKMKNYWRQLLNAKLITQRKFDNL<br>TKAERGGLSELDKAGFIKRQLVETRQITKHVAQILDSRMNTKYDE<br>NDKLIREVKVITLKSKLVSDFRKDFQFYKVREINNYHHAHDAYLN<br>AVVGTALIKKYPKLESEFVYGDYKVYDVRKMIAKSEQEIGKATAK<br>YFFYSNIMNFFKTEITLANGEIRKRPLIETNGETGEIVWDKGRDFA<br>TVRKVLSMPQVNIVKKTEVQTGGFSKESILPKRNSDKLIARKKD<br>WDPKKYGGFDSPTVAYSVLVVAKVEKGKSKKLKSVKELLGITIM<br>ERSSFEKNPIDFLEAKGYKEVKKDLIIKLPKYSLFELENGRKRMLA<br>SAGELQKGNELALPSKYVNFLYLASHYEKLKGSPEDNEQKQLFV<br>EQHKHYLDEIIEQISEFSKRVILADANLDKVLSAYNKHRDKPIREQ<br>AENIIHLFTLTNLGAPAAFKYFDTTIDRKRYTSTKEVLDATLIHQSI<br>TGLYETRIDLSQLGGDKRPAATKKAGQAKKKK |
| WT *Spy*Cas9 in bacterial expression vector | MKSSHHHHHHHHHHGSSMKIEEGKLVIWINGDKGYNGLAEVGK<br>KFEKDTGIKVTVEHPDKLEEKFPQVAATGDGPDIIFWAHDRFGG<br>YAQSGLLAEITPDKAFQDKLYPFTWDAVRYNGKLIAYPIAVEALS<br>LIYNKDLLPNPPKTWEEIPALDKELKAKGKSALMFNLQEPYFTWP<br>LIAADGGYAFKYENGKYDIKDVGVDNAGAKAGLTFLVDLIKNKH |

| | |
|---|---|
| | MNADTDYSIAEAAFNKGETAMTINGPWAWSNIDTSKVNYGVTVL<br>PTFKGQPSKPFVGVLSAGINAASPNKELAKEFLENYLLTDEGLEA<br>VNKDKPLGAVALKSYEEELAKDPRIAATMENAQKGEIMPNIPQM<br>SAFWYAVRTAVINAASGRQTVDEALKDAQTNSSSNNNNNNNNN<br>NLGIEENLYFQSNAMDKKYSIGLDIGTNSVGWAVITDEYKVPSKK<br>FKVLGNTDRHSIKKNLIGALLFDSGETAEATRLKRTARRRYTRRK<br>NRICYLQEIFSNEMAKVDDSFFHRLEESFLVEEDKKHERHPIFGN<br>IVDEVAYHEKYPTIYHLRKKLVDSTDKADLRLIYLALAHMIKFRGH<br>FLIEGDLNPDNSDVDKLFIQLVQTYNQLFEENPINASGVDAKAILS<br>ARLSKSRRLENLIAQLPGEKKNGLFGNLIALSLGLTPNFKSNFDL<br>AEDAKLQLSKDTYDDDLDNLLAQIGDQYADLFLAAKNLSDAILLS<br>DILRVNTEITKAPLSASMIKRYDEHHQDLTLLKALVRQQLPEKYK<br>EIFFDQSKNGYAGYIDGGASQEEFYKFIKPILEKMDGTEELLVKL<br>NREDLLRKQRTFDNGSIPHQIHLGELHAILRRQEDFYPFLKDNRE<br>KIEKILTFRIPYYVGPLARGNSRFAWMTRKSEETITPWNFEEVVD<br>KGASAQSFIERMTNFDKNLPNEKVLPKHSLLYEYFTVYNELTKV<br>KYVTEGMRKPAFLSGEQKKAIVDLLFKTNRKVTVKQLKEDYFKKI<br>ECFDSVEISGVEDRFNASLGTYHDLLKIIKDKDFLDNEENEDILED<br>IVLTLTLFEDREMIEERLKTYAHLFDDKVMKQLKRRRYTGWGRL<br>SRKLINGIRDKQSGKTILDFLKSDGFANRNFMQLIHDDSLTFKEDI<br>QKAQVSGQGDSLHEHIANLAGSPAIKKGILQTVKVVDELVKVMG<br>RHKPENIVIEMARENQTTQKGQKNSRERMKRIEEGIKELGSQILK<br>EHPVENTQLQNEKLYLYYLQNGRDMYVDQELDINRLSDYDVDHI<br>VPQSFLKDDSIDNKVLTRSDKNRGKSDNVPSEEVVKKMKNYWR<br>QLLNAKLITQRKFDNLTKAERGGLSELDKAGFIKRQLVETRQITK<br>HVAQILDSRMNTKYDENDKLIREVKVITLKSKLVSDFRKDFQFYK<br>VREINNYHHAHDAYLNAVVGTALIKKYPKLESEFVYGDYKVYDV<br>RKMIAKSEQEIGKATAKYFFYSNIMNFFKTEITLANGEIRKRPLIET<br>NGETGEIVWDKGRDFATVRKVLSMPQVNIVKKTEVQTGGFSKE<br>SILPKRNSDKLIARKKDWDPKKYGGFDSPTVAYSVLVVAKVEKG<br>KSKKLKSVKELLGITIMERSSFEKNPIDFLEAKGYKEVKKDLIIKLP<br>KYSLFELENGRKRMLASAGELQKGNELALPSKYVNFLYLASHYE<br>KLKGSPEDNEQKQLFVEQHKHYLDEIIEQISEFSKRVILADANLD<br>KVLSAYNKHRDKPIREQAENIIHLFTLTNLGAPAAFKYFDTTIDRK<br>RYTSTKEVLDATLIHQSITGLYETRIDLSQLGGD |

| | |
|---|---|
| WT *Spy*Cas9-2NLS in bacterial expression vector | MKSSHHHHHHHHHHGSSMKIEEGKLVIWINGDKGYNGLAEVGKKFEKDTGIKVTVEHPDKLEEKFPQVAATGDGPDIIFWAHDRFGGYAQSGLLAEITPDKAFQDKLYPFTWDAVRYNGKLIAYPIAVEALSLIYNKDLLPNPPKTWEEIPALDKELKAKGKSALMFNLQEPYFTWPLIAADGGYAFKYENGKYDIKDVGVDNAGAKAGLTFLVDLIKNKHMNADTDYSIAEAAFNKGETAMTINGPWAWSNIDTSKVNYGVTVLPTFKGQPSKPFVGVLSAGINAASPNKELAKEFLENYLLTDEGLEAVNKDKPLGAVALKSYEEELAKDPRIAATMENAQKGEIMPNIPQMSAFWYAVRTAVINAASGRQTVDEALKDAQTNSSSNNNNNNNNNNLGIEENLYFQSNAMDKKYSIGLDIGTNSVGWAVITDEYKVPSKKFKVLGNTDRHSIKKNLIGALLFDSGETAEATRLKRTARRRYTRRKNRICYLQEIFSNEMAKVDDSFFHRLEESFLVEEDKKHERHPIFGNIVDEVAYHEKYPTIYHLRKKLVDSTDKADLRLIYLALAHMIKFRGHFLIEGDLNPDNSDVDKLFIQLVQTYNQLFEENPINASGVDAKAILSARLSKSRRLENLIAQLPGEKKNGLFGNLIALSLGLTPNFKSNFDLAEDAKLQLSKDTYDDDLDNLLAQIGDQYADLFLAAKNLSDAILLSDILRVNTEITKAPLSASMIKRYDEHHQDLTLLKALVRQQLPEKYKEIFFDQSKNGYAGYIDGGASQEEFYKFIKPILEKMDGTEELLVKLNREDLLRKQRTFDNGSIPHQIHLGELHAILRRQEDFYPFLKDNREKIEKILTFRIPYYVGPLARGNSRFAWMTRKSEETITPWNFEEVVDKGASAQSFIERMTNFDKNLPNEKVLPKHSLLYEYFTVYNELTKVKYVTEGMRKPAFLSGEQKKAIVDLLFKTNRKVTVKQLKEDYFKKIECFDSVEISGVEDRFNASLGTYHDLLKIIKDKDFLDNEENEDILEDIVLTLTLFEDREMIEERLKTYAHLFDDKVMKQLKRRRYTGWGRLSRKLINGIRDKQSGKTILDFLKSDGFANRNFMQLIHDDSLTFKEDIQKAQVSGQGDSLHEHIANLAGSPAIKKGILQTVKVVDELVKVMGRHKPENIVIEMARENQTTQKGQKNSRERMKRIEEGIKELGSQILKEHPVENTQLQNEKLYLYYLQNGRDMYVDQELDINRLSDYDVDHIVPQSFLKDDSIDNKVLTRSDKNRGKSDNVPSEEVVKKMKNYWRQLLNAKLITQRKFDNLTKAERGGLSELDKAGFIKRQLVETRQITKHVAQILDSRMNTKYDENDKLIREVKVITLKSKLVSDFRKDFQFYKVREINNYHHAHDAYLNAVVGTALIKKYPKLESEFVYGDYKVYDVRKMIAKSEQEIGKATAKYFFYSNIMNFFKTEITLANGEIRKRPLIETNGETGEIVWDKGRDFATVRKVLSMPQVNIVKKTEVQTGGFSKESILPKRNSDKLIARKKDWDPKKYGGFDSPTVAYSVLVVAKVEKGKSKKLKSVKELLGITIMERSSFEKNPIDFLEAKGYKEVKKDLIIKLPKYSLFELENGRKRMLASAGELQKGNELALPSKYVNFLYLASHYEKLKGSPEDNEQKQLFVEQHKHYLDEIIEQISEFSKRVILADANLDKVLSAYNKHRDKPIREQAENIIHLFTLTNLGAPAAFKYFDTTIDRKRYTSTKEVLDATLIHQSITGLYETRIDLSQLGGDGSPKKKRKVEDPKKKRKVDTG |

3xFLAG tag; NLS sequences; 10 His-tags; MBP; TEV site; Cas9 protein

**Table S7. Protein mutations of Cas9 variants (related to Methods).**

| **Cas9 variants** | **Protein mutations** |
|---|---|
| WT *Spy*Cas9 | |
| SpG | D1135L/S1136W/G1218K/E1219Q/R1335Q/T1337R |
| SpRY | A61R/L1111R/D1135L/S1136W/G1218K/E1219Q/N1317R/A1322R/R1333P/R1335Q/T1337R |
| EQR | D1135E/R1335Q/T1337R |
| VRER | D1135V/G1218R/R1335E/T1337R |
| xPBA | R1333A/R1335A |
| dCas9 | D10A/H840A |
| dSpRY | D10A/A61R/H840A/L1111R/D1135L/S1136W/G1218K/E1219Q/N1317R/A1322R/R1333P/R1335Q/T1337R |

**Table S8. Example of DNA template sequences used for in-vitro transcription (related to Methods).**

| | **Sequences (5′-3′)** |
|---|---|
| sgRNA1 DNA template | TAATACGACTCACTATA**GAGTCCGAGCAGAAGAAGAA**GTTTTAGAGCTAGAAATAGCAAGTTAAAATAAGGCTAGTCCGTTATCAACTTGAAAAAGTGGCACCGAGTCGGTGCTTCG |

Underlined region represents the T7 promoter sequence; **bolded region** represents the 20-bp sgRNA spacer; yellow highlighted region represents the sgRNA scaffold sequence.

**Table S9. Sequences of all the guide RNA used in this study (related to Methods).**

| **sgRNA name** | **Internal ID** | **Sequences (5′-3′)** | **Figures** |
|---|---|---|---|
| sgRNA1 | rHS_77 | **GAGUCCGAGCAGAAGAAGAA**GUUUUAGAGCUAGAAAUAGCAAGUUAAAAUAAGGCUAGUCCGUUAUCAACUUGAAAAAGUGGCACCGAGUCGGUGCUUCG | Figure S2C, S2H, Figure S5E, Figure S5F, Figure S6D |
| sgRNA1$^{AltR}$ | rHS_76 | **mG*mA*mG*UCCGAGCAGAAGAAGAA**GUUUUAGAGCUAGAAAUAGCAAGUUAAAAUAA | Figure 1D, Figure S1E |

| | | | |
|---|---|---|---|
| | | GGCUAGUCCGUUAUCAACUUGAAAAAGUG GCACCGAGUCGG*mU*mG*mC | |
| sgRNA2 | rHS_6 | GG**GACGCAUAAAGAUGAGACAA**GUUUUA GAGCUAGAAAUAGCAAGUUAAAAUAAGGC UAGUCCGUUAUCAACUUGAAAAAGUGGCA CCGAGUCGGUGCUUCG | Figure 1E, F, Figure 3E, Figure 5B-F, Figure 6A, B, Figure S1G-I, Figure S2A, D, Figure S3H, Figure S5A-E, G, Figure S6A-C, Figure S7 |
| sgRNA2.1 | rHS_55 | GG**GACGCAUAAAGAUGAGAGUU**GUUUUA GAGCUAGAAAUAGCAAGUUAAAAUAAGGC UAGUCCGUUAUCAACUUGAAAAAGUGGCA CCGAGUCGGUGCUUCG | Figure S4A, B, E |
| sgRNA3 | rHS_1 | GG**CUGCGUAUUUCUACUCUGUU**GUUUUA GAGCUAGAAAUAGCAAGUUAAAAUAAGGC UAGUCCGUUAUCAACUUGAAAAAGUGGCA CCGAGUCGGUGCUUCG | Figure 2, Figure 3A-C, Figure 4G-4I, Figure S2E, Figure S3A-G, Figure S5E |
| sgRNA4 | rHS_2 | GG**CACACACACACACACACAGG**GUUUUA GAGCUAGAAAUAGCAAGUUAAAAUAAGGC UAGUCCGUUAUCAACUUGAAAAAGUGGCA CCGAGUCGGUGCUUCG | Figure S2F |
| sgRNA4.1 | rHS_5 | GG**CACACACACACACACACCAA**GUUUUAG AGCUAGAAAUAGCAAGUUAAAAUAAGGCU AGUCCGUUAUCAACUUGAAAAAGUGGCAC CGAGUCGGUGCUUCG | Figure 4A-F, Figure S4E-G |
| sgRNA5 | rHS_70 | **GACGCAUAAAGAUGAGACGC**GUUUUAGA GCUAGAAAUAGCAAGUUAAAAUAAGGCUA GUCCGUUAUCAACUUGAAAAAGUGGCACC GAGUCGGUGCUUCG | Figure 3E, Figure S2G, Figure S3H |
| sgRNA6 | rHS_83 | **GAGUGCUAAGGGAACGUUCA**GUUUUAGA GCUAGAAAUAGCAAGUUAAAAUAAGGCUA GUCCGUUAUCAACUUGAAAAAGUGGCACC GAGUCGGUGCUUCG | Figure 3E, Figure S2I, Figure S3H, Figure S5F |
| sgRNA6$^{AltR}$ | rHS_81 | **mG*mA*mG*UGCUAAGGGAACGUUCA**GUU UUAGAGCUAGAAAUAGCAAGUUAAAAUAA GGCUAGUCCGUUAUCAACUUGAAAAAGUG GCACCGAGUCGG*mU*mG*mC | Figure S1E |
| sgRNA7$^{AltR}$ | rHS_82 | **mG*mG*mU*GCUAGCCUUGCGUUCCG**GUU UUAGAGCUAGAAAUAGCAAGUUAAAAUAA GGCUAGUCCGUUAUCAACUUGAAAAAGUG | Figure S1E |

| | | | |
|---|---|---|---|
| | | GCACCGAGUCGG*mU*mG*mC | |
| sgRNA8 | rHS_95 | **GUGAUAAGUGGAAUGCCAUG**GUUUUAGAGCUAGAAAUAGCAAGUUAAAAUAAGGCUAGUCCGUUAUCAACUUGAAAAAGUGGCACCGAGUCGGUGCUUCG | Figure S2B |

AltR represents chemical modifications of IDT guide RNA; m represents 2’-O methylation; * represents phosphorothioate linkage; **bolded region** represents the 20-bp sgRNA spacer; yellow highlighted region represents the sgRNA scaffold sequence.

**Table S10. Sequences of the PCR1 primers for NGS analysis (related to Methods).**

| **Oligonucleotide** | **Sequence (5′-3′)** |
|---|---|
| EMX1 Target Forward Primer | **ACACTCTTTCCCTACACGACGCTCTTCCGATCT**TTTCTCATCTGTGCCCCTCCC |
| EMX1 Target Reverse Primer | **GTGACTGGAGTTCAGACGTGTGCTCTTCCGATCT**GCAGCAAGCAGCACTCTGCC |
| DNMT1 Target Forward Primer | **ACACTCTTTCCCTACACGACGCTCTTCCGATCT**GGAACACGCCCGGTGTCAC |
| DNMT1 Target Reverse Primer | **GTGACTGGAGTTCAGACGTGTGCTCTTCCGATCT**CTGGGGCCGTTTCCCTCAC |
| PCSK9 Target Forward Primer | **ACACTCTTTCCCTACACGACGCTCTTCCGATCT**TCAGCTCCAGGCGGTCCTG |
| PCSK9 Target Reverse Primer | **GTGACTGGAGTTCAGACGTGTGCTCTTCCGATCT**GGCCCGAGAGGAAACAGCAC |

**Bolded region** represents the illumina adapter sequences.

**Table S11. Sequences of oligonucleotides used for ddPCR (related to Methods).**

| **Oligonucleotide** | **Sequence (5′-3′)** |
|---|---|
| EMX1 Target Forward Primer | CCAGAACCGGAGGACAAAGTAC |
| EMX1 Target Reverse Primer | CCACCCTAGTCATTGGAGGTGAC |
| EMX1 Target Probe (FAM) Primer | **/56-FAM/**CTGCTTCGT**/ZEN/**GGCAATGCGCCA |

| | |
|---|---|
| | CC**/3IABkFQ/** |
| EMX1 Reference Forward Primer | GACCACTTGGCCTTCTCCTC |
| EMX1 Reference Reverse Primer | CACTAAACTACAGTGGTGCCTGG |
| EMX1 Reference Probe (HEX) Primer | /5HEX/CCGCCCGCC**/ZEN/**ACCGCAGCCTC **/3IABkFQ/** |

**/56-FAM/** represents 5′-FAM labeling; **/5HEX/** represents 5'-HEX labeling; **/ZEN/** represents internal ZEN quencher; **/3IABkFQ/** represents 3' Iowa Black FQ quencher.

**Table S12. Sequences of the primers used for RT-qPCR (related to Methods).**

| **Oligonucleotide** | **Sequence (5′-3′)** |
|---|---|
| EMX1 sgRNA Forward Primer | GAGTCCGAGCAGAAGAAGAAG |
| EMX1 sgRNA Reverse Primer | CTCGGTGCCACTTTTTCAAG |
| B-actin Forward Primer | ACCTTCTACAATGAGCTGCG |
| B-actin Reverse Primer | CCTGGATAGCAACGTACATGG |

**Table S13. Sequences of all the DNA substrates used in biochemical assays (related to Methods).**

| Internal ID | Description | Sequences (5′-3′) | Figures |
|---|---|---|---|
| HS_1313 | DNA substrate 0-20 bp match to sgRNA1; NTS; Cy5 | /5Cy5/GGAAGGGCCT**GAGTCCGAGCAGAAGAAGA A**GGGCTCCCATCACATCAACCG | Figure S2C, S2H, Figure S5E-F, Figure S6D |
| HS_1339 | DNA substrate 3-20 bp match to sgRNA1 (3-bp mismatch bubble); NTS; Cy5 | /5Cy5/GGAAGGGCCT**GAGTCCGAGCAGAAGAACT T**GGGCTCCCATCACATCAACCG | Figure S6D |
| HS_1276 | DNA substrate 0-20 bp match to sgRNA1; TS; FAM | /56-FAMN/CGGTTGATGTGATGGGAGCCC**TTCTTCTTC TGCTCGGACTC**AGGCCCTTCC | Figure S2C, S2H, Figure S5E-F, Figure S6D |
| HS_33 | DNA substrate 0-20 bp match to sgRNA2; NTS; unlabeled | CGCTCATGCT**GACGCATAAAGATGAGACAA**TGGC GATTACAGTACGTGCG | Figure 4G-I |
| HS_1198 | DNA substrate 0-20 bp match to sgRNA2; NTS; 2AP at position 15 from PAM | CGCTCATGCT**GACGC/i2AmPr/TAAAGATGAGACA A**TGGCGATTACAGTACGTGCG | Figure 1F, Figure S2B |
| HS_284 | DNA substrate 0-20 bp match to sgRNA2; NTS; Cy5 | /5Cy5/CGCTCATGCT**GACGCATAAAGATGAGACAA** TGGCGATTACAGTACGTGCG | Figure 1E, Figure 3E, Figure 5B-F, Figure 6A, B, Figure S1G-I, Figure S2A, S2D, Figure S5A-E, G, Figure S6C |
| HS_500 | DNA substrate 3-20 bp match to sgRNA2 (3-bp mismatch bubble); NTS; Cy5 | /5Cy5/CGCTCATGCT**GACGCATAAAGATGAGAGTT** TGGCGATTACAGTACGTGCG | Figure 3E, Figure S3H |
| HS_34 | DNA substrate 0-20 bp match to sgRNA2; TS; unlabeled | CGCACGTACTGTAATCGCCA**TTGTCTCATCTTTAT GCGTC**AGCATGAGCG | Figure 1F, Figure S2B |
| HS_285 | DNA substrate 0-20 bp match to sgRNA2; TS; FAM | /56-FAMN/CGCACGTACTGTAATCGCCA**TTGTCTCATC TTTATGCGTC**AGCATGAGCG | Figure 1E, Figure 3E, Figure 5B-F, Figure 6A, B, Figure S1G-I, |

| | | | |
|---|---|---|---|
| | | | Figure S2A, S2D, Figure S3H, Figure S5A-E, G, Figure S6C |
| HS_459 | DNA substrate 0-20 bp match to sgRNA2; NTS; Cy5; NGAG PAM | /5Cy5/CGCTCATGCT**GACGCATAAAGATGAGACAA** TGAGGATTACAGTACGTGCG | Figure S7 |
| HS_360 | DNA substrate 0-20 bp match to sgRNA2; TS; FAM; NGAG PAM | /56-FAMN/CGCACGTACTGTAATCCTCA**TTGTCTCATCT TTATGCGTC**AGCATGAGCG | Figure S7 |
| HS_461 | DNA substrate 0-20 bp match to sgRNA2; NTS; Cy5; NGCG PAM | /5Cy5/CGCTCATGCT**GACGCATAAAGATGAGACAA** TGCGGATTACAGTACGTGCG | Figure S7 |
| HS_362 | DNA substrate 0-20 bp match to sgRNA2; TS; FAM; NGCG PAM | /56-FAMN/CGCACGTACTGTAATCCGCA**TTGTCTCATC TTTATGCGTC**AGCATGAGCG | Figure S7 |
| HS_37 | DNA substrate 0-20 bp match to sgRNA3; NTS; Cy5 | /5Cy5/CGCTCATGCT**CTGCGTATTTCTACTCTGTT**T GGCGATTACAGTACGTGCG | Figure S2E, Figure S5E |
| HS_38 | DNA substrate 0-20 bp match to sgRNA3; TS; FAM | /56-FAMN/CGCACGTACTGTAATCGCCA**AACAGAGTAG AAATACGCAG**AGCATGAGCG | Figure S2E, Figure S5E |
| HS_451 | DNA substrate 0-20 bp match to sgRNA4; NTS; Cy5 | /5Cy5/CGCTCATGCT**CACACACACACACACACAG G**TGGCGATTACAGTACGTGCG | Figure S2F |
| HS_452 | DNA substrate 0-20 bp match to sgRNA4; TS; FAM | /56-FAMN/CGCACGTACTGTAATCGCCA**CCTGTGTGTG TGTGTGTGTG**AGCATGAGCG | Figure S2F |
| HS_1237 | DNA substrate 0-20 bp match to sgRNA5; NTS; Cy5 | /5Cy5/CGCTCATGCT**GACGCATAAAGATGAGACG C**TGGCGATTACAGTACGTGCG | Figure 3E, Figure S2G |
| HS_1338 | DNA substrate 3-20 bp match to sgRNA5 (3-bp mismatch bubble); NTS; Cy5 | /5Cy5/CGCTCATGCT**GACGCATAAAGATGAGAGC G**TGGCGATTACAGTACGTGCG | Figure 3E, Figure S3H |
| HS_1238 | DNA substrate 0-20 bp match to sgRNA5; TS; | /56-FAMN/CGCACGTACTGTAATCGCCA**GCGTCTCATC** | Figure 3E, Figure S2G, |

| | | | |
|---|---|---|---|
| | FAM | **TTTATGCGTC**AGCATGAGCG | Figure S3H |
| HS_1316 | DNA substrate 0-20 bp match to sgRNA6; NTS; Cy5 | /5Cy5/ATAAGTGGCA**GAGTGCTAAGGGAACGTTCA** CGGAGACTGAACACTCCTCA | Figure 3E, Figure S2I, Figure S5F |
| HS_1342 | DNA substrate 3-20 bp match to sgRNA6 (3-bp mismatch bubble); NTS; Cy5 | /5Cy5/ATAAGTGGCA**GAGTGCTAAGGGAACGTAGT** CGGAGACTGAACACTCCTCA | Figure 3E, Figure S3H |
| HS_1278 | DNA substrate 0-20 bp match to sgRNA6; TS; FAM | /56-FAMN/TGAGGAGTGTTCAGTCTCCG**TGAACGTTCC CTTAGCACTC**TGCCACTTAT | Figure 3E, Figure S2I, Figure S3H, Figure S5F, |
| HS_721 | DNA competitor contains 10 NGG; strand 1 | CAATGGCAATGGCAATGGCAATGGCAATGGCAAT GGCAATGGCAATGGCAATGGCAATGG | Figure 5F, Figure S5G |
| HS_722 | DNA competitor contains 10 NGG; strand 2 | CCATTGCCATTGCCATTGCCATTGCCATTGCCATT GCCATTGCCATTGCCATTGCCATTG | Figure 5F, Figure S5G |
| HS_723 | DNA competitor contains 5 NGG; strand 1 | CAATGGCAATTTCAATGGCAATTTCAATGGCAATTT CAATGGCAATTTCAATGGCAATTT | Figure 5F, Figure S5G |
| HS_724 | DNA competitor contains 5 NGG; strand 2 | AAATTGCCATTGAAATTGCCATTGAAATTGCCATTG AAATTGCCATTGAAATTGCCATTG | Figure 5F, Figure S5G |
| HS_775 | DNA competitor contains 2 NGG; strand 1 | CAATTTCAATTTCAATGGCAATTTCAATTTCAATTT CAATGGCAATTTCAATTTCAATTT | Figure 5F, Figure S5G |
| HS_776 | DNA competitor contains 2 NGG; strand 2 | AAATTGAAATTGAAATTGCCATTGAAATTGAAATTG AAATTGCCATTGAAATTGAAATTG | Figure 5F, Figure S5G |
| HS_727 | DNA competitor contains 0 NGG; strand 1 | CAATTTCAATTTCAATTTCAATTTCAATTTCAATTTC AATTTCAATTTCAATTTCAATTT | Figure 5F, Figure S5G |
| HS_728 | DNA competitor contains 0 NGG; strand 2 | AAATTGAAATTGAAATTGAAATTGAAATTGAAATTG AAATTGAAATTGAAATTGAAATTG | Figure 5F, Figure S5G |

/56-FAMN/ represents 5′-FAM labeling; /5Cy5/ represents 5′-Cy5 labeling; /i2AmPr/ represents internal 2AP labeling; **bolded region** represents the 20-bp sgRNA spacer; yellow highlighted region represents the NGG PAM for WT *Spy*Cas9, NGAG for EQR, NGCG for VRER

**Table S14. Sequences of the plasmid competitor (related to Methods).**

| | **Sequences (5′-3′)** |
|---|---|
| pGGAselect | CGAAAAATCAATAATCAGACAACAAGATGTGCGAACTCGATATTTT<br>ACACGACTCTCTTTACCAATTCTGCCCCGAATTACACTTAAAACGA<br>CTCAACAGCTTAACGTTGGCTTGCCACGCATTACTTGACTGTAAAA<br>CTCTCACTCTTACCGAACTTGGCCGTAACCTGCCAACCAAAGCGA<br>GAACAAAACATAACATCAAACGAATCGACCGATTGTTAGGTAATCG<br>TCACCTGCAGGAAGGTTTAAACGCATTTAGGTGACACTATAGAAGT<br>GTGTATCGCTCGAGGGATCCGAATTCGAAGTCTTGGTACGGAGCG<br>AGACCGGAGCGAGACGGGAGTCGTCTTCGCTTTCCAGATCTGATA<br>ACTTGTGAAGACGACCATCGTCTCACCATGGTCTCACCATTCCTGT<br>AGACTTCTTAATTAAGACGTCAGAATTCTCGAGGCGGCCGCATGT<br>GAGTCTCCCTATAGTGAGTCGTATTAATTTCGCGGGCGGAACCCC<br>TATTTGTTTATTTTTCTAAATACATTCAAATATGTATCCGCTCATGAG<br>TAGCACCAGGCGTTTAAGGGCACCAATAACTGCCTTAAAAAAATTA<br>CGCCCCGCCCTGCCACTCATCGCAGTACTGTTGTAATTCATTAAG<br>CATTCTGCCGACATGGAAGCCATCACAAACGGCATGATGAACCTG<br>AATCGCCAGCGGCATCAGCACCTTGTCGCCTTGCGTATAATATTTG<br>CCCATGGTGAAAACGGGGGCGAAGAAGTTGTCCATATTGGCCACG<br>TTTAAATCAAAACTGGTGAAACTCACCCAGGGATTGGCTGAGACAA<br>AAAACATATTCTCAATAAACCCTTTAGGGAAATAGGCCAGGTTTTC<br>ACCGTAACACGCCACATCTTGCGAATATATGTGTAGAAACTGCCG<br>GAAATCGTCGTGGTATTCACTCCAGAGCGATGAAAACGTTTCAGTT<br>TGCTCATGGAAAACGGTGTAACAAGGGTGAACACTATCCCATATCA<br>CCAGCTCACCGTCTTTCATTGCCATACGAAATTCCGGATGAGCATT<br>CATCAGGCGGGCAAGAATGTGAATAAAGGCCGGATAAAACTTGTG<br>CTTATTTTTCTTTACGGTCTTTAAAAAGGCCGTAATATCCAGCTGAA<br>CGGTCTGGTTATAGGTACATTGAGCAACTGACTGAAATGCCTCAAA<br>ATGTTCTTTACGATGCCATTGGGATATATCAACGGTGGTATATCCA<br>GTGATTTTTTTCTCCATTTTAGCTTCCTTAGCTCCTGAAAATCTCGA<br>TAACTCAAAAAATACGCCCGGTAGTGATCTTATTTCATTATGGTGA<br>AAGTTGGAACCTCTTACGTGCCGATCAAAGTCTCATTTTCGCCAAA<br>AGTTGTCATGACCAAAATCCCTTAACGTGAGTTTTCGTTCCACTGA<br>GCGTCAGACCCCGTAGAAAAGATCAAAGGATCTTCTTGAGATCCTT<br>TTTTTCTGCGCGTAATCTGCTGCTTGCAAACAAAAAAACCACCGCT<br>ACCAGCGGTGGTTTGTTTGCCGGATCAAGAGCTACCAACTCTTTTT<br>CCGAAGGTAACTGGCTTCAGCAGAGCGCAGATACCAAATACTGTT<br>CTTCTAGTGTAGCCGTAGTTAGGCCACCACTTCAAGAACTCTGTAG<br>CACCGCCTACATACCTCGCTCTGCTAATCCTGTTACCAGTGGCTG<br>CTGCCAGTGGCGATAAGTCGTGTCTTACCGGGTTGGACTCAAGAC |

| | |
|---|---|
| | GATAGTTACCGGATAAGGCGCAGCGGTCGGGCTGAACGGGGGGT TCGTGCACACAGCCCAGCTTGGAGCGAACGACCTACACCGAACTG AGATACCTACAGCGTGAGCTATGAGAAAGCGCCACGCTTCCCGAA GGGAGAAAGGCGGACAGGTATCCGGTAAGCGGCAGGGTCGGAAC AGGAGAGCGCACGAGGGAGCTTCCAGGGGGAAACGCCTGGTATC TTTATAGTCCTGTCGGGTTTCGCCACCTCTGACTTGAGCGTCGATT TTTGTGATGCTCGTCAGGGGGGCGGAGCCTATGGAAAAACGCCA GCAATGCGGCCTTTTTACGGTTCCTGGCCTTTTGCTGGCCTTTTGC TCACATGTTCTTTCCTGCGTTATCCCCTGATTCTGTGGATAACCGT ATTACCGCCTTTGAGTGAGCTGATACCGCTCGCCGCAGCCGAACG ACCGAGCGCAGCGAGTCAGTGAGCGAGGAAGC |

**Table S15. PCR primers and templates used for DNA tether construction for AuRBT experiments (related to Methods).**

| **Name** | **Primer sequence (5′-3′)** | **Length (bps)** | **Template** | **Digest** |
|---|---|---|---|---|
| XD | FW primer: gaagggtctcaTGACTCACTA**T**AGGGCGAAT**T**GGAGCTCCACCGCG<br><br>RW primer: /5FluorT/CACTAAAGGGAACAAAAGCTGGTAC | 4140 | pFO-SE2 | BsaI |
| SPD200 | FW primer: gaagggtctcaCACTATTAAGTCGAAACAGTCGAGCGTGAAC<br><br>RW primer: gaagggtctcaGCTCGCGACGTTGCGCTACGTCAT | 212 | pII-lam1 | BsaI |
| SPMDIG | FW primer: GCGCAGCACGCAGATAAATTC<br><br>RW primer: gatcggtctccAGTGCAAACGTCTGCGTCGCTG | 308 | pII-lam1 | BsaI |

Internal biotin-modified nucleotides (/iBiodT/) are colored red; /5FluorT/ represents 5'-Fluorescein dT.

**Table S16. DNA oligonucleotides used to incorporate sequence of interest (SOI) into DNA tether for AuRBT experiments (related to Methods).**

| Name | Target strand (5′-3′) | Nontarget strand (5′-3′) | Length (bps) |
|---|---|---|---|
| Target1: NGG | /5Phos/gtcaTTGTCGCGAAGTGCAGCGAGATCGCGTTTGTACTCCA**AACAGAGTAGAAATACGCAG**AGCAGCGATTTCTCGTGTGCAGACA | /5Phos/gagcTGTCTGCACACGAGAAATCGCTGCT**CTGCGTATTTCTACTCTGTT**TGGAGTACAAACGCGATCTCGCTGCACTTCGCGACAA | 86 |
| Target1: NCG | /5Phos/gtcaTTGTCGCGAAGTGCAGCGAGATCGCGTTTGTACTCGA**AACAGAGTAGAAATACGCAG**AGCAGCGATTTCTCGTGTGCAGACA | /5Phos/gagcTGTCTGCACACGAGAAATCGCTGCT**CTGCGTATTTCTACTCTGTT**TCGAGTACAAACGCGATCTCGCTGCACTTCGCGACAA | 86 |
| Target2: NGG | /5Phos/gtcaTTGTCGCGAAGTGCAGCGAGATCGCGTTTGTACTCCA**TTGTCTCATCTTTATGCGTC**AGCAGCGATTTCTCGTGTGCAGACA | /5Phos/gagcTGTCTGCACACGAGAAATCGCTGCT**GACGCATAAAGATGAGACAA**TGGAGTACAAACGCGATCTCGCTGCACTTCGCGACAA | 86 |
| Target2: NCG | /5Phos/gtcaTTGTCGCGAAGTGCAGCGAGATCGCGTTTGTACTCGA**TTGTCTCATCTTTATGCGTC**AGCAGCGATTTCTCGTGTGCAGACA | /5Phos/gagcTGTCTGCACACGAGAAATCGCTGCT**GACGCATAAAGATGAGACAA**TCGAGTACAAACGCGATCTCGCTGCACTTCGCGACAA | 86 |

/5Phos/ represents 5’-phosphorylation; **bolded region** represents the 20-bp sgRNA spacer;

yellow highlighted region represents the PAM.

**Table S17. Example DNA tether used for AuRBT experiments (related to Methods).**

| | Target strand (5’-3’) |
|---|---|
| Target2: NGG | /5FluorT/CACTAAAGGGAACAAAAGCTGGTACCGGGCCCCCCCTCGAGCGGTACCCCACTTACCCACCCCGGAAATTTGAGTTATAAACGTTGTTTGAGCTTTACCTAGTCTTGGTCGATCAAAAGTTCTGGTACCTTTTCACCATGTCTCCCCCCTTATTCATATAAAAAGAAGCGTATAATCGCACAGTATAACGCTCCTCTGATATATGATCTAGACCCAAGTAATGAGTTACGAATCTGGGAGGTCATCCTCCTCTTCCGAGAGTACACGGCCACCAACGCTAAAAGAAGAACCTAATGGTAAAATAGCTTGGGAAGAAAGTGTCAAAAAATCTAGGGAAAATAACGAAAATGACAGCACTCTCTTGAGGCGAAAGCTAGGTGAGACTCGAAAAGCAATTGAAACTGGAGGATCATCGAGAAATAAACTTTCTGCTTTGACACCCTTGAAAAAAGTGGTTGACGAGAGGAAGGATTCGGTACAACCACAGGTCCCTTCCATGGGTTTTACTTATTCTTTGCCTAATTTGAAGACTTTAAACAGTTTTTCAGATGCTGAGCAAGCACGTATAATGCAAGATTATCTATCCAGGGGGGTAAATCAAGGCAACAGTAATAATTATGTAGACCCACTATATCGGCAATTAAATCCAACTATGGGTAGTAGCAGGAACAGGC |

| | |
|---|---|
| | CTGTTTGGAGTTTAAATCAGCCGTTACCGCATGTATTGGATCGAGGCTTGGC<br>AGCAAAGATGATACAAAAGAATATGGATGCAAGGTCCCGCGCATCATCGAGA<br>CGAGGGTCGACCGATATTTCAAGGGGGGGTTCTACTACGTCAGTGAAAGAC<br>TGGAAAAGGCTCCTTAGAGGTGCAGCACCGGGTAAAAAGCTTGGTGACATC<br>GAAGCTCAAACGCAACGCGATAATACTGTTGGTGCAGATGTGAAACCTACTA<br>AGTTAGAGCCTGAAAACCCACAAAAGCCCTCTAACACGCATATTGAGAATGT<br>TTCACGTAAGAAAAAGCGTACTTCGCATAATGTCAATTTTTCATTAGGCGATG<br>AAAGCTACGCATCCTCCATAGCCGATGCAGAATCCAGAAAATTAAAGAACAT<br>GCAAACCCTCGATGGTTCTACTCCGGTTTATACGAAGCTTCCTGAAGAACTT<br>ATTGAAGAGGAAAATAAAAGTACGAGTGCATTAGATGGTAATGAAATTGGTG<br>CCTCAGAAGATGAAGACGCGGATATAATGACATTTCCTAACTTTTGGGCAAA<br>AATTCGCTATCATATGCGAGAACCGTTTGCGGAGTTTCTCGGGACACTAGTT<br>CTTGTCATTTTTGGTGTTGGTGGTAATCTTCAAGCAACTGTAACAAAAGGTAG<br>TGGTGGTTCCTATGAATCCCTATCATTTGCATGGGGGTTCGGTTGTATGCTT<br>GGTGTTTACGTCGCAGGCGGTATTAGTGGTGGTCATATTAACCCTGCTGTTA<br>CGATTTCAATGGCAATTTTTCGAAAATTCCCCTGGAAAAAGGTGCCCGTATAT<br>ATTGTTGCTCAGATTATCGGTGCATATTTTGGAGGAGCTATGGCTTATGGTTA<br>TTTTTGGAGCTCTATCACAGAATTTGAGGGAGGTCCGCACATAAGAACAACG<br>GCGACCGGTGCGTGTTTGTTTACTGATCCAAAGTCTTACGTCACGTGGAGAA<br>ATGCCTTCTTTGACGAATTCATAGGAGCCTCTATACTTGTGGGTTGTTTGATG<br>GCGCTATTGGATGATAGTAATGCTCCACCTGGCAATGGTATGACCGCATTAA<br>TTATTGGATTCTTAGTCGCTGCAATTGGTATGGCCCTTGGATATCAAACAAGT<br>TTCACAATCAATCCTGCAAGAGATCTCGGTCCTCGCATATTTGCTTCCATGAT<br>TGGCTATGGTCCACATGCTTTTCATCTCACACATTGGTGGTGGACATGGGGA<br>GCCTGGGGTGGTCCAATTGCCGGCGGTATTGCTGGAGCACTCATATATGAC<br>ATTTTCATTTTTACTGGATGCGAATCCCCAGTCAACTACCCAGACAACGGTTA<br>TATTGAGAATAGGGTAGGCTGCAGCCCGGGGGATCCACTAGTTCTAGAGCG<br>GCCGCCGGTACCCCACTTACCCACCCCGGAAATTTGAGTTATAAACGTTGTT<br>TGAGCTTTACCTAGTCTTGGTCGATCAAAAGTTCTGGTACCTTTTCACCATGT<br>CTCCCCCCTTATTCATATAAAAAGAAGCGTATAATCGCACAGTATAACGCTCC<br>TCTGATATATGATCTAGACCCAAGTAATGAGTTACGAATCTGGGAGGTCATC<br>CTCCTCTTCCGAGAGTACACGGCCACCAACGCTAAAAGAAGAACCTAATGGT<br>AAAATAGCTTGGGAAGAAAGTGTCAAAAAATCTAGGGAAAATAACGAAAATG<br>ACAGCACTCTCTTGAGGCGAAAGCTAGGTGAGACTCGAAAAGCAATTGAAAC<br>TGGAGGATCATCGAGAAATAAACTTTCTGCTTTGACACCCTTGAAAAAAGTG<br>GTTGACGAGAGGAAGGATTCGGTACAACCACAGGTCCCTTCCATGGGTTTTA<br>CTTATTCTTTGCCTAATTTGAAGACTTTAAACAGTTTTTCAGATGCTGAGCAA<br>GCACGTATAATGCAAGATTATCTATCCAGGGGGGTAAATCAAGGCAACAGTA<br>ATAATTATGTAGACCCACTATATCGGCAATTAAATCCAACTATGGGTAGTAGC<br>AGGAACAGGCCTGTTTGGAGTTTAAATCAGCCGTTACCGCATGTATTGGATC<br>GAGGCTTGGCAGCAAAGATGATACAAAAGAATATGGATGCAAGGTCCCGCG<br>CATCATCGAGACGAGGGTCGACCGATATTTCAAGGGGGGGTTCTACTACGT<br>CAGTGAAAGACTGGAAAAGGCTCCTTAGAGGTGCAGCACCGGGTAAAAAGC<br>TTGGTGACATCGAAGCTCAAACGCAACGCGATAATACTGTTGGTGCAGATGT<br>GAAACCTACTAAGTTAGAGCCTGAAAACCCACAAAAGCCCTCTAACACGCAT<br>ATTGAGAATGTTTCACGTAAGAAAAAGCGTACTTCGCATAATGTCAATTTTTC<br>ATTAGGCGATGAAAGCTACGCATCCTCCATAGCCGATGCAGAATCCAGAAAA<br>TTAAAGAACATGCAAACCCTCGATGGTTCTACTCCGGTTTATACGAAGCTTCC<br>TGAAGAACTTATTGAAGAGGAAAATAAAAGTACGAGTGCATTAGATGGTAAT<br>GAAATTGGTGCCTCAGAAGATGAAGACGCGGATATAATGACATTTCCTAACT |

| | |
|---|---|
| | TTTGGGCAAAAATTCGCTATCATATGCGAGAACCGTTTGCGGAGTTTCTCGG<br>GACACTAGTTCTTGTCATTTTTGGTGTTGGTGGTAATCTTCAAGCAACTGTAA<br>CAAAAGGTAGTGGTGGTTCCTATGAATCCCTATCATTTGCATGGGGGTTCGG<br>TTGTATGCTTGGTGTTTACGTCGCAGGCGGTATTAGTGGTGGTCATATTAAC<br>CCTGCTGTTACGATTTCAATGGCAATTTTTCGAAAATTCCCCTGGAAAAAGGT<br>GCCCGTATATATTGTTGCTCAGATTATCGGTGCATATTTTGGAGGAGCTATG<br>GCTTATGGTTATTTTTGGAGCTCTATCACAGAATTTGAGGGAGGTCCGCACA<br>TAAGAACAACGGCGACCGGTGCGTGTTTGTTTACTGATCCAAAGTCTTACGT<br>CACGTGGAGAAATGCCTTCTTTGACGAATTCATAGGAGCCTCTATACTTGTG<br>GGTTGTTTGATGGCGCTATTGGATGATAGTAATGCTCCACCTGGCAATGGTA<br>TGACCGCATTAATTATTGGATTCTTAGTCGCTGCAATTGGTATGGCCCTTGGA<br>TATCAAACAAGTTTCACAATCAATCCTGCAAGAGATCTCGGTCCTCGCATATT<br>TGCTTCCATGATTGGCTATGGTCCACATGCTTTTCATCTCACACATTGGTGGT<br>GGACATGGGGAGCCTGGGGTGGTCCAATTGCCGGCGGTATTGCTGGAGCA<br>CTCATATATGACATTTTCATTTTTACTGGATGCGAATCCCCAGTCAACTACCC<br>AGACAACGGTTATATTGAGAATAGGGTAGGGGCCGCCACCGCGGTGGAGCT<br>CC**A**ATTCGCCCT**A**TAGTGAGTCATTGTCGCGAAGTGCAGCGAGATCGCGTTT<br>GTACTCCA**TTGTCTCATCTTTATGCGTC**AGCAGCGATTTCTCGTGTGCAGAC<br>AGCTCGCGACGTTGCGCTACGTCATCTAGTAGTGCGAACTGACTCTCAGTAT<br>TCGACAAATCTTGACGCTGCTGACTGATTTTCTCATTTGCTCATGCATGCTTC<br>TCTAGCGTCAGTAGTTCTGACTGAAGCGTCAGCAGCTCAGCATGAGCACTGT<br>CTAGATGACGATCGCTCGCAGACACGTTCACGCTCGACTGTTTCGACTTAAT<br>AGTGCAAACGTCTGCGTCGCTGCACTTTTTACGTGCGACATACTGTCAGTCG<br>CGCTCTCTTCGCACTCACTCGAGCTGAACTTCAGCAGTGCGATACAGTTCGC<br>TCGAAGCTGCTCTTCAGCTGCTCAGCTCTTTTTCTGCTCTGACATGACGTTAT<br>TCAGCGCGAGCGTATTATCGCGATACTGTTCGTTCACGCGCTGTTCAGTGCG<br>TTGCAGTTCTGCGTGCAGCTCAGTCAGCACTTCACTTTTCGCATCAATGATT<br>GACTGTTTTGCACGTTGCTGCTGTGCGAATTTATCTGCGTGCTGCGC |

/5FluorT/ represents 5'-Fluorescein dT; **A** represents adenine nucleotides paired with biotin-modified thymines, where the streptavidin-coated gold rotor bead could be attached; **bolded region** represents the 20-bp sgRNA spacer; yellow highlighted region represents the PAM; cyan highlighted region represents the segment with multiple incorporated dUTP-digoxigenin (Roche).

### Supplemental Notes

#### Note S1. A kinetic model of early target engagement by dCas9 and dSpRY

Our kinetic model (Fig. 3B) integrates experimental findings with prior Cas9 kinetic studies[1,4-7] to explain differences between dCas9 and dSpRY observed in AuRBT experiments. Analysis of AuRBT yields measurements of four transition rates for each enzyme and [RNP] condition: $k_{C \to I}$, $k_{I \to C}$, $k_{I \to O}$, and $k_{O \to I}$ . Three of these transition rates were found to be approximately [RNP]-independent as expected for the model (Fig. S3E-G), and these measured rates correspond directly to model parameters: $k_{I \to C}$, $k_{I \to O}$, and $k_{O \to I}$ . The remaining measured rate $k_{C \to I}$ varies with [RNP], which was modeled by separating the directly observed C state into the substates $C_{free}$ and $C_{bound}$. Our measurements of $k_{C \to I}$ versus [RNP] (Fig. 3A) are modeled by considering a two-step process (Fig. 3B):

1. **Initial DNA binding ($C_{free} \leftrightarrow C_{bound}$)**: This step is characterized by the equilibrium dissociation constant $K_{d,init}$ which can be expressed as:

$$K_{d,init} = \frac{k_{off,init}}{k_{on,init}}$$

   where $k_{on,init}$ is the association rate of initial binding, and $k_{off,init}$ is the dissociation rate.

2. **Conformational transition ($C_{bound} \leftrightarrow I$)**: This step involves DNA unwinding to form the *R*-loop seed intermediate. The forward and reverse rates for the transition are $k_{open}$ and $k_{close}$, respectively (with $k_{close}$ equal to $k_{I \to C}$).

Every scored dwell in C must begin in $C_{bound}$ since the enzyme has just returned from I. Therefore, the measured transition rate $k_{C \to I}$ corresponds to the inverse of the waiting time for reaching state I starting in $C_{bound}$, after possible reversible transitions to $C_{free}$ via dissociation and reassociation. It can be shown that this gives the hyperbolic relationship:

$$k_{C\to I} = \frac{k_{open}}{K_{d,init} + [RNP]} \times [RNP]$$

Note the similarity of this expression to the Michaelis-Menten-like relationship for the overall rate of reaching I from $C_{free}$ :

$$k_{Cfree\to I} = \frac{k_{open}}{K_{1/2} + [RNP]} \times [RNP]$$

where $K_{1/2} = \frac{k_{off,init}+k_{open}}{k_{on,init}}$ . The two expressions above are equivalent for $k_{off,init} >> k_{open}$.

For dSpRY, the parameters $K_{d,init}$(dSpRY) and $k_{open}$(dSpRY) are obtained directly from the fit to the hyperbolic saturation curve. At low [RNP], the modeled relationship between $k_{C\to I}$ and [RNP] is approximately linear with slope $k_{on,eff}(dSpRY) = \frac{k_{open}(dSpRY)}{K_{d,init}(dSpRY)}$ = 0.004 $nM^{-1}s^{-1}$ while at high [RNP], $k_{C\to I}$ plateaus and approaches $k_{open}$(dSpRY) = 0.06 $s^{-1}$.

For dCas9, we measure a linear relationship without reaching saturation, and therefore the fit does not separately determine $K_{d,init}$(dCas9) and $k_{open}$(dCas9) but only the ratio $k_{on,eff}(dCas9) = \frac{k_{open}(dCas9)}{K_{d,init}(dCas9)}$ = 0.05 $nM^{-1}s^{-1}$. Since our measurements remain within the linear regime, we can directly constrain $K_{d,init}$(dCas9) >> 80 nM, the highest [dCas9] sampled. Similarly, we can directly constrain $k_{open}$(dCas9) >> 4 $s^{-1}$ , the highest $k_{C\to I}$ measured. Based on prior literature[1,4,5,7], we propose a higher lower bound $K_{d,init}$(dCas9) > 1 μM, which implies $k_{open}$(dCas9) > 50 $s^{-1}$ due to our measurement of $\frac{k_{open}}{K_{d,init}}$.

In this approximate treatment, the DNA segment in the vicinity of the target is treated as a site that can be unoccupied ($C_{free}$), or occupied by an RNP in one of three states ($C_{bound}$ , I, or O). Short-range 1D sliding[8] is not treated explicitly, and the $C_{bound}$ state may be understood as an ensemble of configurations where the enzyme may be in different registries and engagement modes. The free [RNP] is assumed to be equal to the concentration injected into the chamber, given the large excess of RNP over DNA in the single-molecule experiment, and [RNP] is not corrected for specific activity, which does not affect the relative comparison between dSpRY and dCas9 since their measured active fractions are equivalent (Fig. S1H). Interactions between enzymes on the DNA, which could occur at high [RNP] for dSpRY[7] are also not considered in this simple model.

**Note S2. Analysis of ChIP-seq of dCas9 and dSpRY in human cells**

ChIP-seq analysis of dCas9 and dSpRY revealed distinct binding profiles. dCas9 showed strong enrichment at the target site (Fig. S6F-G) and higher reads per kilobase per million mapped reads (RPKM) to its specific seed and PAM sequence (5’-AAGAANGG-3’) (Fig. S6E-G). This suggests dCas9's high specificity for PAM-mediated binding.

In contrast, dSpRY did not show enrichment at the target site (Fig. S6F-G), instead displaying uniform RPKM across regions with and without its corresponding seed and PAM motifs (5’-AAGAANRN-3’) (Fig. S6F-G). This pattern suggests a non-specific binding behavior, likely due to dSpRY’s broader PAM recognition. Although the uniform coverage might partly result from background noise or lower RNP levels due to reduced sgRNA expression (Fig. S6F-G), it highlights dSpRY’s less specific targeting compared to dCas9.
